# A genome-wide cytotoxicity screen of Cluster F1 mycobacteriophage Girr reveals novel inhibitors of *Mycobacterium smegmatis* growth

**DOI:** 10.1101/2023.08.04.552056

**Authors:** Richard S Pollenz, Kaylee Barnhill, Abbigail Biggs, Jackson Bland, Victoria Carter, Michael Chase, Hayley Clark, Caitlyn Coleman, Marshall Daffner, Caitlyn Deam, Alyssa Finocchiaro, Vanessa Franco, Thomas Fuller, Juan Gallardo Pinera, Mae Horne, Zoe Howard, Olivia Kanahan, Christopher Miklaszewski, Sydney Miller, Ryan Morgan, Oluwatobi Onalaja, Louis Otero, Shivani Padhye, Emily Rainey, Fareed Rasul, Alexandra Rodier, Sydni Schlosser, Ava Sciacchitano, Emma Stewart, Rajvi Thakkar, Danielle Heller

## Abstract

Over the past decade, thousands of bacteriophage genomes have been sequenced and annotated. A striking observation from this work is that known structural features and functions cannot be assigned for >65% of the encoded proteins. One approach to begin experimentally elucidating the function of these uncharacterized gene products is genome-wide screening to identify phage genes that confer phenotypes of interest like inhibition of host growth. This study describes the results of a screen evaluating the effects of overexpressing each gene encoded by the temperate Cluster F1 mycobacteriophage Girr on the growth of the host bacterium *Mycobacterium smegmatis*. Overexpression of 29 of the 102 Girr genes (∼28% of the genome) resulted in mild to severe cytotoxicity. Of the 29 toxic genes described, 12 have no known function (NKF) and are predominately small proteins of <125 amino acids. Overexpression of the majority of these 12 cytotoxic NKF proteins resulted in moderate to severe growth reduction and represent novel antimicrobial products. The remaining 17 toxic genes have predicted functions, encoding products involved in phage structure, DNA replication/modification, DNA binding/gene regulation, or other enzymatic activity. Comparison of this dataset with prior genome-wide cytotoxicity screens of mycobacteriophages Waterfoul and Hammy reveals some common functional themes, though several of the predicted Girr functions associated with cytotoxicity in our report, including genes involved in lysogeny, have not been described previously. This study, completed as part of the HHMI-supported SEA-GENES project, highlights the power of parallel, genome-wide overexpression screens to identify novel interactions between phages and their hosts.

## Introduction

The co-evolution of bacteriophages and bacteria has given rise to important discoveries that have shaped modern molecular biology. These include the identification of restriction-modification systems that have revolutionized the ability to manipulate DNA (reviewed in Loenen *et al*., 2014) and the discovery of the CRISPR-Cas systems that allow precise genetic engineering of numerous model systems (reviewed in Mazhar 2018). Additionally, the study of phages has provided insight into elegant gene regulatory and signaling systems (Ptashne 2004), as well as unexpected phenomena such as the expression of phage tail genes by bacteria to fight off competitors that may be infringing on their resources (Patz *et al*., 2019). Thus, phages contain a rich reservoir of continually evolving genes that have the potential to offer important insights into genomics, antibiotic development, and bacterial host immunity.

The Actinobacteriophage Database (phagesDB.org) contains >4,000 annotated actinobacteriophage genomes (Russell and Hatful 2016). The genomes in this database are highly mosaic, and the predicted protein coding genes are grouped into thousands of unique phamilies (phams) based on amino acid similarity of their protein products (Cresawn *et al*. 2011; Gauthier *et al*. 2022). The majority of these phams have no recognizable sequence features and cannot be assigned functions via homology to characterized sequences (Pope *et al*. 2015, 2017; Hatfull 2020). One approach to begin to elucidate the function of these gene products is screening phage gene overexpression libraries for phenotypes of interest in the bacterial host. This type of approach has been employed for several different mycobacteriophages using cytotoxicity in the host *Mycobacterium smegmatis* as the phenotypic endpoint (Ko and Hatfull 2020; Heller *et al*. 2022, Amaya *et al*. 2023). For example, Ko and Hatfull evaluated 193 unrelated genes from 13 diverse phages and found 45 genes (23% of the genes tested) that when overexpressed resulted in various levels of *M. smegmatis* growth inhibition. Furthermore, reports of systematic overexpression of each gene from two Cluster K mycobacteriophages Waterfoul and Hammy shows that up to one-third of genes encoded in a single mycobacteriophage genome can disrupt the growth of *Mycobacterium smegmatis* to varying degrees (Heller *et al*. 2022, Amaya *et al*. 2023).

The current study, which was completed as part of the HHMI-supported SEA-GENES project (Heller and Sivanathan 2022), expands on the data from the studies above and presents a genome-wide overexpression screen of Girr. Girr is a temperate Cluster F1 siphovirus that infects *M. smegmatis* and encodes a single tRNA (Trp) and 102 predicted protein coding genes, of which only 41 can be assigned a predicted function. Consistent with previous reports, overexpression of 29 Girr genes (∼28% of the genome) resulted in some level of cytotoxicity, with overexpression of 11 causing severe growth reduction. The majority of these cytotoxic genes encode small (<125 amino acids) proteins of no known function (NKF), and all but 3 are distinct from those identified in other screens (Ko and Hatfull 2020; Heller *et al*. 2022, Amaya *et al*. 2023), thus, presenting many novel entries to the set of known anti-mycobacterial phage products.

## Materials and Methods

### Growth of mycobacteria and mycobacteriophage

*Mycobacterium smegmatis* mc^2^155 was grown at 37 °C in Middlebrook 7H9 (Difco) broth supplemented with 10% AD (2% w/v Dextrose, 145 mM NaCl, 5% w/v Albumin Fraction V), 0.05% Tween80, and 10 μg/ml cycloheximide (CHX) or on Middlebrook 7H10 or 7H11 (Difco) agar supplemented with 10% AD, 0.5% glycerol and 10 μg/ml CHX. For transformation of *M. smegmatis* mc^2^155, electrocompetent cells were electroporated with ∼100 ng of pExTra plasmid DNA and recovered in 1 ml of 7H9 broth for 2 h at 37 °C with shaking (225 rpm); transformants were selected on 7H10 or 7H11 agar supplemented with 10 μg/ml Kanamycin monosulfate (Fisher Bioscience). Colonies were harvested for the cytotoxicity assay after 4-5 days of selection at 37 °C. Phage Girr was propagated on *M. smegmatis* mc^2^155 lawns grown at 37 °C on 7H10 agar and 7H9 top agar supplemented with 1 mM CaCl_2_.

### Construction of the pExTra Girr Library

pExTra01 is an anhydrotetracycline (aTc) inducible expression shuttle vector with promoter *pTet* directly upstream of the gene insertion site and an *mcherry* transcriptional reporter gene directly downstream of the gene insertion site (Figure 1A; Heller *et al*. 2022). Girr genes were PCR-amplified using gene-specific primers (Integrated DNA Technologies), Q5 DNA polymerase (New England Biolabs Q5 HotStart 2X Master Mix), and a high-titer Girr lysate. Gene-specific primers (Supplemental Table 1) anneal to the first and last 15-25 bp of each gene sequence and introduce a uniform ATG start codon (forward primers) or TGA stop codon (reverse primers). Each primer was also designed to contain regions of identity to the pExTra01 plasmid; forward primers contain a uniform RBS and 5’ 21 bp of homology to pExTra01 downstream of the *pTet* promoter, and all reverse primers contain a separate 5’ 25 bp of homology to pExTra01 upstream of *mcherry* (Supplemental Table 1). These regions of homology allow the Girr gene sequence to be inserted into pExTra downstream of the *pTet* promoter by isothermal assembly (NEB HiFi 2X Master Mix) as previously described (Heller *et al*. 2022). Linearized pExTra01 plasmid was prepared for isothermal assembly reactions via PCR (NEB Q5 HotStart 2X Master Mix) of pExTra01 using divergent primers pExTra_F and pExTra_R. Girr *57* could not be amplified by PCR and was synthesized as a dsDNA gblock fragment with the flanking homology regions (IDT). Recombinant plasmids were recovered by transformation of *Escherichia coli NEB5α F’I*^*Q*^ (New England Biolabs) and selected on LB agar supplemented with 50 μg/ml Kanamycin. All pExTra-Girr plasmids were sequence-verified by Sanger sequencing (Genewiz/Azenta) using sequencing primers pExTra_universalR and pExTra_seqF; longer genes were also sequenced with internal sequencing primers listed in Supplemental Table 1. All plasmid inserts were found to match the published genome sequence (sequence data is available on genesDB.org).

**Table 1:** Girr genes observed to inhibit *Mycobacterium smegmatis* growth upon overexpression.

| GIRR GENE | PHAMILY<sup>1</sup> | LENGTH (AA) | PREDICTED FUNCTION AND FEATURES | CYTOTOXICITY<sup>2</sup> | CYTOTOXIC HOMOLOGS IN OTHER PHAGES |
| --- | --- | --- | --- | --- | --- |
| 6 | 604 | 273 | Major capsid protein | 1 |  |
| 31 | 78171 | 415 | Lysin A | 1 |  |
| 56 | 399 | 161 | HTH DNA binding domain protein | 1 |  |
| 62 | 1005 | 192 | ssDNA binding protein | 1 | Fruitloop gp60<sup>4</sup> |
| 63 | 78110 | 120 | HNH endonuclease (homing) | 1 |  |
| 64 | 78432 | 839 | DNA helicase/methylase | 1 |  |
| 69 | 78275 | 49 | NKF | 1 |  |
| 102 | 76667 | 207 | Glycosyltransferase | 1 |  |
| 5 | 649 | 197 | Scaffolding protein | 2 |  |
| 19 | 293 | 628 | Minor tail protein | 2 |  |
| 22 | 364 | 84 | NKF | 2 |  |
| 32 | 77567 | 250 | HNH endonuclease (restriction) | 2 |  |
| 37 | 70 | 273 | DnaQ-like exonuclease | 2 |  |
| 44 | 78168 | 433 | Integrase | 2 |  |
| 52 | 78264 | 215 | NKF | 2 |  |
| 54 | 78220 | 115 | NKF; TMD (1)<sup>3</sup> | 2 |  |
| 65 | 77148 | 66 | NKF | 2 |  |
| 72 | 1656 | 74 | NKF | 2 |  |
| 100 | 78227 | 487 | Glycosyltransferase | 2 |  |
| 2 | 78090 | 545 | Terminase, large subunit | 3 |  |
| 35 | 807151 | 124 | NKF; TMD (1) | 3 | Waterfoul gp32<sup>5</sup> |
| 36 | 333 | 86 | NKF | 3 |  |
| 46 | 78503 | 190 | Immunity repressor | 3 |  |
| 48 | 78241 | 334 | Antirepressor | 3 |  |
| 51 | 67076 | 63 | NKF | 3 |  |
| 57 | 80778 | 92 | WhiB family transcription factor | 3 | Hammy gp53<sup>6</sup>; TM4 gp49<sup>7</sup> |
| 60 | 328 | 76 | NKF | 3 |  |
| 71 | 1432 | 119 | NKF | 3 |  |
| 73 | 4981 | 107 | NKF | 3 |  |

<sup>1</sup>Phamily numbers were recorded from phagesdb.org as of May 17, 2023.
<sup>2</sup> Growth of strains on media supplemented with 10 ng/ml or 100 ng/ml aTc was compared to the same strains plated on media without aTc. Those scored as 0, had no effect on host growth at aTc-10; those scored as 1 exhibit an aTc-dependent reduction in colony size, those denoted as 2 demonstrate a ~1-3-log difference in growth in the presence of aTc, and those denoted as 3 demonstrate a severe >3-log reduction in growth in the presence of aTc.
<sup>3</sup> TMD= Transmembrane domain and number of predicted helices in parentheses.
<sup>4</sup> Ref: Ko and Hatfull 2020
<sup>5</sup> Ref: Heller *et al.* 2022
<sup>6</sup> Ref: Amaya *et al.* 2023
<sup>7</sup> Ref: Rybniker *et al.* 2010

**Figure 1:**
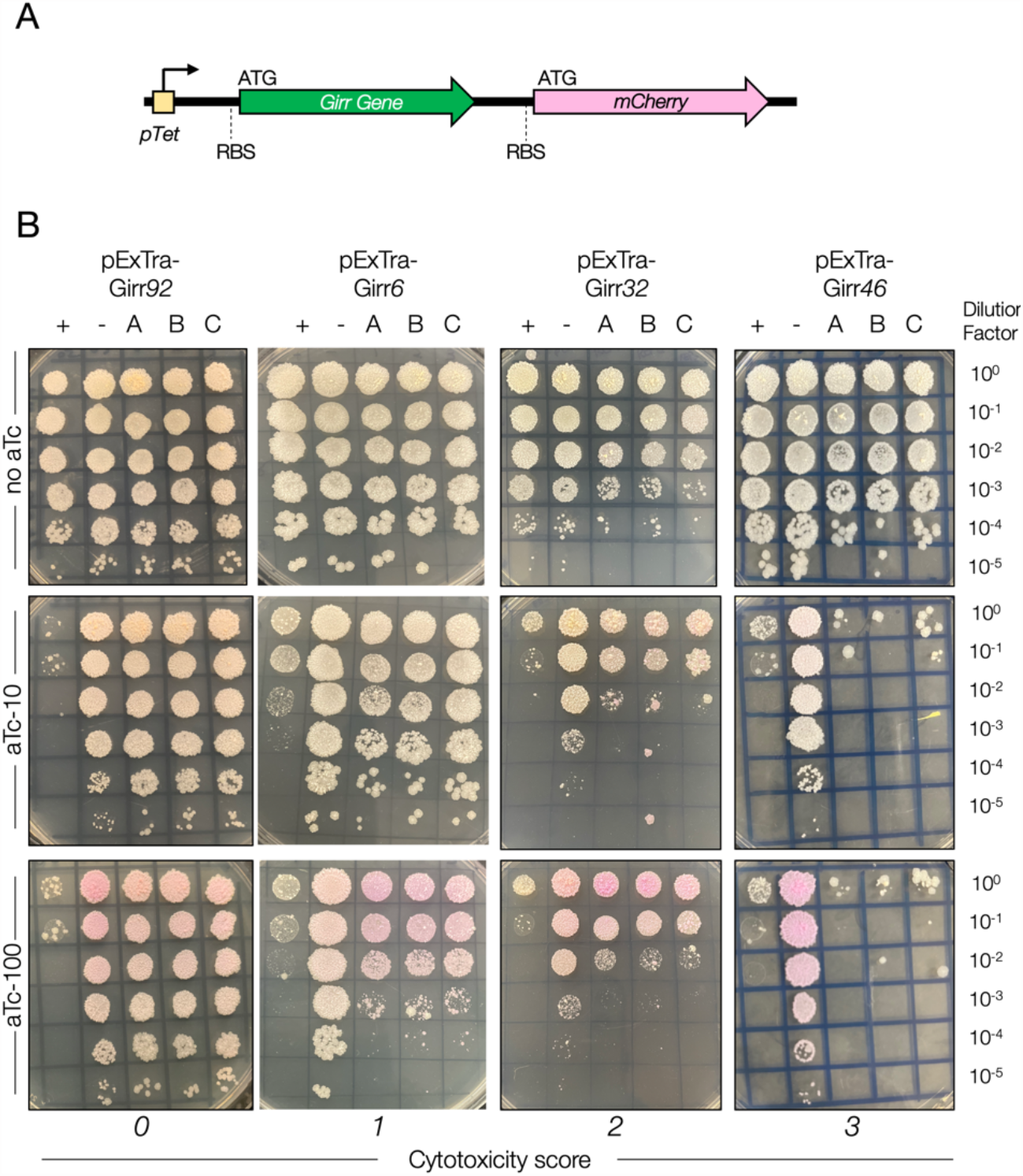
Results of cytotoxicity assays for representative Girr genes. A) pExTra expression plasmids contain the Girr gene downstream of the *pTet* promoter and upstream of *mcherry*. The two genes in this aTc-inducible operon are transcriptionally linked, with each gene having distinct start and stop codons and ribosomal binding sites (RBS) for translation of the two protein products. B) In each assay, colonies of *M. smegmatis* mc^2^155 transformed with the specified pExTra plasmid were resuspended, serially diluted, and spotted on 7H10 Kan media containing 0, 10, or 100 ng/ml aTc. Triplicate colonies (labeled A, B, C) were tested for each Girr gene alongside a positive control strain (+) transformed with pExTra02 (expressing wildtype Fruitloop *52*) and a negative control strain (-) transformed with pExTra03 (expressing Fruitloop *52 I70S*). Cytotoxicity score (0) is represented by Girr *92*. Cytotoxicity score (1) is represented by Girr *6*. Cytotoxicity score (2) is represented by Girr *32*. Cytotoxicity score (3) is represented by Girr *46*.

### Cytotoxicity Screening and Phenotype Scoring

Three colonies of *M. smegmatis* mc^2^155 transformed with each pExTra-Girr gene plasmid were harvested and suspended in 450 μl of 7H9 media by vortexing. Each colony suspension was serially diluted in 7H9 broth and dilution series spotted on 7H10 agar plates supplemented with 10 μg/ml Kanamycin and 0, 10, or 100 ng/ml anhydrotetracycline (aTc; Alfa Aesar). The sample was vigorously shaken before each spot was added to the plate. Two controls were also spotted on each plate to aid in evaluation of gene-mediated growth impacts. The pExTra02 positive control plasmid encodes cytotoxic gene Fruitloop *52*, and pExTra03 served as the negative control plasmid and encodes a non-toxic mutant allele of Fruitloop *52* with an isoleucine to serine substitution at amino acid position 70 (Ko and Hatfull 2018; Heller *et al*. 2022). Growth was monitored over 4-5 days at 37 °C. Cytotoxic phenotypes were scored by comparing the spot dilution series for Girr genes on the control plates lacking aTc to the plates supplemented with aTc inducer as well as by comparing the level of growth of the Girr gene triplicates compared to cells containing the pExTra03 negative control plasmid in the presence of aTc. Cytotoxic phenotypes were classified as either having no effect (score 0), being moderately cytotoxic with a 1-3 log reduction in cell viability (score 2), or being severely cytotoxic, causing complete or near complete (>3-log) inhibition of growth (score 3). Strains were also evaluated for aTc-dependent size reduction in individual colonies (score 1) as compared to the Fruitloop *52-I70S* negative control strain on the same aTc plate and the same strain on plates without inducer. In most cases where genes were scored as 0, the colonies turned pink in the presence of Inducer, indicating aTc-dependent *mcherry* gene expression and providing a visual indicator of expression through the the *pTet* operon.

All genes with a cytotoxic phenotype were evaluated in at least two independent experiments. In general, most experiments showed a high level of consistency of growth/no growth across the three Girr gene replicates (Supplemental Figure 1). In some instances, all spotted cells, including cells with the pExTra03 negative control plasmid showed a slight growth reduction in the presence of aTc (see examples Girr *2, 16, 25* in Supplemental Figure 1). In these cases, cytotoxicity was only scored if there was additional notable reduction in growth of Girr strains compared to the negative control cells on the aTc plates. In addition, for some genes, slight variation in the magnitude of cytotoxicity was observed between independent experiments, especially for those scored as 1. In these instances, the genes were scored based on the mildest result. The inclusion of the Fruitloop *52* control strains on each screening plate aided in the evaluation of relative gene-mediated effects, and the cytotoxicity results are scored based on observations on 100 ng/ml aTc plates.

### Girr genomic analysis

The Girr genome map was created using the web-based tool Phamerator (phamerator.org; Cresawn *et al*. 2011). Reported gene functions were based on those available in the Girr GenBank record (Accession MH669003). All functional calls were confirmed using HHPRED (PDB_mmCIF70_10_Jan, SCOPe70_2.08, Pfam-A_v35, NCBI_Conserved_Domains(CD)_v3.19) (Gabler *et al*. 2020). Functional calls were updated based on newly added PDB entries and by comparison of functional calls for other pham members on PhagesDB. This included updating the functions of gp44 (tyrosine integrase), gp56 (HTH DNA binding domain protein) and gp62 (ssDNA binding protein). The number and location of predicted transmembrane helices (TMD) were identified within query protein sequences utilizing three different programs: SOSUI (Hirokawa *et al*., 1998), Deep TMHMM (Hallgren *et al*., 2022) and TOPCONS (Tsirigos *et al*., 2015). TMD proteins were characterized based on agreement of all three programs as detailed in Pollenz et al. (2022). 3D modeling of Girr gp46 was performed using Phyre^2^ (Kelly *et al*., 2015). The identity between gene products was determined by multiple sequence alignment with Clustal Omega (Madeira *et al*. 2022). Gene phamily designations used for phenotypic comparisons of related genes were obtained from phagesDB.org on May 17, 2023.

## Results and Discussion

### Study overview

An arrayed library of all 102 Girr protein coding genes were cloned into the pExtra expression plasmid under the control of the inducible *pTet* promoter and linked to an *mcherry* transcriptional reporter (Figure 1A). All pExTra-Girr gene plasmids were sequence-verified and used to transform *M. smegmatis* for evaluation in a semi-quantitative, plate-based cytotoxicity assay. Colonies of transformed strains were suspended, serially diluted, and spotted in triplicate on media containing increasing concentrations of the inducer aTc alongside control strains expressing wildtype cytotoxic gene Fruitloop *52* or a non-toxic mutant allele (Ko and Hatfull 2018). The impact of Girr gene overexpression on cell growth was scored as: (0) no impact on bacterial growth, (1) reduction in colony size, (2) a 1-3-log reduction in bacterial growth in the presence of aTc, or (3) near complete abolition of bacterial growth in the presence of aTc. Figure 1B shows a representative example of the different levels of toxicity: overexpression of Girr *6* results in mild colony size reduction across the three replicates (score 1), Girr *32* demonstrates moderate cytotoxicity (score 2), and Girr *46* is representative of severe cytotoxicity, with very little bacterial growth observed across the three replicate samples (score 3). In all examples, it can be observed that there is a significant aTc-dependent reduction in the growth of cells expressing the Fruitloop *52* positive control. Tunable induction of *pTet* by aTc is also confirmed by the observation of an aTc-dependent increase in pink colony color due to production of mCherry from the transcriptionally linked reporter gene (Figure 1A; Ehrt *et al*. 2005; Heller *et al*. 2022). Interestingly, pink color was observed for four Girr genes (*36, 44*, 81, and *99*) even in the absence of inducer, suggesting that these sequences may harbor internal promoter sequences (Supplemental Figure 1). The corresponding data and toxicity scores for all 102 genes is presented in Supplemental Figure 1.

### Overexpression screen reveals 29 cytotoxic Girr genes

Overall, overexpression of 29 Girr genes in *M. smegmatis* caused a reduction in bacterial growth, with expression of 21 of the genes leading to moderate to severe toxicity (Figure 2, Table 1). The observation that 28% of the Girr genome inhibits *M. smegmati*s growth is in line with the frequencies observed in the genome-wide screens of phages Waterfoul (34% of the genome was cytotoxic; Heller *et al*. 2022) and Hammy (25% of the genome was cytotoxic; Amaya *et al*. 2023). For almost all the genes that had no measurable impact on growth (69 out of 73), *mcherry* expression was visible, confirming expression through the *pTet* operon (Supplemental Figure 1). However, in our screen, protein levels of Girr gene products were not determined, and we cannot rule out that some gene products classified as non-toxic may be poorly produced in our system.

**Figure 2:**
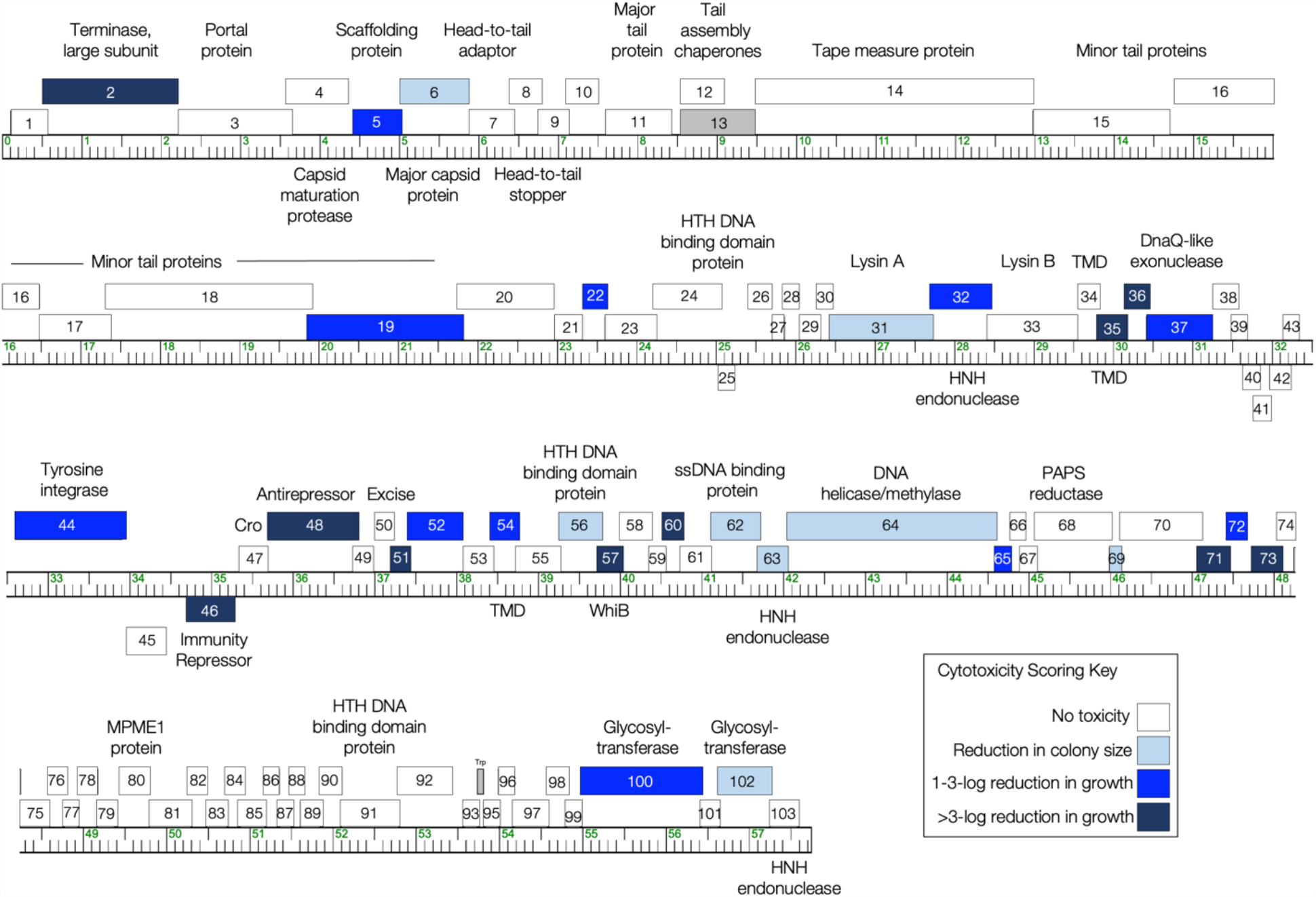
The genome of phage Girr. The Girr genome is shown as a ruler with kbp markers and genes represented by boxes—those above the line are transcribed rightwards and those below are transcribed leftwards. Numbers inside the box correspond to gene numbers and predicted functions are indicated above or below each gene. Box shading corresponds to cytotoxicity scoring, with white boxes designating genes found to have no effect on *M. smegmatis* growth (cytotoxicity score *0*), and the gray box for gene *13* reflecting the ORF for the longer tail assembly chaperone that requires a translational frameshift from the open reading fame of gene *12* to encode the larger protein that cannot be verified to have been synthesized and was exluded from the analysis. The shade of blue corresponds to the severity of growth inhibition using the following scores (a key is also provided in the figure): light blue (score *1*; reduction in colony size; genes *6, 31, 56, 62, 63, 64, 69*, and *102*), medium blue (score *2*; 1–3 log reduction in viability; genes *5, 19, 22, 32, 37, 44, 52, 54, 65, 72*, and *100*), and dark blue (score *3*; >3-log reduction in viability; genes *2, 35, 36, 46, 48, 51, 57, 60, 71*, and *73*).

Amongst the Girr genes identified as cytotoxic, 17 have predicted functions. Three are classified as structural gene products (scaffolding protein gp5, major capsid protein gp6, and minor tail protein gp19), five are involved in DNA replication/modification (DNA helicase/methylase gp64, HNH endonucleases gp63 and gp32, DnaQ gp37, ssDNA binding protein gp62), and four have some predicted enzymatic functions (terminase large subunit gp2, glycosyltransferases gp100 and gp102, and Lysin A protein gp31). Although Lysin A typically works in concert with Lysin B and holin proteins to mediate host lysis, there have been prior reports of some mycobacteriophage Lysin A proteins causing toxicity when individually overproduced (Payne and Hatfull 2012, Amaya et al 2023). Additionally, five cytotoxic Girr gene products are involved in DNA binding/regulation, including putative helix-turn-helix (HTH) DNA-binding protein, gp56, and WhiB family transcription factor gp57; this also includes several genes predicted to be necessary for the lysogenic lifestyle of temperate phage Girr (Ptashne 2004). Overexpression of the genes encoding the tyrosine integrase (gp44) involved in the integration of the phage genome into the bacterial genome, the immunity repressor (gp46) involved in turning off lytic genes during lysogeny, and a putative antirepressor (gp48) that may be involved in gene regulation during the lytic lifestyle, resulted in moderate to severe host growth reduction (Table 1 and supplemental Figure 1). Since these proteins can be classified as DNA binding, the observed toxicity upon overexpression may be a result of the disruption to normal host gene regulation. However, in temperate phages the immunity repressor is typically continually expressed in a lysogen to repress lytic gene expression (Ptashne 2004), and the near complete reduction in cell growth from overproduction of Girr gp46 is surprising.

The prototypical immunity repressor, CI from coliphage lambda, binds DNA as a homodimer and contains an N-terminal DNA binding domain with two essential α-helices and a C-terminal domain involved in homo-dimerization and cooperative binding (Ptashne 2004). Girr gp46 is 190 amino acids and has many hits with >96% probability to transcriptional repressors predicted by HHpred, including the CI repressor proteins from *E. coli* phage lambda (PDB 3KZ3_A; Lui *et al*., 2010) and *E. coli* phage 186 (2FJR_B; Pinkett et al., 2006). Phyre^2^ was utilized to model the N-terminal 74 amino acids of Girr gp46 on a structure of the lambda repressor (d1rioa; PDB 3KZ3_A) (Kelly *et al*., 2015). The 3D modeling predicted two N-terminal α-helices in Girr gp46 between amino acids 27-34 and 46-54 that precisely align to the DNA binding α-helices of lambda CI (Figure 3A, Ptashne 2004). The C-terminal portion of Girr gp46 has no recognizable features nor homology to characterized immunity repressors, and it is unknown whether it plays a similar role in dimerization. To assess if the predicted DNA binding domain is necessary for observed cytotoxicity, a Girr *46* truncation lacking helix 2 (Girr*46Δ34*) was cloned into the pExTra overexpression vector. As shown in Figure 3B, expression of Girr*46Δ34* resulted in severe toxicity, comparable to that observed with expression of full length Girr *46*, suggesting that an intact DNA-binding domain is not necessary for the host growth inhibition observed in our assay.

**Figure 3:**
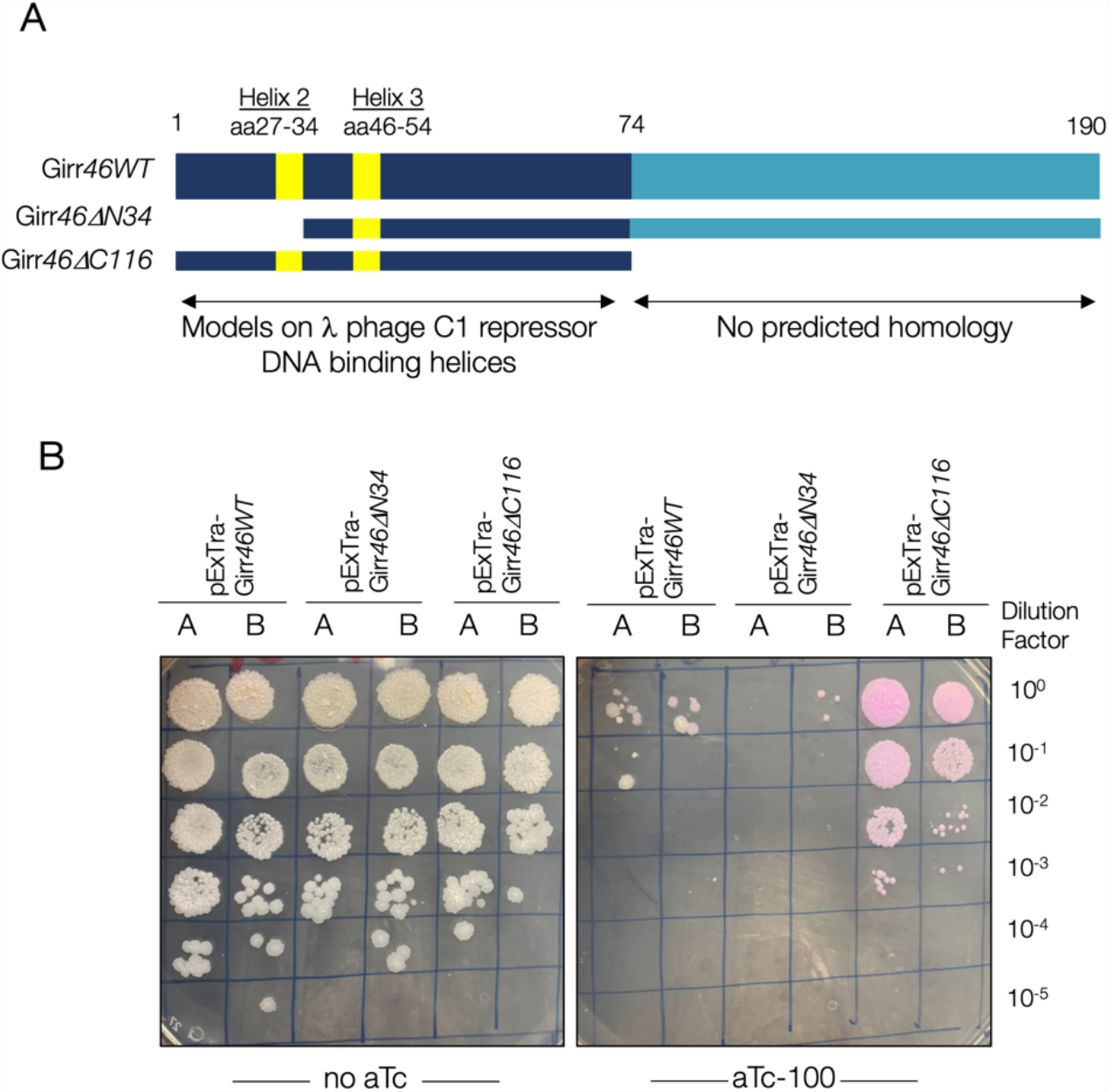
Truncation analysis of the Girr immunity repressor gp46. A) Schematic representation of Girr gp46 and the tested truncations. Girr gp46 is 190 amino acids and the 74 amino acid N-terminal DNA binding region and predicted alpha-helices (yellow) are shown. Girr*46ΔN34* removes the first 34 amino acids and helix 2. Girr*46ΔC116* removes the C-terminal 116 amino acids and includes only the N-terminal 74 amino acids. B) In each assay, colonies of *M. smegmatis* mc^2^155 transformed with the specified Girr *46* pExTra construct were resuspended, serially diluted, and spotted on 7H10 Kan media containing 0 or 100 ng/ml aTc. Duplicate colonies (labeled A and B) were tested for each Girr *46* gene construct.

Overexpression of a truncation lacking the last 116 residues of gp46 (Girr*46Δ116*) showed similar growth with or without induction and aTc-dependent *mcherry* expression, suggesting that the last 116 amino acids of gp46 are required for host interference. Although expression of the Girr*46Δ116* allele from pExTra did result in pink colony color, we cannot rule out that the Δ116 variant may be misfolded or may not stably accumulate in our system. Cytotoxicity was not previously reported for overexpression of the immunity repressors encoded by Waterfoul (gp42) or Hammy (gp45), though these proteins have low sequence similarity to Girr gp46 (Heller *et al*. 2022; Amaya *et al*. in review). Currently there are >1,700 predicted immunity repressors grouped to 50 different phamilies in the Actinobacteriophage Database, and it will be interesting to explore how widespread host growth inhibition by overexpression of immunity repressors is amongst temperate actinobacteriophages.

The remaining 12 cytotoxic genes are of unknown function (NKF) and are distinct from any of the toxic NKF genes identified in any of the other cytotoxicity screens reported to date (Ko and Hatfull 2020; Heller et al., 2022; Amaya et al., in review). In general, the protein products of the Girr NKF genes are small, with 11 of the genes encoding proteins of <87 amino acids; only Girr gp52 is >200 amino acids (Table 1). The majority of these NKF genes are moderately to severely toxic with all but one scored as 2 or 3 on the toxicity scale (Table 1). The protein products have no conserved domains except for gp35 and gp54 which each contain a single transmembrane domain (TMD).

In all, a diverse set of Girr genes were observed to inhibit growth of the host *M. smegmatis* upon overexpression. While some of the effects reported here may be due to artificially high protein levels or off-target effects, the toxicity mediated by many may be directly related to their physiological role in the phage life cycle, making them intriguing targets for future analysis. These results also underscore the complex nature of host interactions that can be mediated by a single phage genome.

### Conservation of cytotoxicity for gene phamilies encoded by Girr

We next utilized the Actinobacteriphage Database (PhagesDB.org) to review each gene phamily encoded by Girr and determine if they contained known toxic genes from any of the other mycobacteriophages that have been previously evaluated. Table 1 shows that three Girr growth inhibitors reported in this study are members of phams that were previously identified as cytotoxic: Girr gp35 is homologous to Waterfoul gp32, Girr gp57 (WhiB family transcription factor) is a homolog to cytotoxic WhiB proteins Hammy gp53 and TM4 gp49, and Girr gp62 (ssDNA binding protein) is a homolog of cytotoxic Fruitloop gp60 identified by Ko and Hatfull (Heller et al. 2022, Amaya et al. 2023, Ko and Hatfull 2020, Rybniker et al. 2010). Thus, the current screen has identified a diverse set of 26 novel phams that can impact host growth upon overexpression.

Ko and Hatfull previously evaluated 36 genes from Cluster F1 mycobacteriophage Fruitloop for impacts on *M. smegmatis* growth in a different overexpression system; Girr encodes homologous pham members for 14 of these (Ko and Hatfull 2018, 2020). For 10 of these shared phams, the data presented here align with the prior Fruitloop data: this includes the pham represented by cytotoxic gene products Fruitloop gp60 and Girr gp62, as well as 9 phams that were reported as non-toxic in both studies (phams represented by Girr gp26, gp74, gp75, gp80, gp85, gp86, gp93, gp97, gp101). There were 4 instances where one pham member was found to be toxic but the other was not (phams represented by Girr gp52, gp53, gp56, and gp95). For example, Fruitloop gp52 encodes a 92 amino acid protein that is not essential for Fruitloop growth, but results in a moderate to severe cytotoxic phenotype when expressed in *M. smegmatis* (Ko and Hatfull 2018, see pExTra02 positive controls on all assays in Supplemental Figure 1). Ko and Hatfull reported that Fruitloop gp52 cytotoxicity is due to disruption of the essential *M. smegmatis* cell wall biogenesis protein Wag31 and proposed that gp52 interacts with this host factor to block superinfection by heterotypic phages (Ko and Hatfull 2018). Although Girr gp53 shares 85% amino acid identity with Fruitloop gp52, in the current study, overproduction of Girr gp53 caused no reduction in *M. smegmatis* growth (Figure 4A and Supplemental Figure 1). *M. smegmatis* cells transformed with pExTra-Girr*53* turned slightly pink on the higher level of aTc, validating that the *pTet* operon was expressed and mCherry protein is produced (Figure 4A). Detailed analysis of the amino acid alignment of Girr gp53 to Fruitloop gp52 shows that there are 13 amino acid substitutions and 1 insertion between the two sequences; nine of the amino acid substitutions occur in the N-terminal 29 amino acids, a region previously shown to not be essential for gp52-mediated toxicity (Ko and Hatfull 2018). However, Girr gp53 has a non-conserved H/V substitution at amino acid 62 and an I/V substitution at amino acid 70, which is part of a helix predicted to be involved in Wag31 interaction, and indeed is the same amino acid changed in the Fruitloop gp52I/S non-toxic mutant that is used as the negative control in the current studies (Ko and Hatfull 2018). Isoleucine is found at this position for most homologous phamily members, though valine is found in 29 of the 213 members (<14%) of this gene phamily. All members of this pham also contain a C-terminal D/EAA ssrA-like tag associated with protein degradation (Figure 4B; Ko and Hatfull 2018). Interestingly, Girr has a glycine residue, not present in other pham members, inserted just prior to the EAA domain at amino acid 91 (Figure 4B), and lack of cytotoxicity upon overproduction of Girr gp53 may be because it is not produced to the same level as Fruitloop gp52. Further work is needed to determine how these sequence variations contribute to the observed phenotypic divergence within this pham. Importantly, these findings show the utility of evaluating multiple proteins within a given pham to help identify the sequence determinants of phage-mediated phenotypes.

**Figure 4:**
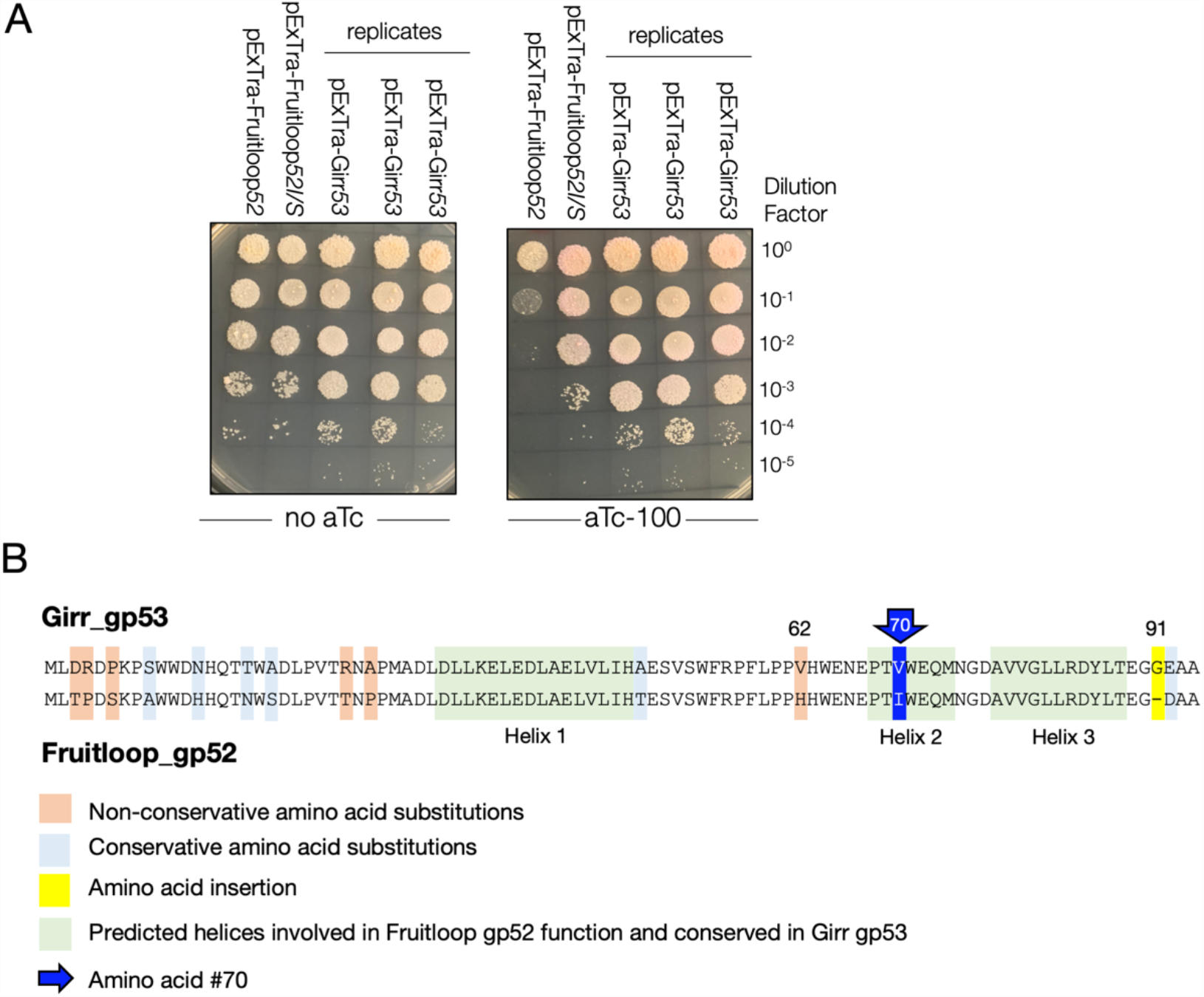
Cytotoxicity and amino acid alignment of Girr gp53 and Fruitloop gp52. A) In each assay, colonies of *M. smegmatis* mc^2^155 transformed with the specified pExTra plasmid were resuspended, serially diluted, and spotted on 7H10 Kan media containing 0 or 100 ng/ml aTc. Triplicate colonies were tested for Girr *53* alongside Fruitloop *52* and the Fruitloop *52 I70S* mutant that has reduced toxicity (Ko and Hatfull 2018). B) The amino acid alignment of Girr gp53 and Fruitloop gp52. The predicted helices involved in Fruitloop gp52 function that are conserved in Girr 53 are highlighted in green. The 13 amino acid substitutions are highlighted with blue (conservative substitutions), or orange (non-conservative substitutions). The G amino acid substitution at 91 is highlighted in yellow and the I/V substitution at amino acid 70 is highlighted in blue.

### Patterns of phenotypic conservation across mycobacteriophage genomes

The list of toxic Girr genes was also reviewed by function to identify commonality to toxic Waterfoul and Hammy genes previously discovered through similar systematic pExTra screening methods. This was important since protein products can be annotated with the same function yet be placed in a different phams because of low amino acid identity (Cresawn *et al*., 2011; Gauthier *et al*., 2022). Several themes regarding the classification of the toxic genes emerged after this analysis. First, in all three screens, there were multiple structural genes from each phage that were toxic. Overproduction of the major capsid protein from Girr (gp6) as well as two members of a different major capsid protein pham, Waterfoul (gp11) and Hammy (gp12), caused similar growth defects in the host. Additionally, two toxic minor tail proteins were identified in Girr (gp19) and Waterfoul (gp21). We note that whereas the tape measure protein from both Hammy (gp20) and Waterfoul (gp19) was highly toxic, the tape measure protein from Girr (gp14) was not. The tape measure proteins from Waterfoul and Hammy are grouped in the same pham, share 58% amino acid identity, and contain several transmembrane domains (TMD); however, these proteins share <23% amino acid identity to the Girr tape measure protein which also lacks any predicted TMDs. Thus, although these proteins likely perform the same function in tailed phages, the results of the three different screens show that toxicity cannot be generalized to functional calls and requires analysis of each protein product. It was surprising to find that numerous structural genes that do not have known enzymatic activity were toxic. However, recent studies demonstrate that expression of phage structural components can trigger diverse bacterial responses that may impact growth (Zhang et al., 2022; Stoker-Avihail et al., 2023).

The second theme was the observation that several toxic genes in each phage encode proteins involved in DNA replication or modification. Finding these types of toxic genes is not unexpected since overexpression of enzymes that cut and modify DNA could feasibly impact normal cellular physiology. Furthermore, polymerases and single strand binding proteins have also been recently shown to interact with bacterial anti-phage defense systems (Stoker-Avihail et al., 2023). In Girr, the DNA helicase/methylase (gp64) and DnaQ-like exonuclease (gp37) are toxic. In Waterfoul, DNA primase/polymerase (gp65) and DNA helicase (gp65) are toxic, while in Hammy, the DnaQ-like DNA polymerase III subunit (gp58), DNA primase/polymerase (gp68) and RusA-like resolvase (gp69) are toxic. Additionally, Girr, Waterfoul and Hammy all have HNH endonucleases that are toxic. In Girr, expression of HNH endonuclease genes *63* and *32* results in mild and moderate growth inhibition, respectively. HNH endonuclease genes in Waterfoul (*62* and *63)* and Hammy (*67*, in the same pham as Waterfoul *63*), also confer mild toxicity. Interestingly, each of the three phages have an additional non-toxic HNH endonuclease that is found as the 3’ terminal gene in the linear phage map: Girr *103*, Waterfoul *95* and Hammy *95* (Heller *et al*. 2022; Amaya *et al*. 2023). HNH endonucleases (also termed His-Me finger) define a diverse family of nucleases that share limited amino acid identity and include nicking endonucleases, homing endonucleases, and restriction endonucleases (Mehta *et al*., 2004). Thus, the three annotated HNH endonucleases from Girr may have distinct functional activities that impact the level of toxicity in the current overexpression assay.

Finally, Waterfoul and Hammy have a lysis cassette gene organization similar to Girr, with a *lysA, lysB* and two downstream genes encoding proteins with TMDs that may act as holins. The LysA proteins from Girr (gp31), Hammy (gp29) and Waterfoul (gp29) are all grouped in different phams and have <35% amino acid identity. It has been shown previously that some endolysins can cause toxicity in *M. smegmatis* when individually overproduced, and expression of Girr gp31, like Hammy gp29 is mildly toxic, whereas Waterfoul gp29 is non-toxic (Heller et al., 2022; Amaya *et al*. 2023). Interestingly, phage Bxz1 gp236 is in the same pham as Girr gp31 but was not found to be toxic to *M. smegmatis* when expressed in a liquid growth assay utilizing a different expression system (Payne and Hatfull 2012). In contrast, overexpression of the downstream genes encoding the single TMD protein (Girr *35*, Waterfoul *32*, and Hammy *32)* significantly inhibits bacterial growth, whereas the overexpression of the annotated holin genes containing 2-4 TMDs (Girr *34*, Waterfoul *31* and Hammy *31)* had no measurable effect on growth. The single TMD proteins encoded by Girr *35*, Waterfoul *32*, and Hammy *32* have an N-out-C-in membrane topology and a highly charged cytoplasmic C-terminal domain that is characteristic of a holin involved in bacterial lysis (Cahill and Young 2019). The high level of toxicity following the expression of these proteins is especially intriguing given the low sequence conservation of the three proteins: although Girr gp35 and Waterfoul gp32 are members of the same pham, they only share 35% amino acid identity, and Hammy gp32 is part of a separate pham with only ∼20% identity to the other two proteins. Although the single TMD proteins are not annotated as holins, a recent report on >80 actinobacteriophages that infect *Gordonia rubripertincta* (Pollenz et al., 2022) shows that the lysis cassette may contain up to four different holin-like genes where the terminal gene typically encodes a protein with a single TMD. A holin-like function of the single TMD proteins is also supported by work from Catalão et al. (2011) on the F1 cluster phage Ms6. Ms6 gp27 is a single TMD protein in the same pham as Girr gp35 and Waterfoul gp32. In these studies, it was demonstrated that production of gp27 was detrimental to *E. coli* growth in liquid and that deletion of both gene *27* and the gene encoding a two TMD protein (gp26) from Ms6 was lethal to phage survival. Thus, the current data provide preliminary evidence that these single TMD products may play a convergent role as holin-like proteins and offer compelling candidates for further functional characterization of the mycobacteriophage lysis pathway.

### Additional insights

This report represents the third genome-wide screen for cytotoxic genes from a mycobacteriophage and the first from a Cluster F1 phage. Girr has <6% gene content overlap to Cluster K5 phage Waterfoul or Cluster K6 phage Hammy which have ∼50% shared gene content (Cresawn eta l., 2011; Russell and Hatfull 2016). Thus, Girr has a very limited number of genes that are grouped to the same phams as Hammy and Waterfoul, providing an opportunity to assess genome-wide patterns of mycobacterial growth inhibition across more distantly related mycobacteriophage genomes. All three screens recovered a similar fraction of toxic genes within a single phage genome (∼25-34% of each genome). While some of the effects reported here may be a consequence of overexpression rather than biologically relevant interaction with the host, especially those scored for minor toxicity (score 1), many of the genes, including 11 NKF genes, caused notable growth defects even on the intermediate aTc concentration, making them especially intriguing targets for future functional analysis.

The cytotoxic genes identified in these three screens are found throughout the genome and include structural, DNA replication/modification, DNA binding, and enzymes, as well as many cytotoxic NKF genes. As seen with Waterfoul and Hammy, we find that toxic Girr genes are often found in close genomic proximity to each other, suggesting they may act in concert with each other throughout the Girr life cycle. Two clusters of growth inhibitory genes, Girr *51-54* and Girr *71-73*, are found downstream of *cro* (gene *47*) and are likely expressed early in the lytic life cycle (Ko and Hatfull 2018), making them good candidates for products involved in the takeover of the host processes. Surprisingly, a cluster of three Girr genes associated with lysogeny were toxic upon overexpression, and it will be interesting to further explore the importance of these phenotypes for the phage and lysogen. Recent work has highlighted several examples of prophage-mediated defense where phages encode products that interfere with host factors to block secondary infection by other phage (Dedrick et al. 2017, Ko and Hatfull 2018, Gentile et al. 2019, Montgomery et al. 2019), and it is possible that some of the cytotoxic genes identified here may play a similar role in phage-phage competition. This finding of clusters of toxic genes also suggests that additional toxic genes may be present in the Girr genome but require the interaction of multiple protein products to observe the toxic phenotype. Thus, the expression of various operonic areas of multiple genes may be necessary to fully evaluate the Girr genes found to be non-toxic in this study.

In summary, the current study adds 26 novel genes, 12 of which have no known function, to the growing database of cytotoxic genes identified from mycobacteriophages. These results highlight the power of the SEA-GENES strategy: performing systematic overexpression screens, in parallel, for a diverse set of phages not only reveals candidates for new phage-host interactions, but also provides opportunities for phenotypic comparisons of related gene products. As shown here, these comparisons can reveal striking differences in toxicity for even highly similar genes, offering opportunities for detailed downstream analyses to understand the determinants of these distinct outcomes. Since to date only a very small fraction of the known mycobacteriophage genes have been experimentally explored (Rybniker *et al*. 2008, 2010, 2011; Mehla *et al*. 2017; Ko and Hatfull 2018, 2020; Heller *et al*. 2022; Amaya *et al*. 2023), the continued genome-wide screening of diverse phage genomes will undoubtedly provide novel insights into protein function, phage biology and evolution, and the identification of possible antimicrobial agents.

## Data Availability

All plasmids and plasmid sequences reported in this study are available upon request. The authors affirm that all data necessary for confirming the conclusions of this article are represented fully within the article and its tables and figures. Extended data can be found at genesDB.org.

## Acknowledgements

This work was conducted as part of the HHMI-supported Science Education Alliance GENES (Gene-function Exploration by a Network of Emerging Scientists) project. We thank members of the Science Education Alliance for their research support including Viknesh Sivanathan, Deborah Jacobs-Sera, Kaylia Edwards, and Bethany Wise. We thank New England Biolabs (NEB) and Integrated DNA Technologies (IDT) for providing reagent support.

## References

Anders, K. R., N. Barekzi, A. A. Best, G. D. Frederick, D. V. Mavrodi et al., 2017 Genome Sequences of Mycobacteriophages Amgine, Amohnition, Bella96, Cain, DarthP, Hammy, Krueger, LastHope, Peanam, PhelpsODU, Phrank, SirPhilip, Slimphazie, and Unicorn. Genome Announc 5: e01202–17.

Catalão, M. J., C. Milho, F. Gil, J. Moniz-Pereira, and M. Pimentel, 2011 A Second Endolysin Gene Is Fully Embedded In-Frame with the lysA Gene of Mycobacteriophage Ms6. Plos One 6: e20515.

Catalão, M. J., and M. Pimentel, 2018 Mycobacteriophage Lysis Enzymes: Targeting the Mycobacterial Cell Envelope. Viruses 10: 428.

Cresawn, S. G., M. Bogel, N. Day, D. Jacobs-Sera, R. W. Hendrix et al., 2011 Phamerator: a bioinformatic tool for comparative bacteriophage genomics. Bmc Bioinformatics 12: 395–395.

Dedrick, R. M., L. J. Marinelli, G. L. Newton, K. Pogliano, J. Pogliano et al., 2013 Functional requirements for bacteriophage growth: gene essentiality and expression in mycobacteriophage Giles. Mol Microbiol 88: 577–589.

Dedrick, R. M., D. Jacobs-Sera, C. A. G. Bustamante, R. A. Garlena, T. N. Mavrich et al., 2017 Prophage-mediated defence against viral attack and viral counter-defence. Nat Microbiol 2: 16251.

Dedrick, R. M., C. A. G. Bustamante, R. A. Garlena, R. S. Pinches, K. Cornely et al., 2019a Mycobacteriophage ZoeJ: A broad host-range close relative of mycobacteriophage TM4. Tuberculosis 115: 14–23.

Dedrick, R. M., C. A. Guerrero-Bustamante, R. A. Garlena, D. A. Russell, K. Ford et al., 2019b Engineered bacteriophages for treatment of a patient with a disseminated drug-resistant Mycobacterium abscessus. Nat Med 25: 730–733.

Ehrt, S., X. V. Guo, C. M. Hickey, M. Ryou, M. Monteleone et al., 2005 Controlling gene expression in mycobacteria with anhydrotetracycline and Tet repressor. Nucleic Acids Res 33: e21–e21.

Gabler, F., S. Nam, S. Till, M. Mirdita, M. Steinegger et al., 2020 Protein Sequence Analysis Using the MPI Bioinformatics Toolkit. Curr Protoc Bioinform 72: e108.

Gauthier, C. H., S. G. Cresawn, and G. F. Hatfull, 2022 PhaMMseqs: a new pipeline for constructing phage gene phamilies using MMseqs2. G3 Genes Genomes Genetics 12: jkac233.

Gentile, G. M., K. S. Wetzel, R. M. Dedrick, M. T. Montgomery, R. A. Garlena et al., 2019 More Evidence of Collusion: a New Prophage-Mediated Viral Defense System Encoded by Mycobacteriophage Sbash. Mbio 10: e00196–19.

Guerrero-Bustamante, C. A., R. M. Dedrick, R. A. Garlena, D. A. Russell, and G. F. Hatfull, 2021 Toward a Phage Cocktail for Tuberculosis: Susceptibility and Tuberculocidal Action of Mycobacteriophages against Diverse Mycobacterium tuberculosis Strains. Mbio 12: e00973–21.

Hallgren, J., K. D. Tsirigos, M. D. Pedersen, J. J. A. Armenteros, P. Marcatili et al., 2022 DeepTMHMM predicts alpha and beta transmembrane proteins using deep neural networks.

Hampton, H. G., B. N. J. Watson, and P. C. Fineran, 2020 The arms race between bacteria and their phage foes. Nature 577: 327–336.

Hatfull, G. F., 2020 Actinobacteriophages: Genomics, Dynamics, and Applications. Ann Rev Virol 7: 37– 61.

Hatfull, G. F., 2018 Mycobacteriophages. Microbiol Spectr 6:.

Heller, D., I. Amaya, A. Mohamed, I. Ali, D. Mavrodi et al., 2022 Systematic overexpression of genes encoded by mycobacteriophage Waterfoul reveals novel inhibitors of mycobacterial growth. G3 Genes Genomes Genetics 12: jkac140.

Heller, D., and V. Sivanathan, 2022 Publishing student-led discoveries in genetics. G3 Genes Genomes Genetics 12: jkac141.

Huiting, E., X. Cao, J. Ren, J. S. Athukoralage, Z. Luo et al. Bacteriophages inhibit and evade cGAS-like immune function in bacteria. Cell 186: 864–876.e21.

Ko, C.-C., and G. F. Hatfull, 2020 Identification of mycobacteriophage toxic genes reveals new features of mycobacterial physiology and morphology. Sci Rep-uk 10: 14670.

Ko, C., and G. F. Hatfull, 2018 Mycobacteriophage Fruitloop gp52 inactivates Wag31 (DivIVA) to prevent heterotypic superinfection. Mol Microbiol 108: 443–460.

LeRoux, M., and M. T. Laub, 2022 Toxin-Antitoxin Systems as Phage Defense Elements. Annu Rev Microbiol 76: 21–43.

Liu, J., M. Dehbi, G. Moeck, F. Arhin, P. Bauda et al., 2004 Antimicrobial drug discovery through bacteriophage genomics. Nat Biotechnol 22: 185–

Madeira, F., M. Pearce, A. R. N. Tivey, P. Basutkar, J. Lee et al., 2022 Search and sequence analysis tools services from EMBL-EBI in 2022. Nucleic Acids Res 50: W276–W279.

Mehla, J., R. M. Dedrick, J. H. Caufield, J. Wagemans, N. Sakhawalkar et al., 2017 Virus-host protein-protein interactions of mycobacteriophage Giles. Sci Rep-uk 7: 16514.

Millman, A., S. Melamed, A. Leavitt, S. Doron, A. Bernheim et al., 2022 An expanded arsenal of immune systems that protect bacteria from phages. Cell Host Microbe 30: 1556–1569.e5.

Miroux, B., and J. E. Walker, 1996 Over-production of Proteins inEscherichia coli: Mutant Hosts that Allow Synthesis of some Membrane Proteins and Globular Proteins at High Levels. J Mol Biol 260: 289–298.

Molshanski-Mor, S., I. Yosef, R. Kiro, R. Edgar, M. Manor et al., 2014 Revealing bacterial targets of growth inhibitors encoded by bacteriophage T7. Proc National Acad Sci 111: 18715–18720.

Montgomery, M. T., C. A. G. Bustamante, R. M. Dedrick, D. Jacobs-Sera, and G. F. Hatfull, 2019 Yet More Evidence of Collusion: a New Viral Defense System Encoded by Gordonia Phage CarolAnn. Mbio 10: e02417–18.

Parikh, A., D. Kumar, Y. Chawla, K. Kurthkoti, S. Khan et al., 2013 Development of a New Generation of Vectors for Gene Expression, Gene Replacement, and Protein-Protein Interaction Studies in Mycobacteria. Appl Environ Microb 79: 1718–1729.

Payne, K. M., and G. F. Hatfull, 2012 Mycobacteriophage Endolysins: Diverse and Modular Enzymes with Multiple Catalytic Activities. Plos One 7: e34052.

Payne, K., Q. Sun, J. Sacchettini, and G. F. Hatfull, 2009 Mycobacteriophage Lysin B is a novel mycolylarabinogalactan esterase. Mol Microbiol 73: 367–381.

Pedulla, M. L., M. E. Ford, J. M. Houtz, T. Karthikeyan, C. Wadsworth et al., 2003 Origins of Highly Mosaic Mycobacteriophage Genomes. Cell 113: 171–182.

Pollenz, R. S., J. Bland, and W. H. Pope, 2022 Bioinformatic characterization of endolysins and holin-like membrane proteins in the lysis cassette of phages that infect Gordonia rubripertincta. Plos One 17: e0276603.

Pope, W. H., C. A. Bowman, D. A. Russell, D. Jacobs-Sera, D. J. Asai et al., 2015 Whole genome comparison of a large collection of mycobacteriophages reveals a continuum of phage genetic diversity. Elife 4: e06416.

Pope, W. H., T. N. Mavrich, R. A. Garlena, C. A. Guerrero-Bustamante, D. Jacobs-Sera et al., 2017 Bacteriophages of Gordonia spp. Display a Spectrum of Diversity and Genetic Relationships. Mbio 8: e01069–17.

Russell, D. A., and G. F. Hatfull, 2016 PhagesDB: the actinobacteriophage database. Bioinformatics 33: 784–786.

Rybniker, J., K. Krumbach, E. van Gumpel, G. Plum, L. Eggeling et al., 2011 The cytotoxic early protein 77 of mycobacteriophage L5 interacts with MSMEG_3532, an L-serine dehydratase of Mycobacterium smegmatis. J Basic Microb 51: 515– 522.

Rybniker, J., A. Nowag, E. V. Gumpel, N. Nissen, N. Robinson et al., 2010 Insights into the function of the WhiB-like protein of mycobacteriophage TM4 – a transcriptional inhibitor of WhiB2. Mol Microbiol 77: 642–657.

Rybniker, J., G. Plum, N. Robinson, P. L. Small, and P. Hartmann, 2008 Identification of three cytotoxic early proteins of mycobacteriophage L5 leading to growth inhibition in Mycobacterium smegmatis. Microbiology+ 154: 2304–2314.

Stokar-Avihail, A., T. Fedorenko, J. Hör, J. Garb, A. Leavitt et al., 2023 Discovery of phage determinants that confer sensitivity to bacterial immune systems. Cell.

Tal, N., B. R. Morehouse, A. Millman, A. Stokar-Avihail, C. Avraham et al., 2021 Cyclic CMP and cyclic UMP mediate bacterial immunity against phages. Cell 184: 5728–5739.e16.

Tran, N. Q., L. F. Rezende, U. Qimron, C. C. Richardson, and S. Tabor, 2008 Gene 1.7 of bacteriophage T7 confers sensitivity of phage growth to dideoxythymidine. Proc National Acad Sci 105: 9373–9378.

Tran, N. Q., S. Tabor, C. J. Amarasiriwardena, A. W. Kulczyk, and C. C. Richardson, 2012 Characterization of a Nucleotide Kinase Encoded by Bacteriophage T7*. J Biol Chem 287: 29468–29478.

Tran, N. Q., S. Tabor, and C. C. Richardson, 2014 Genetic Requirements for Sensitivity of Bacteriophage T7 to Dideoxythymidine. J Bacteriol 196: 2842–2850.

Wagner, S., M. L. Bader, D. Drew, and J.-W. de Gier, 2006 Rationalizing membrane protein overexpression. Trends Biotechnol 24: 364–371.

Zhang, T., H. Tamman,K. C. ‘t Wallant, T. Kurata, M. LeRoux et al., 2022 Direct activation of a bacterial innate immune system by a viral capsid protein. Nature 612: 132–140.

