## Supplemental Materials for "A genome-wide cytotoxicity screen of Cluster F1 mycobacteriophage Girr reveals novel inhibitors of *Mycobacterium smegmatis* growth"

**Supplemental Figure 1:** Shown are the results of representative cytotoxicity assays for the 102 Girr genes screened in this study. Each strain was spotted in triplicate alongside *M. smegmatis*/pExTra-Fruitloop52 (+) and pExTra-Fruitloop52I70S (-) control strains on 7H10 Kan supplemented with 0, 10, or 100 ng/ml aTc. In all experiments, $10^0$ to $10^{-5}$ dilutions are shown. Plates were monitored over 4 or 5 days at 37 ºC, with results shown to best illustrate effects on colony color and size. Colony color was scored using the indicated key shown at the bottom of the data card.

#### Gene *1*; Score *0*

Images taken after 4 days at 37 ºC

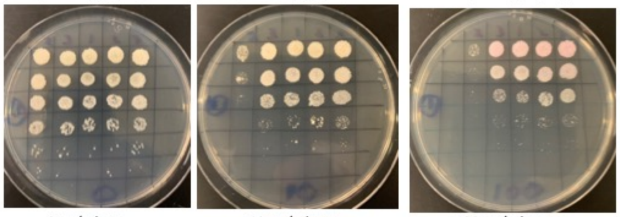


| Lane | Gene ID | Plasmid name | Gene name | Toxic/Non-toxic | Colony color on 100 ng/ml aTc plate* |
|---|---|---|---|---|---|
| + Toxic control | -- | pExTra02 | Fruitloop *52* | Toxic | - |
| - Non-toxic control | -- | pExTra03 | Fruitloop *52* mutant | Non-toxic | ++ |
| 1 | -- | pExTra-Girr1 | Girr *1* replicate 1 | Non-toxic | ++ |
| 2 | -- | pExTra-Girr1 | Girr *1* replicate 2 | Non-toxic | ++ |
| 3 | -- | pExTra-Girr1 | Girr *1* replicate 3 | Non-toxic | ++ |

*Key: NG (no growth) - (no pink color) +(faint pink color) ++(obvious pink color) +++ (dark pink color)

#### Gene *2*; Score *3*

Images taken after 4 days at 37 ºC

REPLICATE EXPERIMENT

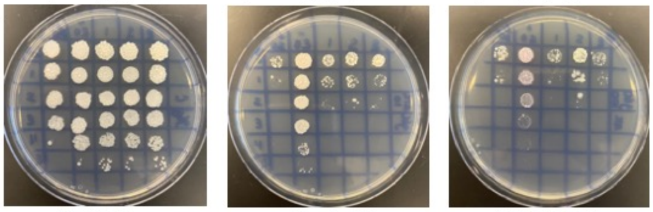


| Lane | Gene ID | Plasmid name | Gene name | Toxic/Non-toxic | Colony color on 100 ng/ml aTc plate* |
|---|---|---|---|---|---|
| + Toxic control | -- | pExTra02 | Fruitloop *52* | Toxic | - |
| - Non-toxic control | -- | pExTra03 | Fruitloop *52* mutant | Non-toxic | ++ |
| 1 | -- | pExTra-Girr2 | Girr *2* replicate 1 | Toxic | - |
| 2 | -- | pExTra-Girr2 | Girr *2* replicate 2 | Toxic | - |
| 3 | -- | pExTra-Girr2 | Girr *2* replicate 3 | Toxic | - |

*Key: NG (no growth) - (no pink color) +(faint pink color) ++(obvious pink color) +++ (dark pink color)

#### Gene *3*; Score *0*

Images taken after 4 days at 37 ºC on 7H11 agar

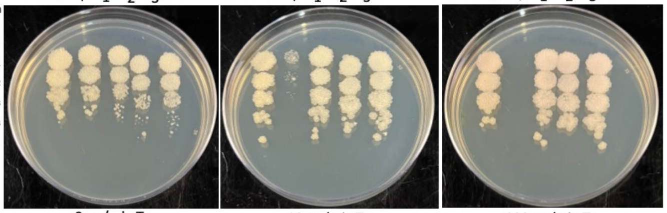


| Lane | Plasmid name | Gene name, replicate | Toxic/Non-toxic | Colony color on 100 ng/ml aTc plate* |
|---|---|---|---|---|
| - Non-toxic control | pExTra03 | Fruitloop *52* mutant | Non-toxic | + |
| + Toxic control | pExTra02 | Fruitloop *52* | Toxic | - |
| 1 | pExTra-Girr*3* | Girr *3* replicate 1 | Non-toxic | + |
| 2 | pExTra-Girr*3* | Girr *3* replicate 2 | Non-toxic | + |
| 3 | pExTra-Girr*3* | Girr *3* replicate 3 | Non-toxic | + |

*Key: NG (no growth) - (no pink color) +(faint pink color) ++(obvious pink color) +++ (dark pink color)

#### Gene *4*; Score *0*

Images taken after 5 days at 37 ºC

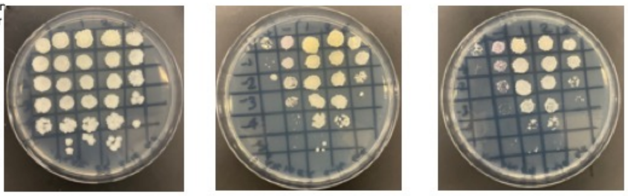


| Lane | Gene ID | Plasmid name | Gene name | Toxic/Non-toxic | Colony color on 100 ng/ml aTc plate* |
|---|---|---|---|---|---|
| + Toxic control | -- | pExTra02 | Fruitloop *52* | Toxic | - |
| - Non-toxic control | -- | pExTra03 | Fruitloop *52* mutant | Non-toxic | + |
| 1 | ----- | pExTra-Girr4 | Girr *4* replicate 1 | Non-toxic | - |
| 2 | ----- | pExTra-Girr4 | Girr *4* replicate 2 | Non-toxic | - |
| 3 | --------- | pExTra-Girr4 | Girr *4* replicate 3 | Non-toxic | - |

*Key: NG (no growth) - (no pink color) +(faint pink color) ++(obvious pink color) +++ (dark pink color)

#### Gene *5*; Score *2*

Images taken after 4 days at 37 ºC

REPLCATE EXPERIMENT 2023

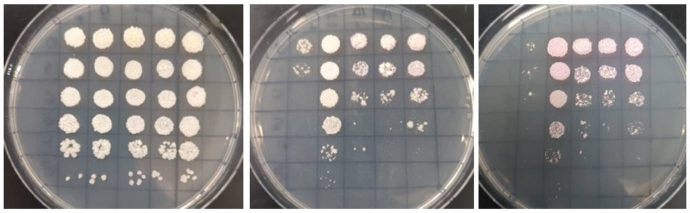


| Lane | Gene ID | Plasmid name | Gene name | Toxic/Non-toxic | Colony color on 100 ng/ml aTc plate* |
|---|---|---|---|---|---|
| + Toxic control | -- | pExTra02 | Fruitloop *52* | Toxic | - |
| - Non-toxic control | -- | pExTra03 | Fruitloop *52* mutant | Non-toxic | ++ |
| 1 | -- | pExTra-Girr5 | Girr 5 replicate 1 | Toxic | ++ |
| 2 | -- | pExTra-Girr5 | Girr 5 replicate 2 | Toxic | ++ |
| 3 | -- | pExTra-Girr5 | Girr 5 replicate 3 | Toxic | ++ |

*Key: NG (no growth) - (no pink color) +(faint pink color) ++(obvious pink color) +++ (dark pink color)

#### Gene *6*; Score *1*

Images taken after 5 days at 37 ºC

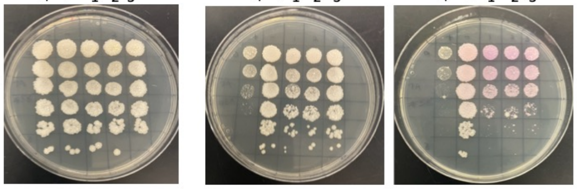


| Lane | Gene ID | Plasmid name | Gene name | Toxic/Non-toxic | Colony color on 100 ng/ml aTc plate* |
|---|---|---|---|---|---|
| + Toxic control | -- | pExTra02 | Fruitloop 52 | Toxic | - |
| - Non-toxic control | -- | pExTra03 | Fruitloop 52 mutant | Non-toxic | + |
| 1 | -- | pExTra-Girr6 | Girr *6* replicate 1 | Toxic | ++ |
| 2 | -- | pExTra-Girr6 | Girr *6* replicate 2 | Toxic | ++ |
| 3 | -- | pExTra-Avani58 | Girr *6* replicate 3 | Toxic | ++ |

*Key: NG (no growth) - (no pink color) +(faint pink color) ++(obvious pink color) +++ (dark pink color)

#### Gene *7*; Score *0*

Images taken after 4 days at 37 ºC

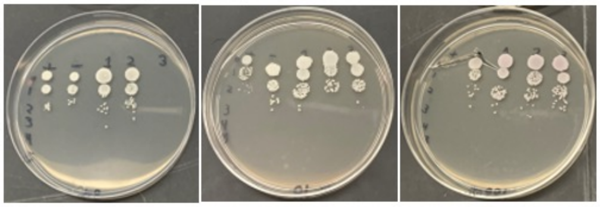


| Lane | Gene ID | Plasmid name | Gene name | Toxic/Non-toxic | Colony color on 100 ng/ml aTc plate* |
|---|---|---|---|---|---|
| + Toxic control | -- | pExTra02 | Fruitloop *52* | Toxic | - |
| - Non-toxic control | -- | pExTra03 | Fruitloop *52 mutant* | Non-toxic | + |
| 1 | -- | pExTra-Girr7 | Girr 7 replicate 1 | Non-toxic | + |
| 2 | -- | pExTra-Girr7 | Girr 7 replicate 2 | Non-toxic | + |
| 3 | -- | pExTra-Girr7 | Girr 7 replicate 3 | Non-toxic | + |

*Key: NG (no growth) - (no pink color) +(faint pink color) ++(obvious pink color) +++ (dark pink color)

#### Gene *8*; Score *0*

Images taken after 4 days at 37 ºC on 7H11 agar

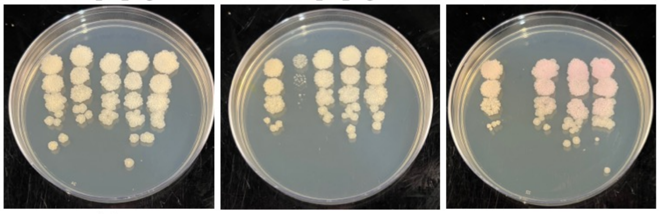


| Lane | Plasmid name | Gene name, replicate | Toxic/Non-toxic | Colony color on 100 ng/ml aTc plate* |
|---|---|---|---|---|
| - Non-toxic control | pExTra03 | Fruitloop *52* mutant | Non-toxic | + |
| + Toxic control | pExTra02 | Fruitloop *52* | Toxic | - |
| 1 | pExTra-Girr*8* | Girr *8* replicate 1 | Non-toxic | ++ |
| 2 | pExTra-Girr*8* | Girr *8* replicate 2 | Non-toxic | ++ |
| 3 | pExTra-Girr*8* | Girr *8* replicate 3 | Non-toxic | ++ |

*Key: NG (no growth) - (no pink color) +(faint pink color) ++(obvious pink color) +++ (dark pink color)

### Gene *9*; Score *0*

Images taken after 5 days at 37 ºC

Replicate experiment

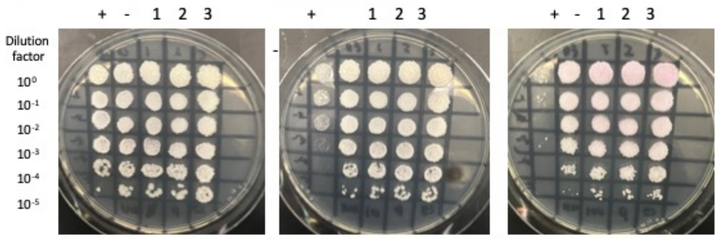


0 ng/ml aTc | 10 ng/ml aTc | 100 ng/ml aTc

| Lane | Gene ID | Plasmid name | Gene name | Toxic/Non-toxic | Colony color on 100 ng/ml aTc plate* |
|---|---|---|---|---|---|
| + Toxic control | -- | pExTra02 | Fruitloop *52* | Toxic | - |
| - Non-toxic control | -- | pExTra03 | Fruitloop *52* mutant | Non-toxic | + |
| 1 | -- | pExTra-Girr9 | Girr 9 replicate 1 | Non-toxic | + |
| 2 | -- | pExTra-Girr9 | Girr 9 replicate 2 | Non-toxic | + |
| 3 | -- | pExTra-Girr9 | Girr 9 replicate 3 | Non-toxic | + |

*Key: NG (no growth) - (no pink color) +(faint pink color) ++(obvious pink color) +++ (dark pink color)

### Gene *10*; Score *0*

Images taken after 4 days at 37 ºC

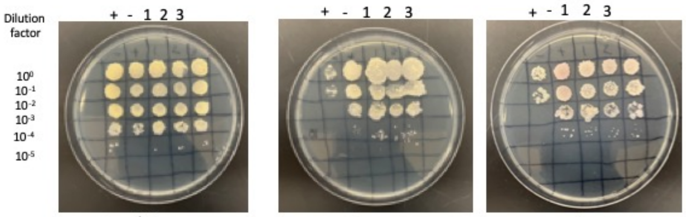


0 ng/ml aTc | 10 ng/ml aTc | 100 ng/ml aTc

| Lane | Gene ID | Plasmid name | Gene name | Toxic/Non-toxic | Colony color on 100 ng/ml aTc plate* |
|---|---|---|---|---|---|
| + Toxic control | -- | pExTra02 | Fruitloop *52* | Toxic | - |
| - Non-toxic control | -- | pExTra03 | Fruitloop *52* mutant | Non-toxic | ++ |
| 1 | 60329149 | pExTra-Girr10 | Girr *10* replicate 1 | Non-Toxic | - |
| 2 | 60329149 | pExTra-Girr10 | Girr *10* replicate 2 | Non-Toxic | - |
| 3 | 60329149 | pExTra-Girr10 | Girr *10* replicate 3 | Non-Toxic | - |

*Key: NG (no growth) - (no pink color) +(faint pink color) ++(obvious pink color) +++ (dark pink color)

### Gene *11*; Score *0*

Images taken after 4 days at 37 ºC

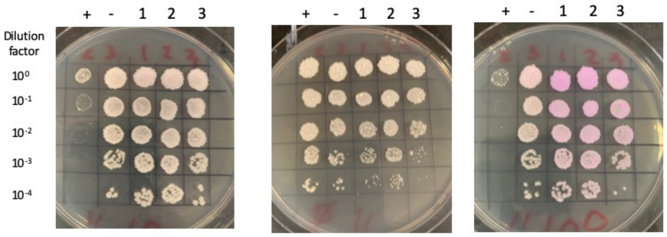


0 ng/ml aTc | 10 ng/ml aTc | 100 ng/ml aTc

| Lane | Gene ID | Plasmid name | Gene name | Toxic/Non-toxic | Colony color on 100 ng/ml aTc plate* |
|---|---|---|---|---|---|
| + Toxic control | -- | pExTra02 | Fruitloop *52* | Toxic | - |
| - Non-toxic control | -- | pExTra03 | Fruitloop *52* mutant | Non-toxic | ++ |
| 1 | -- | pExTra-Girr11 | Girr *11* replicate 1 | Non-toxic | +++ |
| 2 | -- | pExTra-Girr11 | Girr *11* replicate 2 | Non-toxic | +++ |
| 3 | -- | pExTra-Girr11 | Girr *11* replicate 3 | Non-toxic | +++ |

*Key: NG (no growth) - (no pink color) +(faint pink color) ++(obvious pink color) +++ (dark pink color)

### Gene *12*; Score *0*

Images taken after 4 days at 37 ºC

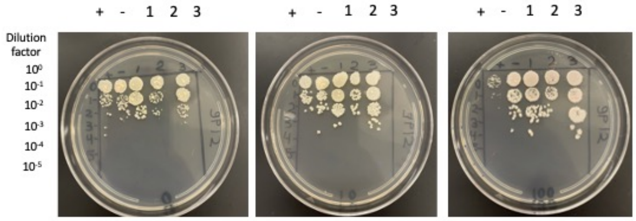


0 ng/ml aTc | 10 ng/ml aTc | 100 ng/ml aTc

| Lane | Gene ID | Plasmid name | Gene name | Toxic/Non-toxic | Colony color on 100 ng/ml aTc plate* |
|---|---|---|---|---|---|
| + Toxic control | -- | pExTra02 | Fruitloop *52* | Toxic | - |
| - Non-toxic control | -- | pExTra03 | Fruitloop *52 mutant* | Non-toxic | + |
| 1 | | pExTra-Girr 12 | Girr *12* replicate 1 | Non-toxic | + |
| 2 | | pExTra-Girr 12 | Girr *12* replicate 2 | Non-toxic | + |
| 3 | | pExTra-Girr 12 | Girr *12* replicate 3 | Non-toxic | + |

*Key: NG (no growth) - (no pink color) +(faint pink color) ++(obvious pink color) +++ (dark pink color)

### Gene *13*; Score *0*

Images taken after 5 days at 37 ºC

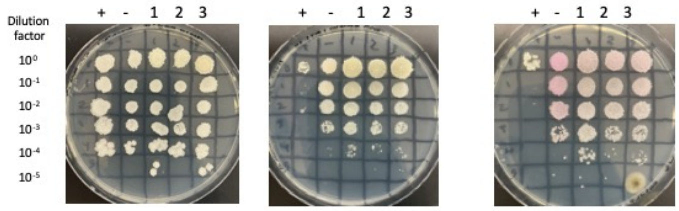


0 ng/ml aTc | 10 ng/ml aTc | 100 ng/ml aTc

| Lane | Gene ID | Plasmid name | Gene name | Toxic/Non-toxic | Colony color on 100 ng/ml aTc plate* |
|---|---|---|---|---|---|
| + Toxic control | -- | pExTra02 | Fruitloop *52* | Toxic | - |
| - Non-toxic control | -- | pExTra03 | Fruitloop *52* mutant | Non-toxic | ++ |
| 1 | -- | pExTra-Girr13 | Girr *13* replicate 1 | Non-toxic | + |
| 2 | -- | pExTra-Girr13 | Girr *13* replicate 2 | Non-toxic | + |
| 3 | -- | pExTra-Girr13 | Girr *13* replicate 3 | Non-toxic | + |

*Key: NG (no growth) - (no pink color) +(faint pink color) ++(obvious pink color) +++ (dark pink color)

### Gene *14*; Score *0*

Images taken after 4 days at 37 ºC

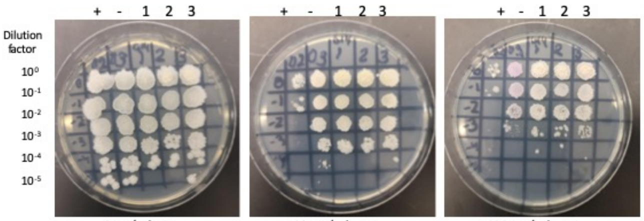


0 ng/ml aTc | 10 ng/ml aTc | 100 ng/ml aTc

| Lane | Gene ID | Plasmid name | Gene name | Toxic/Non-toxic | Colony color on 100 ng/ml aTc plate* |
|---|---|---|---|---|---|
| + Toxic control | -- | pExTra02 | Fruitloop *52* | Toxic | - |
| - Non-toxic control | -- | pExTra03 | Fruitloop *52* mutant | Non-toxic | ++ |
| 1 | 60329152 | pExTra-Girr14 | Girr *14* replicate 1 | Non-toxic | + |
| 2 | 60329152 | pExTra-Girr14 | Girr *14* replicate 2 | Non-toxic | + |
| 3 | 60329152 | pExTra-Girr14 | Girr *14* replicate 3 | Non-toxic | + |

*Key: NG (no growth) - (no pink color) +(faint pink color) ++(obvious pink color) +++ (dark pink color)

### Gene *15*; Score *0*

Images taken after 5 days at 37 ºC

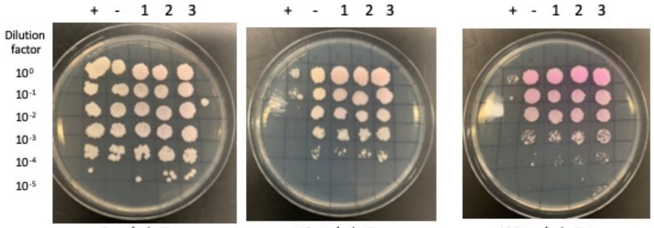


0 ng/ml aTc | 10 ng/ml aTc | 100 ng/ml aTc

| Lane | Gene ID | Plasmid name | Gene name | Toxic/Non-toxic | Colony color on 100 ng/ml aTc plate* |
|---|---|---|---|---|---|
| + Toxic control | -- | pExTra02 | Fruitloop *52* | Toxic | - |
| - Non-toxic control | -- | pExTra03 | Fruitloop *52* mutant | Non-toxic | +++ |
| 1 | -- | pExTra-Girr15 | Girr *15* replicate 1 | Non-toxic | +++ |
| 2 | -- | pExTra-Girr15 | Girr *15* replicate 2 | Non-toxic | +++ |
| 3 | -- | pExTra-Girr15 | Girr *15* replicate 3 | Non-toxic | +++ |

*Key: NG (no growth) - (no pink color) +(faint pink color) ++(obvious pink color) +++ (dark pink color)

### Gene *16*; Score *0*

Images taken after 5 days at 37 ºC

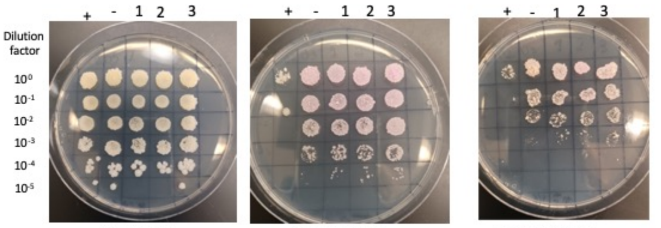


0 ng/ml aTc | 10 ng/ml aTc | 100 ng/ml aTc

| Lane | Gene ID | Plasmid name | Gene name | Toxic/Non-toxic | Colony color on 100 ng/ml aTc plate* |
|---|---|---|---|---|---|
| + Toxic control | -- | pExTra02 | Fruitloop *52* | Toxic | - |
| - Non-toxic control | -- | pExTra03 | Fruitloop *52* mutant | Non-toxic | + |
| 1 | -- | pExTra-Girr16 | Girr 16 replicate 1 | Non-toxic | + |
| 2 | -- | pExTra-Girr16 | Girr *16* replicate 2 | Non-toxic | ++ |
| 3 | -- | pExTra-Girr16 | Girr *16* replicate 3 | Non-toxic | + |

*Key: NG (no growth) - (no pink color) +(faint pink color) ++(obvious pink color) +++ (dark pink color)

#### Gene *17*; Score *0*

Images taken after 5 days at 37 ºC

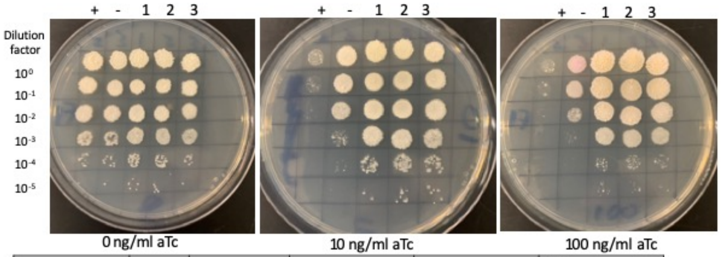


| Lane | Gene ID | Plasmid name | Gene name | Toxic/Non-toxic | Colony color on 100 ng/ml aTc plate* |
|---|---|---|---|---|---|
| + Toxic control | -- | pExTra02 | Fruitloop *52* | Toxic | - |
| - Non-toxic control | -- | pExTra03 | Fruitloop *52* mutant | Non-toxic | ++ |
| 1 | -- | pExTra-Girr17 | Girr *17* replicate 1 | Non-toxic | - |
| 2 | -- | pExTra-Girr17 | Girr *17* replicate 2 | Non-toxic | - |
| 3 | -- | pExTra-Girr17 | Girr *17* replicate 3 | Non-toxic | - |

*Key: NG (no growth) - (no pink color) +(faint pink color) ++(obvious pink color) +++ (dark pink color)

#### Gene *18*; Score 0

Images taken after 5 days at 37 ºC

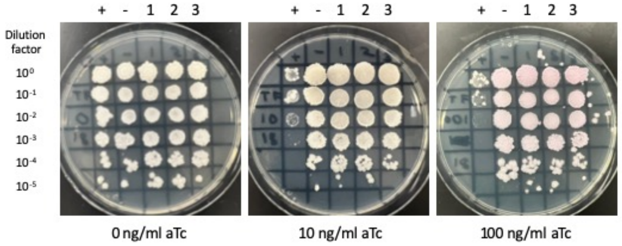


| Lane | Gene ID | Plasmid name | Gene name | Toxic/Non-toxic | Colony color on 100 ng/ml aTc plate* |
|---|---|---|---|---|---|
| + Toxic control | -- | pExTra02 | Fruitloop *52* | Toxic | - |
| - Non-toxic control | -- | pExTra03 | Fruitloop *52* mutant | Non-toxic | ++ |
| 1 | -- | pExTra-Girr18 | Girr *18* replicate 1 | Non-toxic | ++ |
| 2 | -- | pExTra-Girr18 | Girr *18* replicate 2 | Non-toxic | ++ |
| 3 | -- | pExTra-Girr18 | Girr *18* replicate 3 | Non-toxic | ++ |

*Key: NG (no growth) - (no pink color) +(faint pink color) ++(obvious pink color) +++ (dark pink color)

#### Gene *19*; Score *2*

Images taken after 4 days at 37 ºC

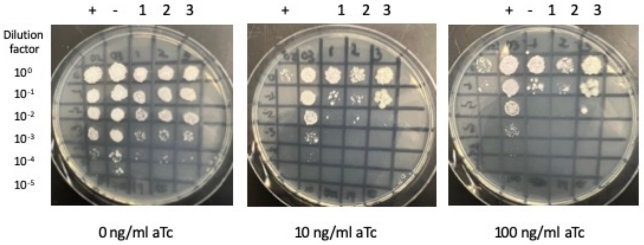


| Lane | Gene ID | Plasmid name | Gene name | Toxic/Non-toxic | Colony color on 100 ng/ml aTc plate* |
|---|---|---|---|---|---|
| + Toxic control | -- | pExTra02 | Fruitloop *52* | Toxic | - |
| - Non-toxic control | -- | pExTra03 | Fruitloop *52* mutant | Non-toxic | + |
| 1 | -- | pExTra-Girr19 | Girr 19 replicate 1 | toxic | + |
| 2 | -- | pExTra-Girr19 | Girr 19 replicate 2 | toxic | + |
| 3 | -- | pExTra-Girr19 | Girr 19 replicate 3 | toxic | + |

*Key: NG (no growth) - (no pink color) +(faint pink color) ++(obvious pink color) +++ (dark pink color)

#### Gene *20*; Score *0*

Images taken after 4 days at 37 ºC

REPEAT EXPERIMENT

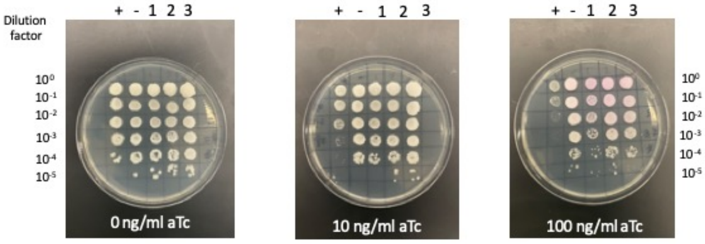


| Lane | Gene ID | Plasmid name | Gene name | Toxic/Non-toxic | Colony color on 100 ng/ml aTc plate* |
|---|---|---|---|---|---|
| + Toxic control | -- | pExTra02 | Fruitloop *52* | Toxic | - |
| - Non-toxic control | -- | pExTra03 | Fruitloop *52* mutant | Non-toxic | + |
| 1 | 60329159 | pExTra-Girr20 | Girr *20* replicate 1 | Non-Toxic | ++ |
| 2 | 60329159 | pExTra-Girr20 | Girr *20* replicate 2 | Non-Toxic | ++ |
| 3 | 60329159 | pExTra-Girr20 | Girr *20* replicate 3 | Non-Toxic | ++ |

*Key: NG (no growth) - (no pink color) +(faint pink color) ++(obvious pink color) +++ (dark pink color)

#### Gene *21*; Score *0*

Images taken after 5 days at 37 ºC

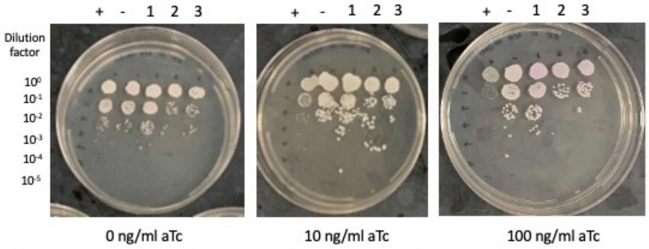


| Lane | Gene ID | Plasmid name | Gene name | Toxic/Non-toxic | Colony color on 100 ng/ml aTc plate* |
|---|---|---|---|---|---|
| + Toxic control | -- | pExTra02 | Fruitloop *52* | Toxic | - |
| - Non-toxic control | -- | pExTra03 | Fruitloop *52 mutant* | Non-toxic | + |
| 1 | -- | pExTra-Girr 21 | Girr 21 replicate 1 | Non-toxic | ++ |
| 2 | -- | pExTra-Girr 21 | Girr 21replicate 2 | Non-toxic | ++ |
| 3 | -- | pExTra-Girr 21 | Girr 21replicate 3 | Non-toxic | ++ |

*Key: NG (no growth) - (no pink color) +(faint pink color) ++(obvious pink color) +++ (dark pink color)

#### Gene *22*; Score *2*

Images taken after 4 days at 37 ºC, 3 days at 4 ºC FIRST EXPERIMENT

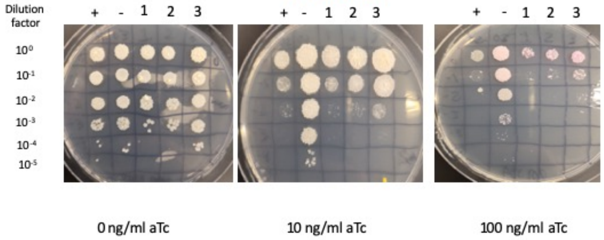


| Lane | Gene ID | Plasmid name | Gene name | Toxic/Non-toxic | Colony color on 100 ng/ml aTc plate* |
|---|---|---|---|---|---|
| + Toxic control | -- | pExTra02 | Fruitloop *52* | Toxic | - |
| - Non-toxic control | -- | pExTra03 | Fruitloop *52* mutant | Non-toxic | ++ |
| 1 | -- | pExTra-Girr22 | Girr *22* replicate 1 | Toxic | ++ |
| 2 | -- | pExTra-Girr22 | Girr *22* replicate 2 | Toxic | ++ |
| 3 | -- | pExTra-Girr22 | Girr *22* replicate 3 | Toxic | ++ |

*Key: NG (no growth) - (no pink color) +(faint pink color) ++(obvious pink color) +++ (dark pink color)

#### Gene *23*; Score *0*

Images taken after 4 days at 37 ºC

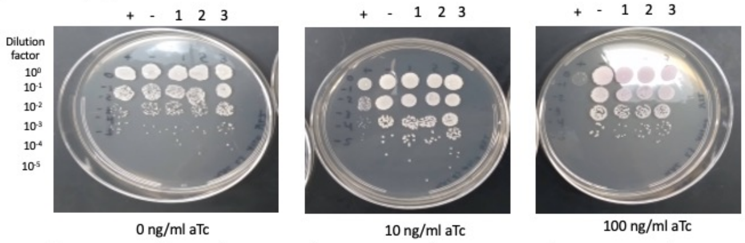


| Lane | Gene ID | Plasmid name | Gene name | Toxic/Non-toxic | Colony color on 100 ng/ml aTc plate* |
|---|---|---|---|---|---|
| + Toxic control | -- | pExTra02 | Fruitloop *52* | Toxic | - |
| - Non-toxic control | -- | pExTra03 | Fruitloop *52 mutant* | Non-toxic | + |
| 1 | | pExTra-Girr23 | Girr 23 replicate 1 | Non-toxic | ++ |
| 2 | | pExTra-Girr23 | Girr 23 replicate 2 | Non-toxic | ++ |
| 3 | | pExTra-Girr23 | Girr 23 replicate 3 | Non-toxic | ++ |

*Key: NG (no growth) - (no pink color) +(faint pink color) ++(obvious pink color) +++ (dark pink color)

#### Gene *24*; Score *0*

Images taken after 4 days at 37 ºC

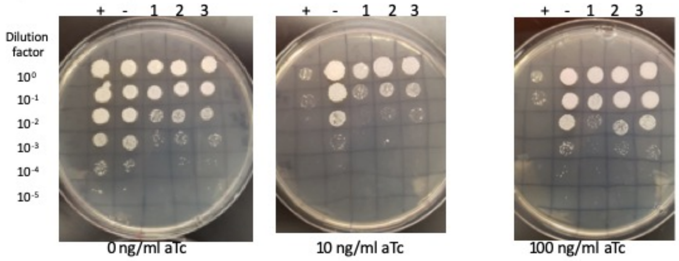


| Lane | Gene ID | Plasmid name | Gene name | Toxic/Non-toxic | Colony color on 100 ng/ml aTc plate* |
|---|---|---|---|---|---|
| + Toxic control | 6940650 | pExTra02 | Fruitloop *52* | Toxic | - |
| - Non-toxic control | 6940650 | pExTra03 | Fruitloop *52* mutant | Non-toxic | ++ |
| 1 | 60329163 | pExTra-Girr24 | Girr *24* replicate 1 | Non-toxic | + |
| 2 | 60329163 | pExTra-Girr24 | Girr *24* replicate 2 | Non-toxic | + |
| 3 | 60329163 | pExTra-Girr24 | Girr *24* replicate 3 | Non-toxic | + |

*Key: NG (no growth) - (no pink color) +(faint pink color) ++(obvious pink color) +++ (dark pink color)

#### Gene *25*; Score *0*

Images taken after 4 days at 37 ºC REPLCATE EXPERIMENT

\+ - 1 2 3 | + - 1 2 3 | + - 1 2 3

Dilution factor: $10^0$, $10^{-1}$, $10^{-2}$, $10^{-3}$, $10^{-4}$, $10^{-5}$

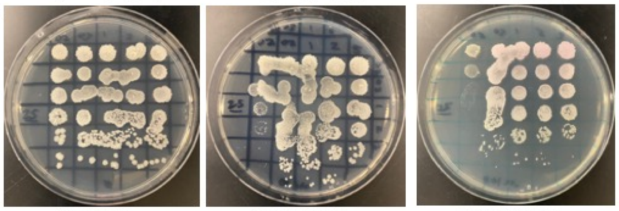

0 ng/ml aTc | 10 ng/ml aTc | 100 ng/ml aTc

| Lane | Gene ID | Plasmid name | Gene name | Toxic/Non-toxic | Colony color on 100 ng/ml aTc plate* |
|---|---|---|---|---|---|
| + Toxic control | -- | pExTra02 | Fruitloop *52* | Toxic | - |
| - Non-toxic control | -- | pExTra03 | Fruitloop *52* mutant | Non-toxic | ++ |
| 1 | -- | pExTra-Girr25 | Girr *25* replicate 1 | Non-toxic | ++ |
| 2 | -- | pExTra-Girr25 | Girr *25* replicate 2 | Non-toxic | ++ |
| 3 | -- | pExTra-Girr25 | Girr *25* replicate 3 | Non-toxic | ++ |

*Key: NG (no growth) - (no pink color) +(faint pink color) ++(obvious pink color) +++ (dark pink color)

#### Gene *26*; Score *0*

Images taken after 4 days at 37 ºC

\+ - 1 2 3 | + - 1 2 3 | + - 1 2 3

Dilution factor: $10^0$, $10^{-1}$, $10^{-2}$, $10^{-3}$, $10^{-4}$, $10^{-5}$

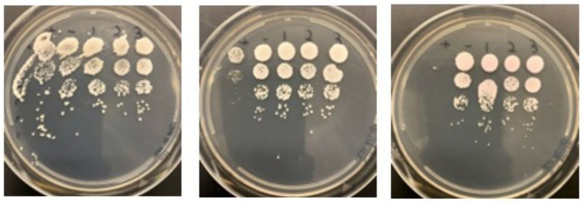

0 ng/ml aTc | 10 ng/ml aTc | 100 ng/ml aTc

| Lane | Gene ID | Plasmid name | Gene name | Toxic/Non-toxic | Colony color on 100 ng/ml aTc plate* |
|---|---|---|---|---|---|
| + Toxic control | -- | pExTra02 | Fruitloop *52* | Toxic | - |
| - Non-toxic control | -- | pExTra03 | Fruitloop *52 mutant* | Non-toxic | + |
| 1 | -- | pExTra-Girr 26 | Girr *26* replicate 1 | Non-toxic | + |
| 2 | -- | pExTra-Girr 26 | *Girr 26* replicate 2 | Non-toxic | + |
| 3 | -- | pExTra-Girr 26 | *Girr 26* replicate 3 | Non-toxic | + |

*Key: NG (no growth) - (no pink color) +(faint pink color) ++(obvious pink color) +++ (dark pink color)

#### Gene *27*; Score *0*

Images taken after 3 days at 37 ºC on 7H11 agar

\- + 1 2 3 | - + 1 2 3 | - + 1 2 3

Dilution factor: $10^0$, $10^{-1}$, $10^{-2}$, $10^{-3}$, $10^{-4}$, $10^{-5}$

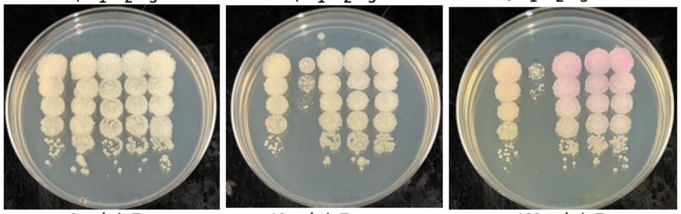

0 ng/ml aTc | 10 ng/ml aTc | 100 ng/ml aTc

| Lane | Plasmid name | Gene name, replicate | Toxic/Non-toxic | Colony color on 100 ng/ml aTc plate* |
|---|---|---|---|---|
| - Non-toxic control | pExTra03 | Fruitloop *52* mutant | Non-toxic | + |
| + Toxic control | pExTra02 | Fruitloop *52* | Toxic | - |
| 1 | pExTra-Girr*27* | Girr *27* replicate 1 | Non-toxic | ++ |
| 2 | pExTra-Girr*27* | Girr *27* replicate 2 | Non-toxic | ++ |
| 3 | pExTra-Girr*27* | Girr *27* replicate 3 | Non-toxic | ++ |

*Key: NG (no growth) - (no pink color) +(faint pink color) ++(obvious pink color) +++ (dark pink color)

#### Gene *28*; Score *0*

Images taken after 4 days at 37 ºC

\+ - 1 2 3 | + - 1 2 3 | + - 1 2 3

Dilution factor: $10^0$, $10^{-1}$, $10^{-2}$, $10^{-3}$, $10^{-4}$, $10^{-5}$

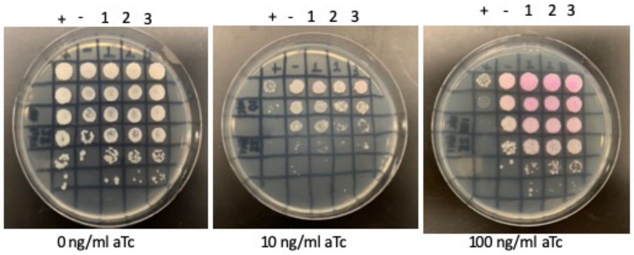

0 ng/ml aTc | 10 ng/ml aTc | 100 ng/ml aTc

| Lane | Gene ID | Plasmid name | Gene name | Toxic/Non-toxic | Colony color on 100 ng/ml aTc plate* |
|---|---|---|---|---|---|
| + Toxic control | -- | pExTra02 | Fruitloop *52* | Toxic | - |
| - Non-toxic control | -- | pExTra03 | Fruitloop *52* mutant | Non-toxic | + |
| 1 | -- | pExTra-Girr28 | Girr *28* replicate 1 | Non-toxic | ++ |
| 2 | -- | pExTra-Girr28 | Girr *28* replicate 2 | Non-toxic | ++ |
| 3 | -- | pExTra-Girr28 | Girr *28* replicate 3 | Non-toxic | ++ |

*Key: NG (no growth) - (no pink color) +(faint pink color) ++(obvious pink color) +++ (dark pink color)

#### Gene *29*; Score *0*

Images taken after 5 days at 37 ºC

\+ - 1 2 3 | + - 1 2 3 | + - 1 2 3

Dilution factor: $10^0$, $10^{-1}$, $10^{-2}$, $10^{-3}$, $10^{-4}$, $10^{-5}$

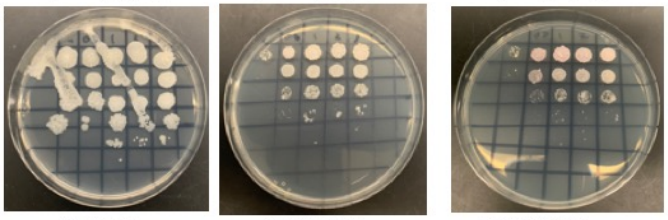

0 ng/ml aTc | 10 ng/ml aTc | 100 ng/ml aTc

| Lane | Gene ID | Plasmid name | Gene name | Toxic/Non-toxic | Colony color on 100 ng/ml aTc plate* |
|---|---|---|---|---|---|
| + Toxic control | -- | pExTra02 | Fruitloop *52* | Toxic | - |
| - Non-toxic control | -- | pExTra03 | Fruitloop *52* mutant | Non-toxic | + |
| 1 | -- | pExTra-Girr29 | Girr *29* replicate 1 | Non-toxic | + |
| 2 | -- | pExTra-Girr29 | Girr *29* replicate 2 | Non-toxic | + |
| 3 | -- | pExTra-Girr29 | Girr *29* replicate 3 | Non-toxic | + |

*Key: NG (no growth) - (no pink color) +(faint pink color) ++(obvious pink color) +++ (dark pink color)

#### Gene *30*; Score *0*

Images taken after 4 days at 37 ºC

\+ - 1 2 3 | + - 1 2 3 | + - 1 2 3

Dilution factor: $10^0$, $10^{-1}$, $10^{-2}$, $10^{-3}$, $10^{-4}$, $10^{-5}$

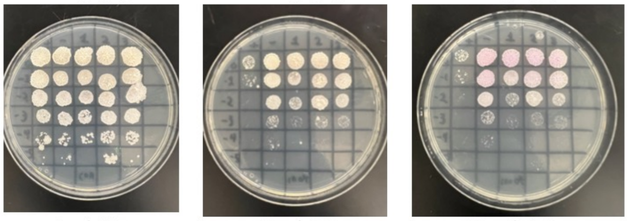

0 ng/ml aTc | 10 ng/ml aTc | 100 ng/ml aTc

| Lane | Gene ID | Plasmid name | Gene name | Toxic/Non-toxic | Colony color on 100 ng/ml aTc plate* |
|---|---|---|---|---|---|
| + Toxic control | -- | pExTra02 | Fruitloop *52* | Toxic | - |
| - Non-toxic control | -- | pExTra03 | Fruitloop *52 I/S* | Non-toxic | ++ |
| 1 | NKF | pExTra-Girr30 | Girr *30* replicate 1 | Non-toxic | ++ |
| 2 | NKF | pExTra-Girr30 | Girr *30* replicate 1 | Non-toxic | ++ |
| 3 | NKF | pExTra-Girr30 | Girr *30* replicate 1 | Non-toxic | ++ |

*Key: NG (no growth) - (no pink color) +(faint pink color) ++(obvious pink color) +++ (dark pink color)

#### Gene *31*; Score *1*

Images taken after 3 days at 37 ºC REPLICATE EXPERIMENT #2

\+ - 1 2 3 | + - 1 2 3 | + - 1 2 3

Dilution factor: $10^0$, $10^{-1}$, $10^{-2}$, $10^{-3}$, $10^{-4}$, $10^{-5}$

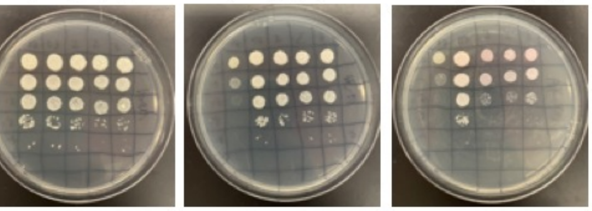

0 ng/ml aTc | 10 ng/ml aTc | 100 ng/ml aTc

| Lane | Gene ID | Plasmid name | Gene name | Toxic/Non-toxic | Colony color on 100 ng/ml aTc plate* |
|---|---|---|---|---|---|
| + Toxic control | -- | pExTra02 | Fruitloop *52* | Toxic | - |
| - Non-toxic control | -- | pExTra03 | Fruitloop *52* mutant | Non-toxic | + |
| 1 | -- | pExTra-Girr31 | Girr 31 replicate 1 | Toxic | ++ |
| 2 | -- | pExTra-Girr31 | Girr *31* replicate 2 | Toxic | + |
| 3 | -- | pExTra-Girr31 | Girr *31* replicate 3 | Toxic | ++ |

*Key: NG (no growth) - (no pink color) +(faint pink color) ++(obvious pink color) +++ (dark pink color)

#### Gene *32*; Score *2*

Images taken after 4 days at 37 ºC REPLCATE EXPERIMENT 2023

\+ - 1 2 3 | + - 1 2 3 | + - 1 2 3

Dilution factor: $10^0$, $10^{-1}$, $10^{-2}$, $10^{-3}$, $10^{-4}$, $10^{-5}$

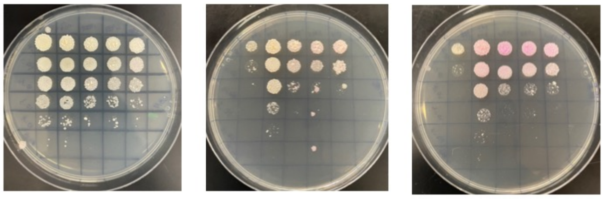

0 ng/ml aTc | 10 ng/ml aTc | 100 ng/ml aTc

| Lane | Gene ID | Plasmid name | Gene name | Toxic/Non-toxic | Colony color on 100 ng/ml aTc plate* |
|---|---|---|---|---|---|
| + Toxic control | | pExTra02 | Fruitloop *52* | Toxic | - |
| - Non-toxic control | -- | pExTra03 | Fruitloop *52* mutant | Non-toxic | ++ |
| 1 | | pExTra-Girr32 | Girr 32 replicate 1 | Toxic | +++ |
| 2 | | pExTra-Girr32 | Girr 32 replicate 2 | Toxic | ++ |
| 3 | | pExTra-Girr32 | Girr 32 replicate 3 | Toxic | ++ |

*Key: NG (no growth) - (no pink color) +(faint pink color) ++(obvious pink color) +++ (dark pink color)

#### Gene *33*; Score *0*

Images taken after 4 days at 37 ºC

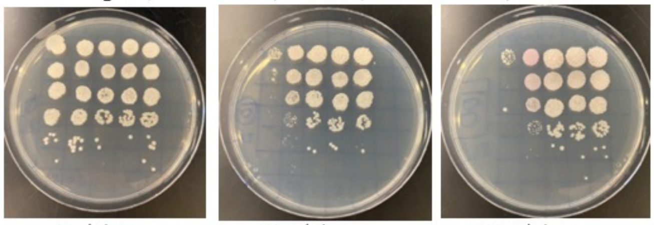


0 ng/ml aTc — 10 ng/ml aTc — 100 ng/ml aTc

| Lane | Gene ID | Plasmid name | Gene name | Toxic/Non-toxic | Colony color on 100 ng/ml aTc plate* |
|---|---|---|---|---|---|
| + Toxic control | -- | pExTra02 | Fruitloop *52* | Toxic | - |
| - Non-toxic control | -- | pExTra03 | Fruitloop *52* mutant | Non-toxic | ++ |
| 1 | -- | pExTra-Girr33 | Girr *33* replicate 1 | Non-toxic | + |
| 2 | -- | pExTra-Girr33 | Girr 33 replicate 2 | Non-toxic | + |
| 3 | -- | pExTra-Girr33 | Girr *33* replicate 3 | Non-toxic | + |

*Key: NG (no growth) - (no pink color) +(faint pink color) ++(obvious pink color) +++ (dark pink color)

#### Gene *34*; Score *0*

Images taken after 4 days at 37 ºC

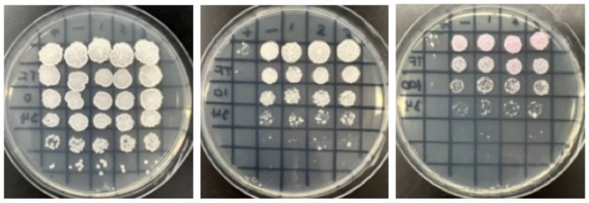


0 ng/ml aTc — 10 ng/ml aTc — 100 ng/ml aTc

| Lane | Gene ID | Plasmid name | Gene name | Toxic/Non-toxic | Colony color on 100 ng/ml aTc plate* |
|---|---|---|---|---|---|
| + Toxic control | -- | pExTra02 | Fruitloop *52* | Toxic | - |
| - Non-toxic control | -- | pExTra03 | Fruitloop *52* mutant | Non-toxic | + |
| 1 | -- | pExTra-Girr34 | Girr *34* replicate 1 | Non-toxic | ++ |
| 2 | -- | pExTra-Girr34 | Girr *34* replicate 2 | Non-toxic | ++ |
| 3 | -- | pExTra-Girr34 | Girr *34* replicate 3 | Non-toxic | ++ |

*Key: NG (no growth) - (no pink color) +(faint pink color) ++(obvious pink color) +++ (dark pink color)

#### Gene *35*; Score *3*

Images taken after 5 days at 37 ºC

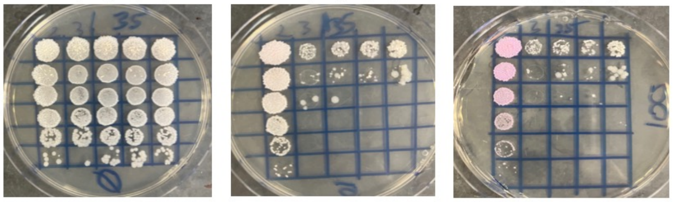


0 ng/ml aTc — 10 ng/ml aTc — 100 ng/ml aTc

| Lane | Gene ID | Plasmid name | Gene name | Toxic/Non-toxic | Colony color on 100 ng/ml aTc plate* |
|---|---|---|---|---|---|
| - Non-toxic control | -- | pExTra03 | Fruitloop 52 mutant | Non-Toxic | +++ |
| + Toxic control | -- | pExTra02 | Fruitloop *52* | Toxic | - |
| 1 | -- | pExTra-Girr35 | Girr *35* replicate 1 | Toxic | - |
| 2 | -- | pExTra-Girr35 | Girr *35* replicate 2 | Toxic | - |
| 3 | -- | pExTra-Girr35 | Girr *35* replicate 3 | Toxic | - |

*Key: NG (no growth) - (no pink color) +(faint pink color) ++(obvious pink color) +++ (dark pink color)

#### Gene *36*; Score *3*

Images taken after 4 days at 37 ºC and 3 days at RT

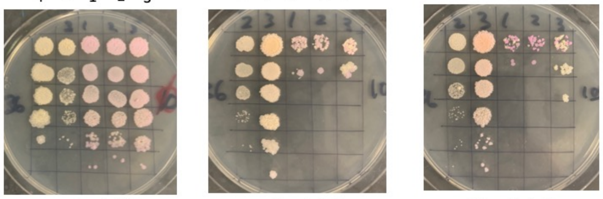


0 ng/ml aTc — 10 ng/ml aTc — 100 ng/ml aTc

| Lane | Gene ID | Plasmid name | Gene name | Toxic/Non-toxic | Colony color on 100 ng/ml aTc plate* |
|---|---|---|---|---|---|
| + Toxic control | -- | pExTra02 | Fruitloop *52* | Toxic | - |
| - Non-toxic control | -- | pExTra03 | Fruitloop *52* mutant | Non-toxic | ++ |
| 1 | -- | pExTra-Girr36 | Girr 36 replicate 1 | Toxic | - |
| 2 | -- | pExTra-Girr36 | Girr 36 replicate 2 | Toxic | - |
| 3 | -- | pExTra-Girr36 | Girr 36 replicate 3 | Toxic | - |

*Key: NG (no growth) - (no pink color) +(faint pink color) ++(obvious pink color) +++ (dark pink color)

#### Gene *37*; Score *2*

mages taken after 4 days at 37 ºC

REPLICATE EXPERIMENT

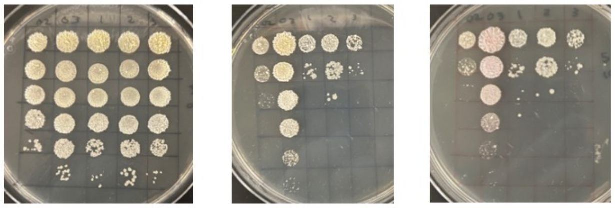


0 ng/ml aTc — 10 ng/ml aTc — 100 ng/ml aTc

| Lane | Gene ID | Plasmid name | Gene name | Toxic/Non-toxic | Colony color on 100 ng/ml aTc plate* |
|---|---|---|---|---|---|
| + Toxic control | -- | pExTra02 | Fruitloop *52* | Toxic | - |
| - Non-toxic control | -- | pExTra03 | Fruitloop *52* mutant | Non-toxic | ++ |
| 1 | -- | pExTra-Girr37 | Girr 37 replicate 1 | Toxic | - |
| 2 | -- | pExTra-Girr73 | Girr 37 replicate 2 | Toxic | - |
| 3 | -- | pExTra-Girr37 | Girr 37 replicate 3 | Toxic | - |

*Key: NG (no growth) - (no pink color) +(faint pink color) ++(obvious pink color) +++ (dark pink color)

#### Gene *38*; Score *0*

Images taken after 4 days at 37 ºC

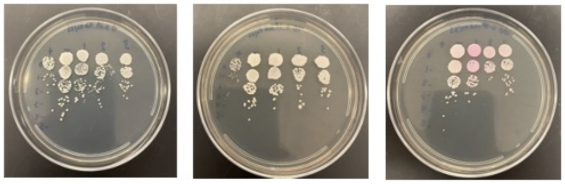


0 ng/ml aTc — 10 ng/ml aTc — 100 ng/ml aTc

| Lane | Gene ID | Plasmid name | Gene name | Toxic/Non-toxic | Colony color on 100 ng/ml aTc plate* |
|---|---|---|---|---|---|
| + Toxic control | -- | pExTra02 | Fruitloop *52* | Toxic | - |
| - Non-toxic control | -- | pExTra03 | Fruitloop *52 mutant* | Non-toxic | + |
| 1 | -- | PExTra-Girr 38 | Girr 38 replicate 1 | Non-toxic | +++ |
| 2 | -- | pExTra-Girr 38 | Girr 38 replicate 2 | Non-toxic | + |
| 3 | -- | pExTra-Girr 38 | Girr 38 replicate 3 | Non-toxic | + |

*Key: NG (no growth) - (no pink color) +(faint pink color) ++(obvious pink color) +++ (dark pink color)

#### Gene *39*; Score *0*

Images taken after 4 days at 37 ºC on 7H11 agar

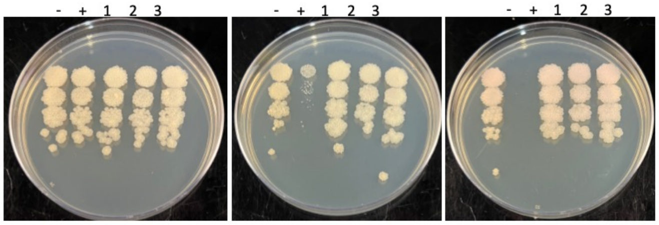


0 ng/ml aTc — 10 ng/ml aTc — 100 ng/ml aTc

| Lane | Plasmid name | Gene name, replicate | Toxic/Non-toxic | Colony color on 100 ng/ml aTc plate* |
|---|---|---|---|---|
| - Non-toxic control | pExTra03 | Fruitloop 52 mutant | Non-toxic | + |
| + Toxic control | pExTra02 | Fruitloop *52* | Toxic | - |
| 1 | pExTra-Girr*39* | Girr *39* replicate 1 | Non-toxic | + |
| 2 | pExTra-Girr*39* | Girr *39* replicate 2 | Non-toxic | + |
| 3 | pExTra-Girr*39* | Girr *39* replicate 3 | Non-toxic | + |

*Key: NG (no growth) - (no pink color) +(faint pink color) ++(obvious pink color) +++ (dark pink color)

#### Gene *40*; Score *0*

Images taken after 4 days at 37 ºC

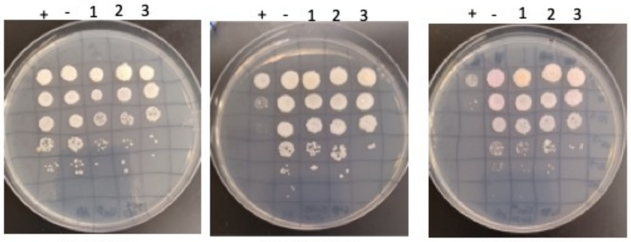


0 ng/ml aTc — 10 ng/ml aTc — 100 ng/ml aTc

| Lane | Gene ID | Plasmid name | Gene name | Toxic/Non-toxic | Colony color on 100 ng/ml aTc plate* |
|---|---|---|---|---|---|
| + Toxic control | -- | pExTra02 | Fruitloop 52 | Toxic | - |
| - Non-toxic control | -- | pExTra03 | Fruitloop *52* mutant | Non-toxic | ++ |
| 1 | -- | pExTra-Girr40 | Girr *40* replicate 1 | Non-toxic | + |
| 2 | -- | pExTra-Girr40 | Girr *40* replicate 2 | Non-toxic | + |
| 3 | -- | pExTra-Girr40 | Girr *40* replicate 3 | Non-toxic | + |

*Key: NG (no growth) - (no pink color) +(faint pink color) ++(obvious pink color) +++ (dark pink color)

#### Gene *41*; Score *0*

Images taken after 4 days at 37 ºC

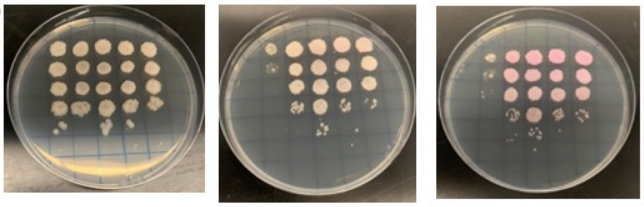


0 ng/ml aTc | 10 ng/ml aTc | 100 ng/ml aTc

| Lane | Gene ID | Plasmid name | Gene name | Toxic/Non-toxic | Colony color on 100 ng/ml aTc plate* |
|---|---|---|---|---|---|
| + Toxic control | -- | pExTra02 | Fruitloop *52* | Toxic | - |
| - Non-toxic control | -- | pExTra03 | Fruitloop *52* mutant | Non-toxic | ++ |
| 1 | -- | pExTra-Girr41 | Girr 41 replicate 1 | Non-toxic | ++ |
| 2 | -- | pExTra-Girr41 | Girr 41 replicate 2 | Non-toxic | ++ |
| 3 | -- | pExTra-Girr41 | Girr 41 replicate 3 | Non-toxic | ++ |

*Key: NG (no growth) - (no pink color) +(faint pink color) ++(obvious pink color) +++ (dark pink color)

#### Gene *42*; Score *0*

Images taken after 5 days at 37 ºC

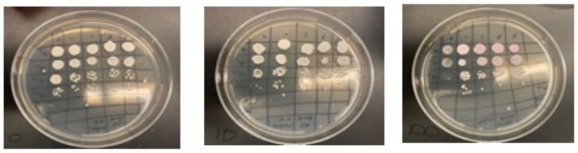


0 ng/ml aTc | 10 ng/ml aTc | 100 ng/ml aTc

| Lane | Gene ID | Plasmid name | Gene name | Toxic/Non-toxic | Colony color on 100 ng/ml aTc plate* |
|---|---|---|---|---|---|
| + Toxic control | -- | pExTra02 | Fruitloop *52* | Toxic | - |
| - Non-toxic control | -- | pExTra03 | Fruitloop *52 mutant* | Non-toxic | + |
| 1 | — | pExTra-GIRR42 | GIRR42 replicate 1 | Non-toxic | ++ |
| 2 | — | pExTra-GIRR42 | GIRR42 replicate 2 | Non-toxic | ++ |
| 3 | — | pExTra-GIRR42 | GIRR42 replicate 3 | Non-toxic | ++ |

*Key: NG (no growth) - (no pink color) +(faint pink color) ++(obvious pink color) +++ (dark pink color)

#### Gene *43*; Score *0*

Images taken after 4 days at 37 ºC

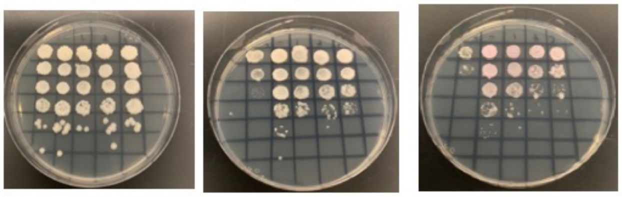


0 ng/ml aTc | 10 ng/ml aTc | 100 ng/ml aTc

| Lane | Gene ID | Plasmid name | Gene name | Toxic/Non-toxic | Colony color on 100 ng/ml aTc plate* |
|---|---|---|---|---|---|
| + Toxic control | -- | pExTra02 | Fruitloop *52* | Toxic | - |
| - Non-toxic control | -- | pExTra03 | Fruitloop *52* mutant | Non-toxic | ++ |
| 1 | -- | pExTra-Girr43 | Girr 43 replicate 1 | Non-toxic | + |
| 2 | -- | pExTra-Girr43 | Girr 43 replicate 2 | Non-toxic | + |
| 3 | -- | pExTra-Girr43 | Girr 43 replicate 3 | Non-toxic | + |

*Key: NG (no growth) - (no pink color) +(faint pink color) ++(obvious pink color) +++ (dark pink color)

#### Gene *44*; Score *2*

Images taken after 4 days at 37 ºC

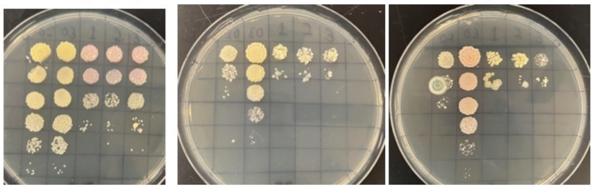


0 ng/ml aTc | 10 ng/ml aTc | 100 ng/ml aTc

| Lane | Gene ID | Plasmid name | Gene name | Toxic/Non-toxic | Colony color on 100 ng/ml aTc plate* |
|---|---|---|---|---|---|
| + Toxic control | -- | pExTra02 | Fruitloop *52* | Toxic | - |
| - Non-toxic control | -- | pExTra03 | Fruitloop *52* mutant | Non-toxic | ++ |
| 1 | -- | pExTra-Girr44 | Girr 44 replicate 1 | Toxic | - |
| 2 | -- | pExTra-Girr44 | Girr 44 replicate 2 | Toxic | - |
| 3 | -- | pExTra-Girr44 | Girr 44 replicate 3 | Toxic | - |

*Key: NG (no growth) - (no pink color) +(faint pink color) ++(obvious pink color) +++ (dark pink color)

#### Gene *45*; Score *0*

Images taken after 4 days at 37 ºC

REPLCATE EXPERIMENT 2023

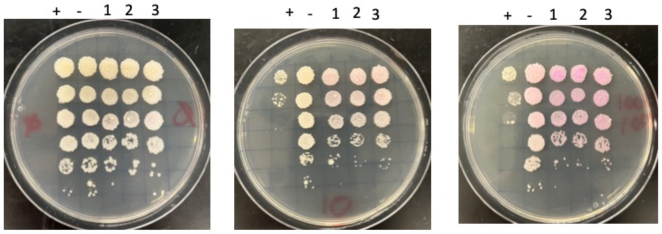


0 ng/ml aTc | 10 ng/ml aTc | 100 ng/ml aTc

| Lane | Gene ID | Plasmid name | Gene name | Toxic/Non-toxic | Colony color on 100 ng/ml aTc plate* |
|---|---|---|---|---|---|
| + Toxic control | -- | pExTra02 | *Fruitloop 52* | Toxic | - |
| - Non-toxic control | -- | pExTra03 | *Fruitloop 52 mutant* | Non-toxic | ++ |
| 1 | -- | pExTra-Girr45 | *Girr 45 replicate 1* | Non-toxic | +++ |
| 2 | -- | pExTra-Girr45 | *Girr 45 replicate 2* | Non-toxic | +++ |
| 3 | -- | pExTra-Girr45 | *Girr 45 replicate 3* | Non-toxic | +++ |

*Key: NG (no growth) - (no pink color) +(faint pink color) ++(obvious pink color) +++ (dark pink color)

#### Gene *46*; Score *3*

Images taken after 5 days at 37 ºC

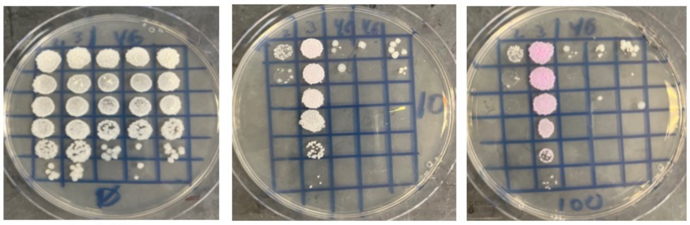


0 ng/ml aTc | 10 ng/ml aTc | 100 ng/ml aTc

| Lane | Gene ID | Plasmid name | Gene name | Toxic/Non-toxic | Colony color on 100 ng/ml aTc plate* |
|---|---|---|---|---|---|
| + Toxic control | -- | pExTra02 | Fruitloop *52* | Toxic | - |
| - Non-toxic control | -- | pExTra03 | Fruitloop *52* mutant | Non-toxic | +++ |
| 1 | -- | pExTra-Girr46 | Girr *46* replicate 1 | Toxic | - |
| 2 | -- | pExTra-Girr46 | Girr *46* replicate 2 | Toxic | - |
| 3 | -- | pExTra-Girr46 | Girr *46* replicate 3 | Toxic | - |

*Key: NG (no growth) - (no pink color) +(faint pink color) ++(obvious pink color) +++ (dark pink color)

#### Gene *47*; Score *0*

Images taken after 4 days at 37 ºC

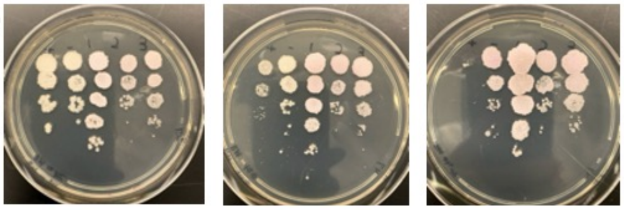


0 ng/ml aTc | 10 ng/ml aTc | 100 ng/ml aTc

| Lane | Gene ID | Plasmid name | Gene name | Toxic/Non-toxic | Colony color on 100 ng/ml aTc plate* |
|---|---|---|---|---|---|
| + Toxic control | -- | pExTra02 | Fruitloop *52* | Toxic | - |
| - Non-toxic control | -- | pExTra03 | Fruitloop *52 mutant* | Non-toxic | + |
| 1 | -- | pExTra-Girr 47 | Girr *47* replicate 1 | Non-toxic | ++ |
| 2 | -- | pExTra-Girr 47 | *Girr 47* replicate 2 | Non-toxic | ++ |
| 3 | -- | pExTra-Girr 47 | *Girr 47* replicate 3 | Non-toxic | ++ |

*Key: NG (no growth) - (no pink color) +(faint pink color) ++(obvious pink color) +++ (dark pink color)

#### Gene *48*; Score *3*

Images taken after 3 days at 37 ºC

REPLCATE EXPERIMENT

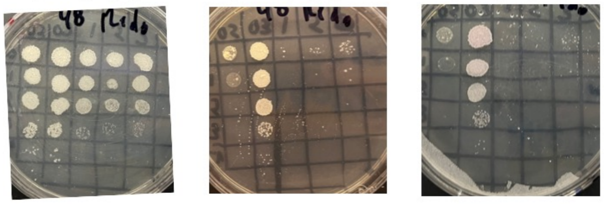


0 ng/ml aTc | 10 ng/ml aTc | 100 ng/ml aTc

| Lane | Gene ID | Plasmid name | Gene name | Toxic/Non-toxic | Colony color on 100 ng/ml aTc plate* |
|---|---|---|---|---|---|
| + Toxic control | | pExTra02 | Fruitloop *52* | Toxic | - |
| - Non-toxic control | | pExTra03 | Fruitloop *52* mutant | Non-toxic | + |
| 1 | | pExTra-Girr48 | Girr *48* replicate 1 | Toxic | NG |
| 2 | | pExTra-Girr48 | Girr *48* replicate 2 | Toxic | NG |
| 3 | | pExTra-Girr48 | Girr *48* replicate 3 | Toxic | NG |

*Key: NG (no growth) - (no pink color) +(faint pink color) ++(obvious pink color) +++ (dark pink color)

Images taken after 5 days at 37 ºC

#### Gene *49*; Score *0*

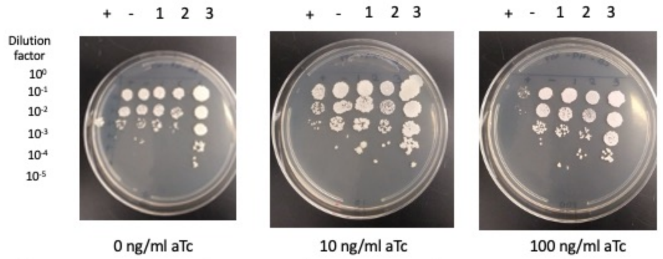


0 ng/ml aTc | 10 ng/ml aTc | 100 ng/ml aTc

| Lane | Gene ID | Plasmid name | Gene name | Toxic/Non-toxic | Colony color on 100 ng/ml aTc plate* |
|---|---|---|---|---|---|
| + Toxic control | -- | pExTra02 | Fruitloop *52* | Toxic | - |
| - Non-toxic control | -- | pExTra03 | Fruitloop *52 mutant* | Non-toxic | + |
| 1 | -- | pExTra-Girr49 | Girr *49* replicate 1 | Non-toxic | ++ |
| 2 | -- | pExTra-Girr49 | Girr *49* replicate 2 | Non-toxic | + |
| 3 | -- | pExTra-Girr49 | Girr *49* replicate 3 | Non-toxic | ++ |

*Key: NG (no growth) - (no pink color) +(faint pink color) ++(obvious pink color) +++ (dark pink color)

Images taken after 4 days at 37 ºC and 3 days at 4ºC

#### Gene *50*; Score *0*

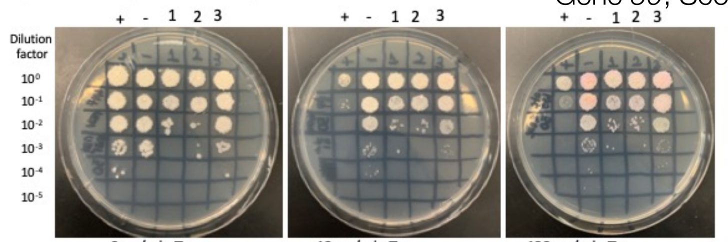


0 ng/ml aTc | 10 ng/ml aTc | 100 ng/ml aTc

| Lane | Gene ID | Plasmid name | Gene name | Toxic/Non-toxic | Colony color on 100 ng/ml aTc plate* |
|---|---|---|---|---|---|
| + Toxic control | -- | pExTra02 | Fruitloop *52* | Toxic | - |
| - Non-toxic control | -- | pExTra03 | Fruitloop *52* mutant | Non-toxic | ++ |
| 1 | -- | pExTra-Girr50 | Girr *50* replicate 1 | Non-toxic | + |
| 2 | -- | pExTra-Girr50 | Girr *50* replicate 2 | Non-toxic | + |
| 3 | -- | pExTra-Girr50 | Girr *50* replicate 3 | Non-toxic | ++ |

*Key: NG (no growth) - (no pink color) +(faint pink color) ++(obvious pink color) +++ (dark pink color)

Images taken after 4 days at 37 ºC FIRST EXPERIMENT

#### Gene *51*; Score *3*

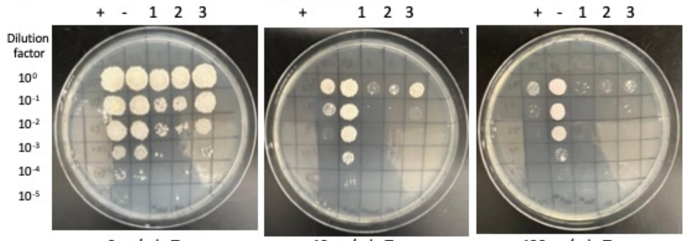


0 ng/ml aTc | 10 ng/ml aTc | 100 ng/ml aTc

| Lane | Gene ID | Plasmid name | Gene name | Toxic/Non-toxic | Colony color on 100 ng/ml aTc plate* |
|---|---|---|---|---|---|
| + Toxic control | -- | pExTra02 | Fruitloop *52* | Toxic | - |
| - Non-toxic control | -- | pExTra03 | Fruitloop *52* mutant | Non-toxic | ++ |
| 1 | 60329186 | pExTra-Girr51 | Girr *51* replicate 1 | Toxic | NG |
| 2 | 60329186 | pExTra-Girr51 | Girr *51* replicate 2 | Toxic | NG |
| 3 | 60329186 | pExTra-Girr51 | Girr *51* replicate 3 | Toxic | NG |

*Key: NG (no growth) - (no pink color) +(faint pink color) ++(obvious pink color) +++ (dark pink color)

Images taken after 5 days at 37 ºC Second Experiment

#### Gene *52*; Score *2*

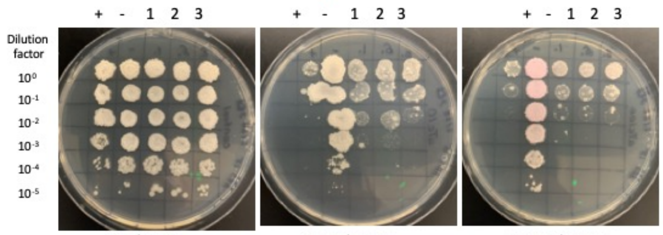


0 ng/ml aTc | 10 ng/ml aTc | 100 ng/ml aTc

| Lane | Gene ID | Plasmid name | Gene name | Toxic/Non-toxic | Colony color on 100 ng/ml aTc plate* |
|---|---|---|---|---|---|
| + Toxic control | -- | pExTra02 | Fruitloop *52* | Toxic | - |
| - Non-toxic control | -- | pExTra03 | Fruitloop *52* mutant | Non-toxic | ++ |
| 1 | -- | pExTra-Girr52 | Girr *52* replicate 1 | Toxic | + |
| 2 | -- | pExTra-Girr52 | Girr 52 replicate 2 | Toxic | + |
| 3 | -- | pExTra-Girr52 | Girr *52* replicate 3 | Toxic | + |

*Key: NG (no growth) - (no pink color) +(faint pink color) ++(obvious pink color) +++ (dark pink color)

Images taken after 4 days at 37 ºC REPLCATE EXPERIMENT

#### Gene *53*; Score *0*

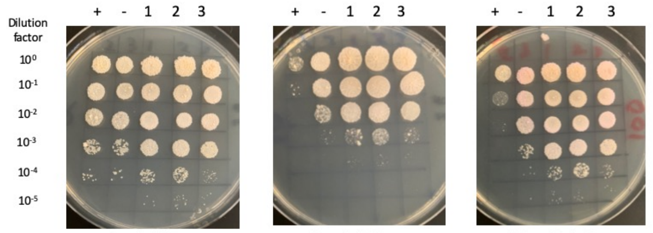


0 ng/ml aTc | 10 ng/ml aTc | 100 ng/ml aTc

| Lane | Gene ID | Plasmid name | Gene name | Toxic/Non-toxic | Colony color on 100 ng/ml aTc plate* |
|---|---|---|---|---|---|
| + Toxic control | -- | pExTra02 | Fruitloop *52* | Toxic | - |
| - Non-toxic control | -- | pExTra03 | Fruitloop *52* mutant | Non-toxic | ++ |
| 1 | -- | pExTra-Girr53 | Girr *53* replicate 1 | Non-toxic | + |
| 2 | -- | pExTra-Girr53 | Girr *53* replicate 2 | Non-toxic | + |
| 3 | -- | pExTra-Girr53 | Girr *53* replicate 3 | Non-toxic | + |

*Key: NG (no growth) - (no pink color) +(faint pink color) ++(obvious pink color) +++ (dark pink color)

Images taken after 4 days at 37 ºC and 3 days at RT

#### Gene *54*; Score *2*

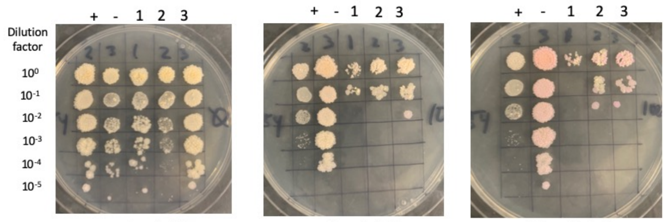


0 ng/ml aTc | 10 ng/ml aTc | 100 ng/ml aTc

| Lane | Gene ID | Plasmid name | Gene name | Toxic/Non-toxic | Colony color on 100 ng/ml aTc plate* |
|---|---|---|---|---|---|
| + Toxic control | -- | pExTra02 | Fruitloop *52* | Toxic | - |
| - Non-toxic control | -- | pExTra03 | Fruitloop *52* mutant | Non-toxic | ++ |
| 1 | -- | pExTra-Girr54 | Girr 54 replicate 1 | Toxic | - |
| 2 | -- | pExTra-Girr54 | Girr 54 replicate 2 | Toxic | - |
| 3 | -- | pExTra-Girr54 | Girr 54 replicate 3 | Toxic | - |

*Key: NG (no growth) - (no pink color) +(faint pink color) ++(obvious pink color) +++ (dark pink color)

Images taken after 4 days at 37 ºC

#### Gene *55*; Score *0*

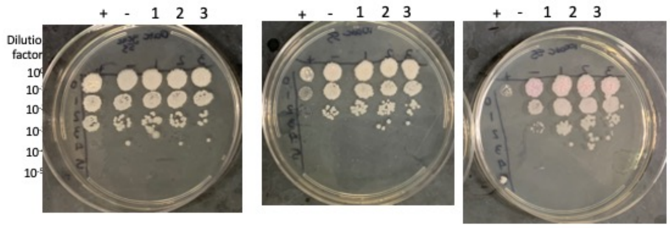


0 ng/ml aTc | 10 ng/ml aTc | 100 ng/ml aTc

| Lane | Gene ID | Plasmid name | Gene name | Toxic/Non-toxic | Colony color on 100 ng/ml aTc plate* |
|---|---|---|---|---|---|
| + Toxic control | -- | pExTra02 | Fruitloop *52* | Toxic | - |
| - Non-toxic control | -- | pExTra03 | Fruitloop *52 mutant* | Non-toxic | + |
| 1 | -- | pExTra-Girr 55 | Girr 55 replicate 1 | Non-toxic | ++ |
| 2 | -- | pExTra-Girr 55 | Girr 55 replicate 2 | Non-toxic | ++ |
| 3 | -- | pExTra-Girr 55 | *Girr 55* replicate 3 | Non-toxic | ++ |

*Key: NG (no growth) - (no pink color) +(faint pink color) ++(obvious pink color) +++ (dark pink color)

Images taken after 5 days at 37 ºC First replicate

#### Gene *56*; Score *1*

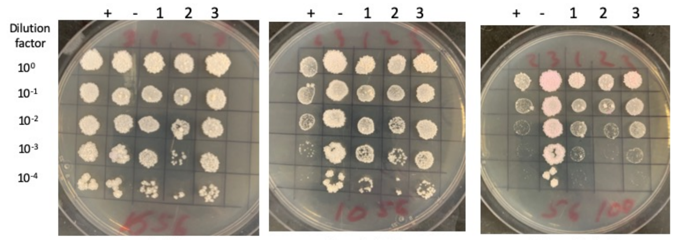


0 ng/ml aTc | 10 ng/ml aTc | 100 ng/ml aTc

| Lane | Gene ID | Plasmid name | Gene name | Toxic/Non-toxic | Colony color on 100 ng/ml aTc plate* |
|---|---|---|---|---|---|
| + Toxic control | -- | pExTra02 | Fruitloop *52* | Toxic | - |
| - Non-toxic control | -- | pExTra03 | Fruitloop *52* mutant | Non-toxic | ++ |
| 1 | -- | pExTra-Girr56 | Girr *56* replicate 1 | Toxic | + |
| 2 | -- | pExTra-Girr56 | Girr *56* replicate 2 | Toxic | + |
| 3 | -- | pExTra-Girr56 | Girr *56* replicate 3 | Toxic | + |

*Key: NG (no growth) - (no pink color) +(faint pink color) ++(obvious pink color) +++ (dark pink color)

#### Gene *57*; Score *3*

Images taken after 5 days at 37 ºC

\+ - 1 2 3 | + - 1 2 3 | + - 1 2 3

Dilution factor: $10^0$, $10^{-1}$, $10^{-2}$, $10^{-3}$, $10^{-4}$, $10^{-5}$

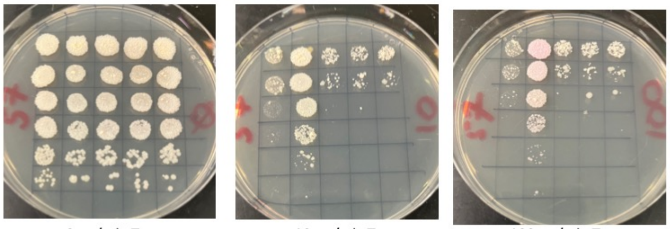

0 ng/ml aTc | 10 ng/ml aTc | 100 ng/ml aTc

| Lane | Gene ID | Plasmid name | Gene name | Toxic/Non-toxic | Colony color on 100 ng/ml aTc plate* |
|---|---|---|---|---|---|
| + Toxic control | -- | pExTra02 | Fruitloop *52* | Toxic | - |
| - Non-toxic control | -- | pExTra03 | Fruitloop *52* mutant | Non-toxic | + |
| 1 | -- | pExTra-Girr57 | Girr 57 replicate 1 | Toxic | - |
| 2 | -- | pExTra-Girr57 | Girr 57 replicate 2 | Toxic | - |
| 3 | -- | pExTra-Girr57 | Girr 57 replicate 3 | Toxic | - |

*Key: NG (no growth) - (no pink color) +(faint pink color) ++(obvious pink color) +++ (dark pink color)

#### Gene *58*; Score *0*

Images taken after 4 days at 37 ºC

\+ - 1 2 3 | + - 1 2 3 | + - 1 2 3

Dilution factor: $10^0$, $10^{-1}$, $10^{-2}$, $10^{-3}$, $10^{-4}$, $10^{-5}$

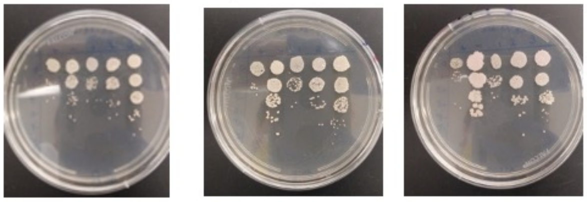

0 ng/ml aTc | 10 ng/ml aTc | 100 ng/ml aTc

| Lane | Gene ID | Plasmid name | Gene name | Toxic/Non-toxic | Colony color on 100 ng/ml aTc plate* |
|---|---|---|---|---|---|
| + Toxic control | -- | pExTra02 | Fruitloop *52* | Toxic | - |
| - Non-toxic control | -- | pExTra03 | Fruitloop *52 mutant* | Non-toxic | + |
| 1 | -- | pExTra-Girr58 | Girr *58* replicate 1 | Non-toxic | + |
| 2 | -- | pExTra-Girr58 | Girr *58* replicate 2 | Non-toxic | + |
| 3 | -- | pExTra-Girr58 | Girr *58* replicate 3 | Non-toxic | ++ |

*Key: NG (no growth) - (no pink color) +(faint pink color) ++(obvious pink color) +++ (dark pink color)

#### Gene *59*; Score *0*

Images taken after 5 days at 37 ºC

\+ - 1 2 3 | + - 1 2 3 | + - 1 2 3

Dilution factor: $10^0$, $10^{-1}$, $10^{-2}$, $10^{-3}$, $10^{-4}$, $10^{-5}$

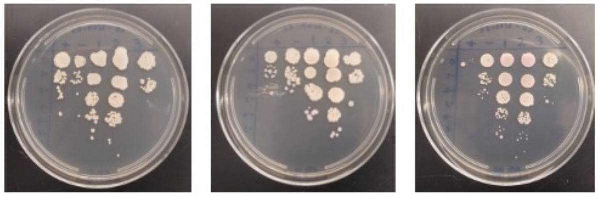

0 ng/ml aTc | 10 ng/ml aTc | 100 ng/ml aTc

| Lane | Gene ID | Plasmid name | Gene name | Toxic/Non-toxic | Colony color on 100 ng/ml aTc plate* |
|---|---|---|---|---|---|
| + Toxic control | -- | pExTra02 | Fruitloop *52* | Toxic | - |
| - Non-toxic control | -- | pExTra03 | Fruitloop *52 mutant* | Non-toxic | + |
| 1 | -- | pExTra-Girr59 | Girr *59* replicate 1 | Non-toxic | + |
| 2 | -- | pExTra-Girr59 | Girr *59* replicate 2 | Non-toxic | ++ |
| 3 | -- | pExTra-Girr59 | Girr *59* replicate 3 | Non-toxic | - |

*Key: NG (no growth) - (no pink color) +(faint pink color) ++(obvious pink color) +++ (dark pink color)

#### Gene *60*; Score *3*

Images taken after 4 days at 37 ºC

REPLCATE EXPERIMENT 2023

\+ - 1 2 3 | + - 1 2 3 | + - 1 2 3

Dilution factor: $10^0$, $10^{-1}$, $10^{-2}$, $10^{-3}$, $10^{-4}$, $10^{-5}$

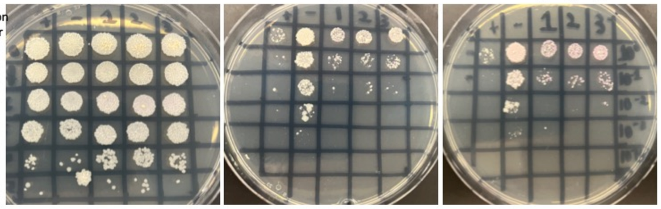

0 ng/ml aTc | 10 ng/ml aTc | 100 ng/ml aTc

| Lane | Gene ID | Plasmid name | Gene name | Toxic/Non-toxic | Colony color on 100 ng/ml aTc plate* |
|---|---|---|---|---|---|
| + Toxic control | -- | pExTra02 | Fruitloop *52* | Toxic | - |
| - Non-toxic control | -- | pExTra03 | Fruitloop *52* mutant | Non-toxic | - |
| 1 | -- | pExTra-Girr60 | Girr *60* replicate 1 | Toxic | + |
| 2 | -- | pExTra-Girr60 | Girr *60* replicate 2 | Toxic | + |
| 3 | -- | pExTra-Girr60 | Girr *60* replicate 3 | Toxic | + |

*Key: NG (no growth) - (no pink color) +(faint pink color) ++(obvious pink color) +++ (dark pink color)

#### Gene *61*; Score *0*

Images taken after 4 days at 37 ºC

\+ - 1 2 3 | + - 1 2 3 | + - 1 2 3

Dilution factor: $10^0$, $10^{-1}$, $10^{-2}$, $10^{-3}$, $10^{-4}$, $10^{-5}$

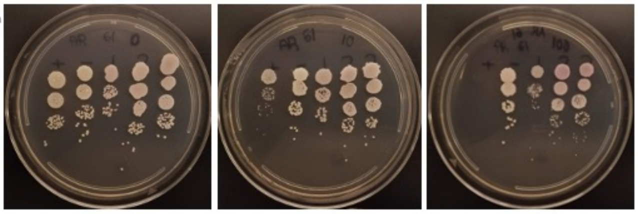

0 ng/ml aTc | 10 ng/ml aTc | 100 ng/ml aTc

| Lane | Gene ID | Plasmid name | Gene name | Toxic/Non-toxic | Colony color on 100 ng/ml aTc plate* |
|---|---|---|---|---|---|
| + Toxic control | -- | pExTra02 | Fruitloop *52* | Toxic | - |
| - Non-toxic control | -- | pExTra03 | Fruitloop *52 mutant* | Non-toxic | + |
| 1 | 131436 | pExTra-Girr 61 | Girr *61* replicate 1 | Non-toxic | + |
| 2 | 131436 | pExTra-Girr 61 | Girr *61* replicate 2 | Non-toxic | ++ |
| 3 | 131436 | pExTra-Girr 61 | Girr *61* replicate 3 | Non-toxic | ++ |

*Key: NG (no growth) - (no pink color) +(faint pink color) ++(obvious pink color) +++ (dark pink color)

#### Gene *62*; Score *1*

Images taken after 3 days at 37 ºC on 7H11 agar

\- + 1 2 3 | - + 1 2 3 | - + 1 2 3

Dilution factor: $10^0$, $10^{-1}$, $10^{-2}$, $10^{-3}$, $10^{-4}$, $10^{-5}$

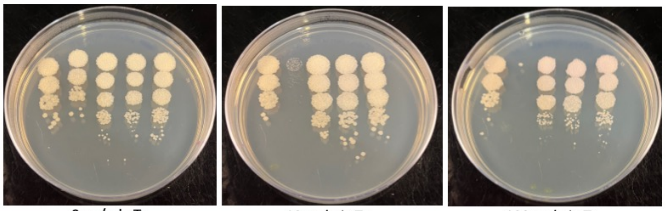

0 ng/ml aTc | 10 ng/ml aTc | 100 ng/ml aTc

| Lane | Plasmid name | Gene name, replicate | Toxic/Non-toxic | Colony color on 100 ng/ml aTc plate* |
|---|---|---|---|---|
| - Non-toxic control | pExTra03 | Fruitloop *52* mutant | Non-toxic | + |
| + Toxic control | pExTra02 | Fruitloop *52* | Toxic | - |
| 1 | pExTra-Girr*62* | Girr *62* replicate 1 | Toxic | + |
| 2 | pExTra-Girr*62* | Girr *62* replicate 2 | Toxic | + |
| 3 | pExTra-Girr*62* | Girr *62* replicate 3 | Toxic | + |

*Key: NG (no growth) - (no pink color) +(faint pink color) ++(obvious pink color) +++ (dark pink color)

#### Gene *63*; Score *1*

Images taken after 4 days at 37 ºC

REPLICATE EXPERIMENT

\+ - 1 2 3 | + - 1 2 3 | + - 1 2 3

Dilution factor: $10^0$, $10^{-1}$, $10^{-2}$, $10^{-3}$, $10^{-4}$, $10^{-5}$

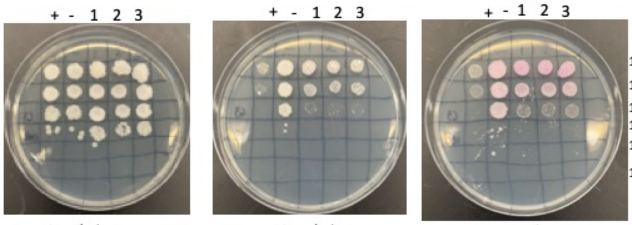

$10^0$, $10^{-1}$, $10^{-2}$, $10^{-3}$, $10^{-4}$, $10^{-5}$

0 ng/ml aTc | 10 ng/ml aTc | 100 ng/ml aTc

| Lane | Gene ID | Plasmid name | Gene name | Toxic/Non-toxic | Colony color on 100 ng/ml aTc plate* |
|---|---|---|---|---|---|
| + Toxic control | -- | pExTra02 | Fruitloop *52* | Toxic | - |
| - Non-toxic control | -- | pExTra03 | Fruitloop *52* mutant | Non-toxic | + |
| 1 | f0329198 | pExTra-Girr63 | Girr *63* replicate 1 | Toxic | ++ |
| 2 | f0329198 | pExTra-Girr63 | Girr *63* replicate 2 | Toxic | ++ |
| 3 | f0329198 | pExTra-Girr63 | Girr *63* replicate 3 | Toxic | +++ |

*Key: NG (no growth) - (no pink color) +(faint pink color) ++(obvious pink color) +++ (dark pink color)

#### Gene *64*; Score *1*

Images taken after 4 days at 37 ºC

REPLCATE EXPERIMENT 2023

\+ - 1 2 3 | + - 1 2 3 | + - 1 2 3

Dilution factor: $10^0$, $10^{-1}$, $10^{-2}$, $10^{-3}$, $10^{-4}$, $10^{-5}$

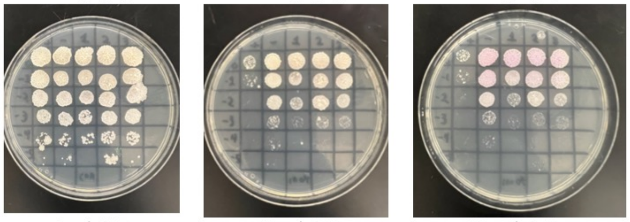

0 ng/ml aTc | 10 ng/ml aTc | 100 ng/ml aTc

| Lane | Gene ID | Plasmid name | Gene name | Toxic/Non-toxic | Colony color on 100 ng/ml aTc plate* |
|---|---|---|---|---|---|
| + Toxic control | -- | pExTra02 | Fruitloop *52* | Toxic | - |
| - Non-toxic control | -- | pExTra03 | Fruitloop *52 I/S* | Non-toxic | ++ |
| 1 | NKF | pExTra-Girr64 | Girr *64* replicate 1 | Toxic | ++ |
| 2 | NKF | pExTra-Girr64 | Girr *64* replicate 2 | Toxic | ++ |
| 3 | NKF | pExTra-Girr64 | Girr *64* replicate 3 | Toxic | ++ |

*Key: NG (no growth) - (no pink color) +(faint pink color) ++(obvious pink color) +++ (dark pink color)

#### Gene *65*; Score *2*

Images taken after 4 days at 37 ºC

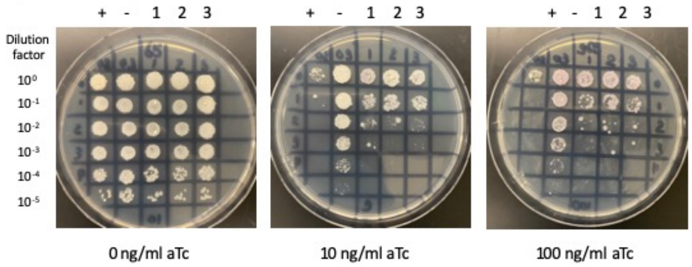


| Lane | Gene ID | Plasmid name | Gene name | Toxic/Non-toxic | Colony color on 100 ng/ml aTc plate* |
|---|---|---|---|---|---|
| + Toxic control | -- | pExTra02 | Fruitloop *52* | Toxic | - |
| - Non-toxic control | -- | pExTra03 | Fruitloop *52* mutant | Non-toxic | + |
| 1 | -- | pExTra-Girr65 | Girr *65* replicate 1 | toxic | + |
| 2 | -- | pExTra-Girr65 | Girr *65* replicate 2 | toxic | + |
| 3 | -- | pExTra-Girr65 | Girr *65* replicate 3 | toxic | + |

*Key: NG (no growth) - (no pink color) +(faint pink color) ++(obvious pink color) +++ (dark pink color)

#### Gene *66*; Score *0*

Images taken after 3 days at 37 ºC

REPLCATE EXPERIMENT

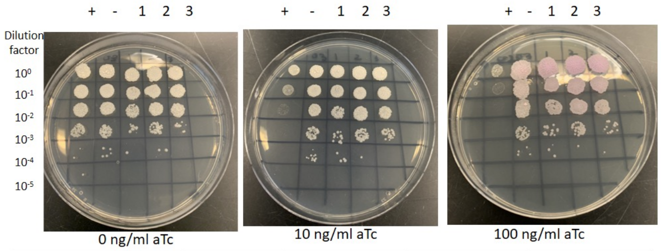


| Lane | Gene ID | Plasmid name | Gene name | Toxic/Non-toxic | Colony color on 100 ng/ml aTc plate* |
|---|---|---|---|---|---|
| + Toxic control | -- | pExTra02 | Fruitloop *52* | Toxic | - |
| - Non-toxic control | -- | pExTra03 | Fruitloop *52* mutant | Non-toxic | + |
| 1 | -- | pExTra-Girr66 | Girr 66 replicate 1 | Non-toxic | ++ |
| 2 | -- | pExTra-Girr66 | Girr 66 replicate 2 | Non-toxic | ++ |
| 3 | -- | pExTra-Girr66 | Girr 66 replicate 3 | Non-toxic | ++ |

*Key: NG (no growth) - (no pink color) +(faint pink color) ++(obvious pink color) +++ (dark pink color)

#### Gene *67*; Score *0*

Images taken after 4 days at 37 ºC

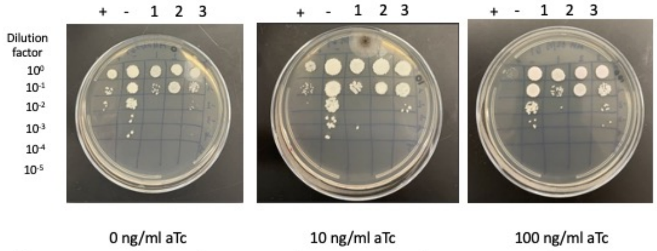


| Lane | Gene ID | Plasmid name | Gene name | Toxic/Non-toxic | Colony color on 100 ng/ml aTc plate* |
|---|---|---|---|---|---|
| + Toxic control | -- | pExTra02 | Fruitloop *52* | Toxic | - |
| - Non-toxic control | -- | pExTra03 | Fruitloop *52 mutant* | Non-toxic | + |
| 1 | -- | PExTra-Girr67 | Girr 67 replicate 1 | Non-toxic | - |
| 2 | -- | pExTra-Girr67 | Girr 67 replicate 2 | Non-toxic | - |
| 3 | -- | pExTra-Girr67 | Girr 67 replicate 3 | Non-toxic | - |

*Key: NG (no growth) - (no pink color) +(faint pink color) ++(obvious pink color) +++ (dark pink color)

#### Gene *68*; Score *0*

Images taken after 4 days at 37ºC

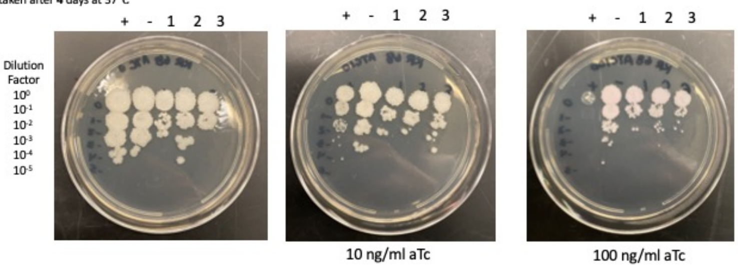


| Lane | Gene ID | Plasmid name | Gene name | Toxic/Non-toxic | Colony color on 100 ng/ml aTc plate* |
|---|---|---|---|---|---|
| + Toxic control | -- | pExTra02 | Fruitloop *52* | Toxic | - |
| - Non-toxic control | -- | pExTra03 | Fruitloop *52 mutant* | Non-toxic | + |
| 1 | — | pExTra-Girr68 | *Girr 68 replicate 1* | Non-toxic | + |
| 2 | — | pExTra-Girr68 | *Girr 68 replicate 2* | Non-toxic | + |
| 3 | — | pExTra-Girr68 | *Girr 68 replicate 3* | Non-toxic | + |

*Key: NG (no growth) - (no pink color) + (faint pink color) ++ (obvious pink color) +++ (dark pink color)

#### Gene *69*; Score *1*

Images taken after 3 days at 37 ºC on 7H11 agar

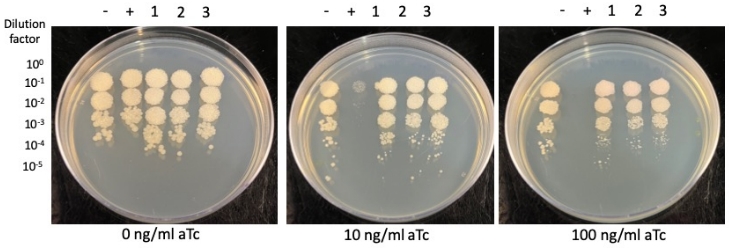


| Lane | Plasmid name | Gene name, replicate | Toxic/Non-toxic | Colony color on 100 ng/ml aTc plate* |
|---|---|---|---|---|
| - Non-toxic control | pExTra03 | Fruitloop *52* mutant | Non-toxic | + |
| + Toxic control | pExTra02 | Fruitloop *52* | Toxic | - |
| 1 | pExTra-Girr*69* | Girr *69* replicate 1 | Toxic | + |
| 2 | pExTra-Girr*69* | Girr *69* replicate 2 | Toxic | + |
| 3 | pExTra-Girr*69* | Girr *69* replicate 3 | Toxic | + |

*Key: NG (no growth) - (no pink color) +(faint pink color) ++(obvious pink color) +++ (dark pink color)

#### Gene *70*; Score *0*

Images taken after 5 days at 37 ºC

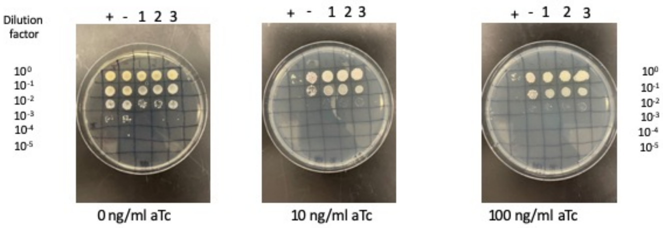


| Lane | Gene ID | Plasmid name | Gene name | Toxic/Non-toxic | Colony color on 100 ng/ml aTc plate* |
|---|---|---|---|---|---|
| + Toxic control | -- | pExTra02 | Fruitloop *52* | Toxic | - |
| - Non-toxic control | -- | pExTra03 | Fruitloop *52* mutant | Non-toxic | + |
| 1 | | pExTra-Girr70 | Girr *70* replicate 1 | Non-Toxic | - |
| 2 | | pExTra-Girr70 | Girr *70* replicate 2 | Non-Toxic | - |
| 3 | | pExTra-Girr70 | Girr *70* replicate 3 | Non-Toxic | - |

*Key: NG (no growth) - (no pink color) +(faint pink color) ++(obvious pink color) +++ (dark pink color)

#### Gene *71*; Score *3*

Images taken after 3 days at 37 ºC

REPLCATE EXPERIMENT

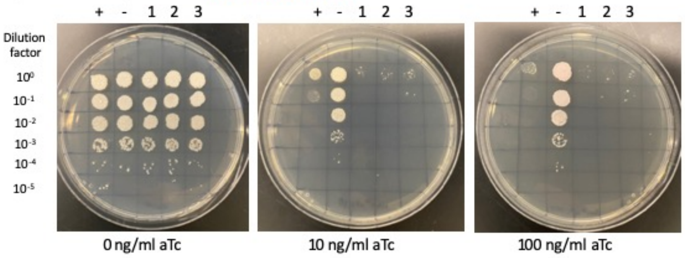


| Lane | Gene ID | Plasmid name | Gene name | Toxic/Non-toxic | Colony color on 100 ng/ml aTc plate* |
|---|---|---|---|---|---|
| + Toxic control | -- | pExTra02 | Fruitloop *52* | Toxic | - |
| - Non-toxic control | -- | pExTra03 | Fruitloop *52* mutant | Non-toxic | + |
| 1 | -- | pExTra-Girr71 | Girr *71* replicate 1 | Toxic | - |
| 2 | -- | pExTra-Girr71 | Girr *71* replicate 2 | Toxic | - |
| 3 | -- | pExTra-Girr71 | Girr *71* replicate 3 | Toxic | - |

*Key: NG (no growth) - (no pink color) +(faint pink color) ++(obvious pink color) +++ (dark pink color)

#### Gene *72*; Score2

Images taken after 4 days at 37 ºC

REPLCATE EXPERIMENT 2023

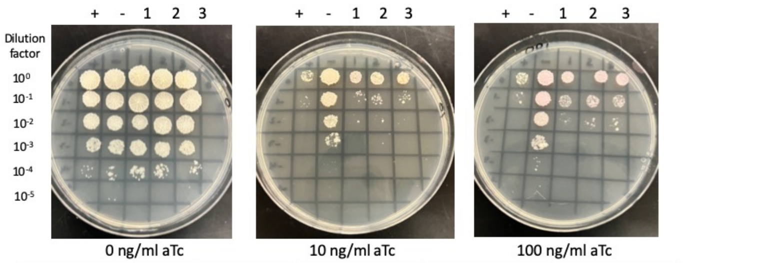


| Lane | Gene ID | Plasmid name | Gene name | Toxic/Non-toxic | Colony color on 100 ng/ml aTc plate* |
|---|---|---|---|---|---|
| + Toxic control | -- | pExTra02 | Fruitloop *52* | Toxic | - |
| - Non-toxic control | -- | pExTra03 | Fruitloop *52* mutant | Non-toxic | ++ |
| 1 | -- | pExTra-Girr72 | Girr *72* replicate 1 | Toxic | + |
| 2 | -- | pExTra-Girr72 | Girr *72* replicate 2 | Toxic | ++ |
| 3 | -- | pExTra-Girr72 | Girr *72* replicate 3 | Toxic | ++ |

*Key: NG (no growth) - (no pink color) +(faint pink color) ++(obvious pink color) +++ (dark pink color)

#### Gene *73*; Score *3*

Images taken after 4 days at 37 ºC

REPLCATE EXPERIMENT 2023

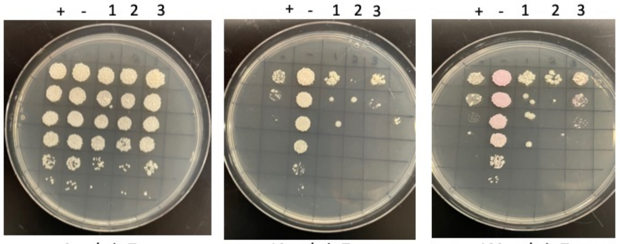


0 ng/ml aTc | 10 ng/ml aTc | 100 ng/ml aTc

| Lane | Gene ID | Plasmid name | Gene name | Toxic/Non-toxic | Colony color on 100 ng/ml aTc plate* |
|---|---|---|---|---|---|
| + Toxic control | -- | pExTra02 | Girr73 | Toxic | - |
| - Non-toxic control | -- | pExTra03 | Girr73 mutant | Non-toxic | ++ |
| 1 | -- | pExTra-Girr73 | Girr *73* replicate 1 | Toxic | - |
| 2 | -- | pExTra-Girr73 | Girr *73* replicate 2 | Toxic | - |
| 3 | -- | pExTra-Girr73 | Girr *73* replicate 3 | Toxic | + |

*Key: NG (no growth) - (no pink color) +(faint pink color) ++(obvious pink color) +++ (dark pink color)

#### Gene *74*; Score *0*

Images taken after 4 days at 37 ºC.

Replicate 2

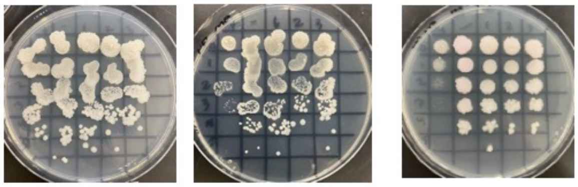


0 ng/ml aTc | 10 ng/ml aTc | 100 ng/ml aTc

| Lane | Gene ID | Plasmid name | Gene name | Toxic/Non-toxic | Colony color on 100 ng/ml aTc plate* |
|---|---|---|---|---|---|
| + Toxic control | -- | pExTra02 | Fruitloop *52* | Toxic | - |
| - Non-toxic control | -- | pExTra03 | Fruitloop *52* mutant | Non-toxic | + |
| 1 | -- | pExTra-Girr74 | Girr 74 replicate 1 | Non-toxic | - |
| 2 | -- | pExTra-Girr74 | Girr 74 replicate 2 | Non-toxic | - |
| 3 | -- | pExTra-Girr74 | Girr 74 replicate 3 | Non-toxic | - |

*Key: NG (no growth) - (no pink color) +(faint pink color) ++(obvious pink color) +++ (dark pink color)

#### Gene *75*; Score *0*

Images taken after 4 days at 37 ºC, and then 3 days at 22 ºC

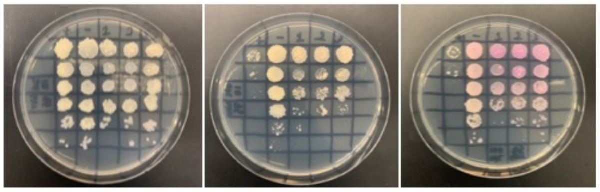


0 ng/ml aTc | 10 ng/ml aTc | 100 ng/ml aTc

| Lane | Gene ID | Plasmid name | Gene name | Toxic/Non-toxic | Colony color on 100 ng/ml aTc plate* |
|---|---|---|---|---|---|
| + Toxic control | -- | pExTra02 | Fruitloop *52* | Toxic | - |
| - Non-toxic control | -- | pExTra03 | Fruitloop *52* mutant | Non-toxic | ++ |
| 1 | -- | pExTra-Girr75 | Girr *75* replicate 1 | Non-toxic | +++ |
| 2 | -- | pExTra-Girr75 | Girr *75* replicate 2 | Non-toxic | +++ |
| 3 | -- | pExTra-Girr75 | Girr *75* replicate 3 | Non-toxic | +++ |

*Key: NG (no growth) - (no pink color) +(faint pink color) ++(obvious pink color) +++ (dark pink color)

#### Gene *76*; Score *0*

Images taken after 4 days at 37 ºC

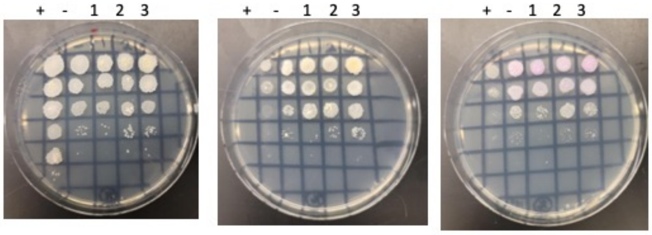


0 ng/ml aTc | 10 ng/ml aTc | 100 ng/ml aTc

| Lane | Gene ID | Plasmid name | Gene name | Toxic/Non-toxic | Colony color on 100 ng/ml aTc plate* |
|---|---|---|---|---|---|
| + Toxic control | 60329210 | pExTra02 | Fruitloop *52* | Toxic | - |
| - Non-toxic control | 60329210 | pExTra03 | Fruitloop *52* mutant | Non-toxic | +++ |
| 1 | 60329210 | pExTra-Girr76 | Girr *76* replicate 1 | Non-toxic | +++ |
| 2 | 60329210 | pExTra-Girr76 | Girr *76* replicate 2 | Non-toxic | +++ |
| 3 | 60329210 | pExTra-Girr76 | Girr *76* replicate 3 | Non-toxic | +++ |

*Key: NG (no growth) - (no pink color) +(faint pink color) ++(obvious pink color) +++ (dark pink color)

#### Gene *77*; Score *0*

Images taken after 4 days at 37 ºC on 7H11 agar

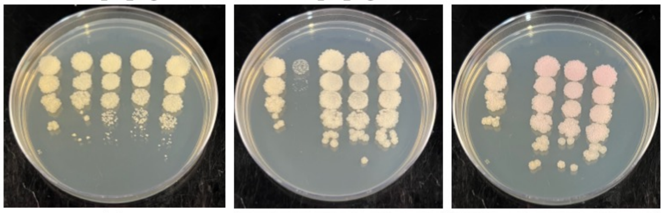


0 ng/ml aTc | 10 ng/ml aTc | 100 ng/ml aTc

| Lane | Plasmid name | Gene name, replicate | Toxic/Non-toxic | Colony color on 100 ng/ml aTc plate* |
|---|---|---|---|---|
| - Non-toxic control | pExTra03 | Fruitloop *52* mutant | Non-toxic | + |
| + Toxic control | pExTra02 | Fruitloop *52* | Toxic | - |
| 1 | pExTra-Girr*77* | Girr *77* replicate 1 | Non-toxic | ++ |
| 2 | pExTra-Girr*77* | Girr *77* replicate 2 | Non-toxic | ++ |
| 3 | pExTra-Girr*77* | Girr *77* replicate 3 | Non-toxic | ++ |

*Key: NG (no growth) - (no pink color) +(faint pink color) ++(obvious pink color) +++ (dark pink color)

#### Gene *78*; Score *0*

Images taken after 4 days at 37 ºC

Replicate experiment

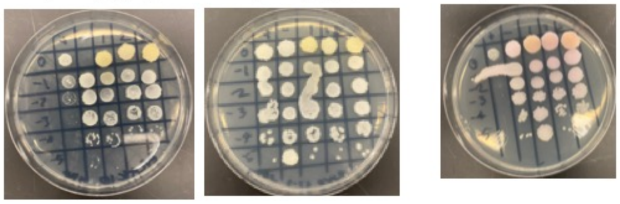


0 ng/ml aTc | 10 ng/ml aTc | 100 ng/ml aTc

| Lane | Gene ID | Plasmid name | Gene name | Toxic/Non-toxic | Colony color on 100 ng/ml aTc plate* |
|---|---|---|---|---|---|
| + Toxic control | -- | pExTra02 | Fruitloop *52* | Toxic | - |
| - Non-toxic control | -- | pExTra03 | Fruitloop *52* mutant | Non-toxic | + |
| 1 | | pExTra-Girr78 | Girr *78* replicate 1 | Non-toxic | ++ |
| 2 | | pExTra-Girr78 | Girr *78* replicate 2 | Non-toxic | ++ |
| 3 | | pExTra-Girr78 | Girr *78* replicate 3 | Non-toxic | +++ |

*Key: NG (no growth) - (no pink color) +(faint pink color) ++(obvious pink color) +++ (dark pink color)

#### Gene *79*; Score *0*

Images taken after 4 days at 37 ºC on 7H11 agar

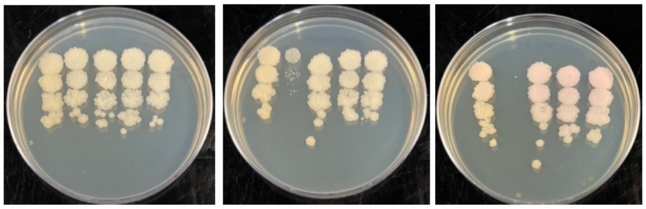


0 ng/ml aTc | 10 ng/ml aTc | 100 ng/ml aTc

| Lane | Plasmid name | Gene name, replicate | Toxic/Non-toxic | Colony color on 100 ng/ml aTc plate* |
|---|---|---|---|---|
| - Non-toxic control | pExTra03 | Fruitloop *52* mutant | Non-toxic | + |
| + Toxic control | pExTra02 | Fruitloop *52* | Toxic | - |
| 1 | pExTra-Girr*79* | Girr *79* replicate 1 | Non-toxic | ++ |
| 2 | pExTra-Girr*79* | Girr *79* replicate 2 | Non-toxic | ++ |
| 3 | pExTra-Girr*79* | Girr *79* replicate 3 | Non-toxic | ++ |

*Key: NG (no growth) - (no pink color) +(faint pink color) ++(obvious pink color) +++ (dark pink color)

#### Gene *80*; Score *0*

Images taken after 5 days at 37 ºC

replicate

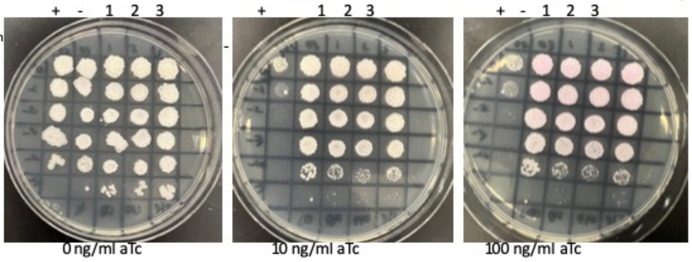


0 ng/ml aTc | 10 ng/ml aTc | 100 ng/ml aTc

| Lane | Gene ID | Plasmid name | Gene name | Toxic/Non-toxic | Colony color on 100 ng/ml aTc plate* |
|---|---|---|---|---|---|
| + Toxic control | -- | pExTra02 | Fruitloop *52* | Toxic | - |
| - Non-toxic control | -- | pExTra03 | Fruitloop *52* mutant | Non-toxic | + |
| 1 | -- | pExTra-Girr80 | Girr 80 replicate 1 | Non-toxic | ++ |
| 2 | -- | pExTra-Girr80 | Girr 80 replicate 2 | Non-toxic | ++ |
| 3 | -- | pExTra-Girr80 | Girr 80 replicate 3 | Non-toxic | ++ |

*Key: NG (no growth) - (no pink color) +(faint pink color) ++(obvious pink color) +++ (dark pink color)

#### Gene *81*; Score *0*

Images taken after 4 days at 37 ºC

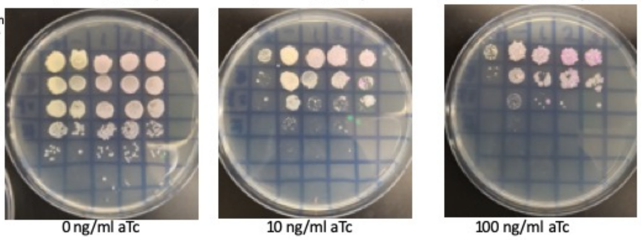


0 ng/ml aTc | 10 ng/ml aTc | 100 ng/ml aTc

| Lane | Gene ID | Plasmid name | Gene name | Toxic/Non-toxic | Colony color on 100 ng/ml aTc plate* |
|---|---|---|---|---|---|
| + Toxic control | | pExTra02 | Fruitloop *52* | Toxic | - |
| - Non-toxic control | | pExTra03 | Fruitloop *52* mutant | Non-toxic | + |
| 1 | | pExTra-Girr81 | Girr *81* replicate 1 | Non-Toxic | ++ |
| 2 | | pExTra-Girr81 | Girr *81* replicate 2 | Non-Toxic | +++ |
| 3 | | pExTra-Girr81 | Girr *81* replicate 3 | Non-Toxic | +++ |

*Key: NG (no growth) - (no pink color) +(faint pink color) ++(obvious pink color) +++ (dark pink color)

#### Gene *82*; Score *0*

Images taken after 4 days at 37 ºC

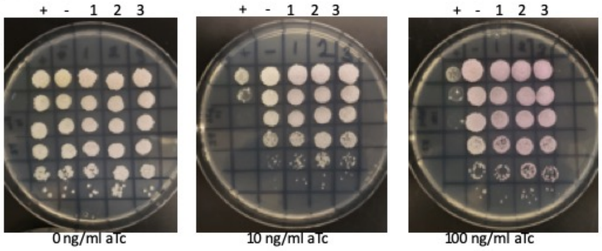


0 ng/ml aTc | 10 ng/ml aTc | 100 ng/ml aTc

| Lane | Gene ID | Plasmid name | Gene name | Toxic/Non-toxic | Colony color on 100 ng/ml aTc plate* |
|---|---|---|---|---|---|
| + Toxic control | | pExTra02 | Fruitloop *52* | Toxic | - |
| - Non-toxic control | | pExTra03 | Fruitloop *52* mutant | Non-toxic | + |
| 1 | | pExTra-Girr82 | Girr *82* replicate 1 | Non-Toxic | ++ |
| 2 | | pExTra-Girr82 | Girr *82* replicate 2 | Non-Toxic | ++ |
| 3 | | pExTra-Girr82 | Girr *82* replicate 3 | Non-Toxic | ++ |

*Key: NG (no growth) - (no pink color) +(faint pink color) ++(obvious pink color) +++ (dark pink color)

#### Gene *83*; Score *0*

Images taken after 4 days at 37 ºC

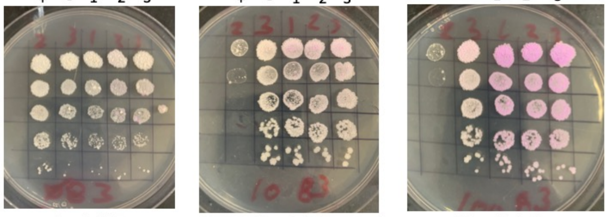


0 ng/ml aTc | 10 ng/ml aTc | 100 ng/ml aTc

| Lane | Gene ID | Plasmid name | Gene name | Toxic/Non-toxic | Colony color on 100 ng/ml aTc plate* |
|---|---|---|---|---|---|
| + Toxic control | -- | pExTra02 | Fruitloop *52* | Toxic | - |
| - Non-toxic control | -- | pExTra03 | Fruitloop *52* mutant | Non-toxic | ++ |
| 1 | -- | pExTra-Girr83 | Girr *83* replicate 1 | Non-toxic | +++ |
| 2 | -- | pExTra-Girr83 | Girr *83* replicate 2 | Non-toxic | +++ |
| 3 | -- | pExTra-Girr83 | Girr *83* replicate 3 | Non-toxic | +++ |

*Key: NG (no growth) - (no pink color) +(faint pink color) ++(obvious pink color) +++ (dark pink color)

#### Gene *84*; Score *0*

Images taken after 4 days at 37 ºC

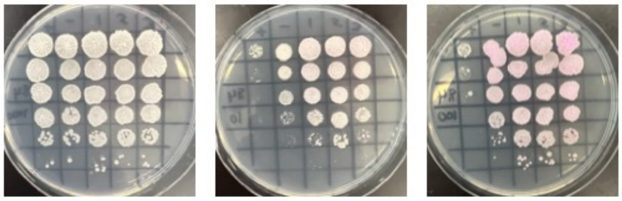


0 ng/ml aTc | 10 ng/ml aTc | 100 ng/ml aTc

| Lane | Gene ID | Plasmid name | Gene name | Toxic/Non-toxic | Colony color on 100 ng/ml aTc plate* |
|---|---|---|---|---|---|
| + Toxic control | -- | pExTra02 | Fruitloop *52* | Toxic | - |
| - Non-toxic control | -- | pExTra03 | Fruitloop *52* mutant | Non-toxic | ++ |
| 1 | -- | pExTra-Girr84 | Girr *84* replicate 1 | Non-toxic | ++ |
| 2 | -- | pExTra-Girr84 | Girr *84* replicate 2 | Non-toxic | ++ |
| 3 | -- | pExTra-Girr84 | Girr *84* replicate 3 | Non-toxic | ++ |

*Key: NG (no growth) - (no pink color) +(faint pink color) ++(obvious pink color) +++ (dark pink color)

#### Gene *85*; Score *0*

Images taken after 4 days at 37 ºC.

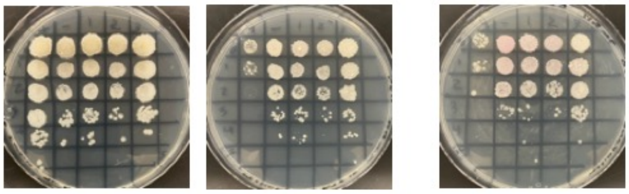


0 ng/ml aTc | 10 ng/ml aTc | 100 ng/ml aTc

| Lane | Gene ID | Plasmid name | Gene name | Toxic/Non-toxic | Colony color on 100 ng/ml aTc plate* |
|---|---|---|---|---|---|
| + Toxic control | -- | pExTra02 | Fruitloop *52* | Toxic | - |
| - Non-toxic control | -- | pExTra03 | Fruitloop *52* mutant | Non-toxic | ++ |
| 1 | -- | pExTra-Girr85 | Girr 85 replicate 1 | Non-toxic | + |
| +2 | -- | pExTra-Girr85 | Girr 85 replicate 2 | Non-toxic | + |
| 3 | -- | pExTra-Girr85 | Girr 85 replicate 3 | Non-toxic | + |

*Key: NG (no growth) - (no pink color) +(faint pink color) ++(obvious pink color) +++ (dark pink color)

#### Gene *86*; Score *0*

Images taken after 4 days at 37 ºC

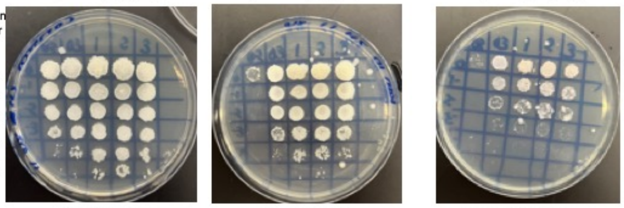


0 ng/ml aTc | 10 ng/ml aTc | 100 ng/ml aTc

| Lane | Gene ID | Plasmid name | Gene name | Toxic/Non-toxic | Colony color on 100 ng/ml aTc plate* |
|---|---|---|---|---|---|
| + Toxic control | -- | pExTra02 | Fruitloop *52* | Toxic | - |
| - Non-toxic control | -- | pExTra03 | Fruitloop *52* mutant | Non-toxic | - |
| 1 | -- | pExTra-Girr86 | Girr *86* replicate 1 | Non-toxic | - |
| 2 | -- | pExTra-Girr86 | Girr *86* replicate 2 | Non-toxic | - |
| 3 | -- | pExTra-Girr86 | Girr *86* replicate 3 | Non-toxic | - |

*Key: NG (no growth) - (no pink color) +(faint pink color) ++(obvious pink color) +++ (dark pink color)

#### Gene *87*; Score *0*

Images taken after 4 days at 37°C

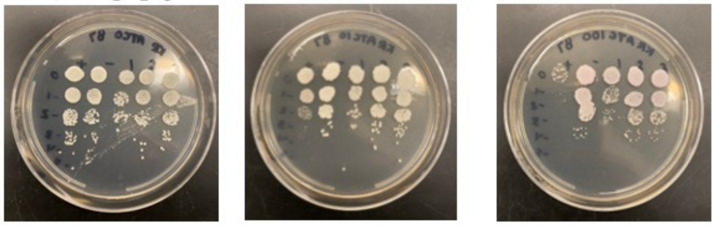


0 ng/ml aTc | 10 ng/ml aTc | 100 ng/ml aTc

| Lane | Gene ID | Plasmid name | Gene name | Toxic/Non-toxic | Colony color on 100 ng/ml aTc plate* |
|---|---|---|---|---|---|
| + Toxic control | -- | pExTra02 | Fruitloop *52* | Toxic | - |
| - Non-toxic control | -- | pExTra03 | Fruitloop *52 mutant* | Non-toxic | + |
| 1 | -- | pExTra-Gir87 | Girr *87* replicate 1 | Non-toxic | - |
| 2 | -- | pExTra-Girr87 | Girr *87* replicate 2 | Non-toxic | - |
| 3 | -- | pExTra-Girr87 | Girr *87* replicate 3 | Non-toxic | - |

*Key: NG (no growth) - (no pink color) + (faint pink color) ++ (obvious pink color) +++ (dark pink color)

#### Gene *88*; Score *0*

Images taken after 4 days at 37 ºC REPLICATE

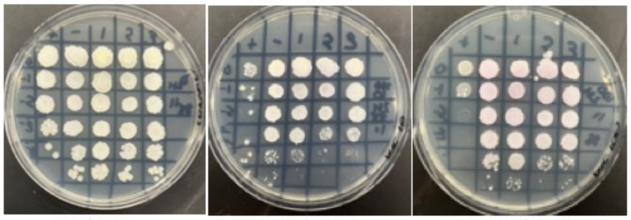


0 ng/ml aTc | 10 ng/ml aTc | 100 ng/ml aTc

| Lane | Gene ID | Plasmid name | Gene name | Toxic/Non-toxic | Colony color on 100 ng/ml aTc plate* |
|---|---|---|---|---|---|
| + Toxic control | | pExTra02 | Fruitloop *52* | Toxic | - |
| - Non-toxic control | | pExTra03 | Fruitloop *52* mutant | Non-toxic | + |
| 1 | | pExTra-Girr88 | Girr *88* replicate 1 | Non-toxic | + |
| 2 | | pExTra-Girr88 | Girr *88* replicate 2 | Non-toxic | + |
| 3 | | pExTra-Girr88 | Girr *88* replicate 3 | Non-toxic | + |

*Key: NG (no growth) - (no pink color) +(faint pink color) ++(obvious pink color) +++ (dark pink color)

#### Gene *89*; Score *0*

Images taken after 5 days at 37 ºC Replicate experiment

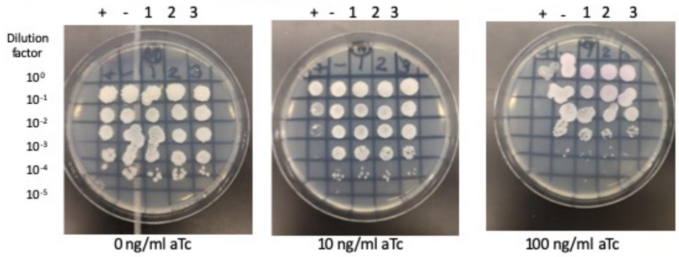


| Lane | Gene ID | Plasmid name | Gene name | Toxic/Non-toxic | Colony color on 100 ng/ml aTc plate* |
|---|---|---|---|---|---|
| + Toxic control | -- | pExTra02 | Fruitloop *52* | Toxic | - |
| - Non-toxic control | -- | pExTra03 | Fruitloop *52* mutant | Non-toxic | + |
| 1 | -- | pExTra-Girr89 | Girr 89 replicate 1 | Non-toxic | +++ |
| 2 | -- | pExTra-Girr89 | Girr 89 replicate 2 | Non-toxic | +++ |
| 3 | -- | pExTra-Girr89 | Girr 89 replicate 3 | Non-toxic | ++ |

*Key: NG (no growth) - (no pink color) +(faint pink color) ++(obvious pink color) +++ (dark pink color)

#### Gene *93*; Score *0*

Images taken after 4 days at 37 ºC

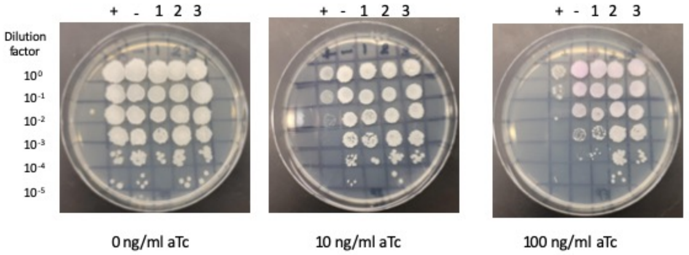


| Lane | Gene ID | Plasmid name | Gene name | Toxic/Non-toxic | Colony color on 100 ng/ml aTc plate* |
|---|---|---|---|---|---|
| + Toxic control | -- | pExTra02 | Fruitloop *52* | Toxic | - |
| - Non-toxic control | -- | pExTra03 | Fruitloop *52* mutant | Non-toxic | ++ |
| 1 | -- | pExTra-Girr93 | Girr 93 replicate 1 | Non-toxic | + |
| 2 | -- | pExTra-Girr93 | Girr 93 replicate 2 | Non-toxic | + |
| 3 | -- | pExTra-Girr93 | Girr 93 replicate 3 | Non-toxic | + |

*Key: NG (no growth) - (no pink color) +(faint pink color) ++(obvious pink color) +++ (dark pink color)

#### Gene *90*; Score *0*

Images taken after 5 days at 37 ºC

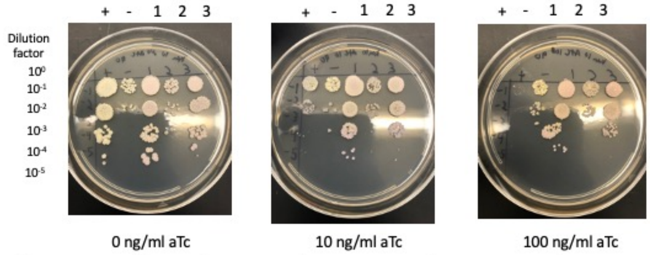


| Lane | Gene ID | Plasmid name | Gene name | Toxic/Non-toxic | Colony color on 100 ng/ml aTc plate* |
|---|---|---|---|---|---|
| + Toxic control | -- | pExTra02 | Fruitloop *52* | Toxic | - |
| - Non-toxic control | -- | pExTra03 | Fruitloop *52 mutant* | Non-toxic | + |
| 1 | -- | pExTra-Girr90 | Girr *90* replicate 1 | Non-toxic | + |
| 2 | -- | pExTra-Girr90 | Girr *90* replicate 2 | Non-toxic | - |
| 3 | -- | pExTra-Girr90 | Girr *90* replicate 3 | Non-toxic | + |

*Key: NG (no growth) - (no pink color) +(faint pink color) ++(obvious pink color) +++ (dark pink color)

#### Gene *95*; Score *0*

Images taken after 4 days at 37 ºC

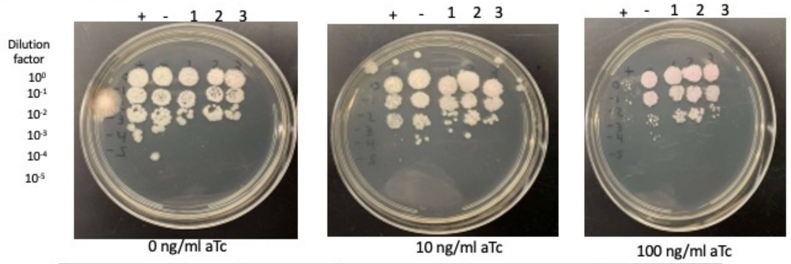


| Lane | Gene ID | Plasmid name | Gene name | Toxic/Non-toxic | Colony color on 100 ng/ml aTc plate* |
|---|---|---|---|---|---|
| + Toxic control | -- | pExTra02 | Fruitloop *52* | Toxic | - |
| - Non-toxic control | -- | pExTra03 | Fruitloop *52 mutant* | Non-toxic | + |
| 1 | | pExTra-Girr95 | Girr 95 replicate 1 | Non-toxic | + |
| 2 | | pExTra-Girr95 | Girr 95 replicate 2 | Non-toxic | + |
| 3 | | pExTra-Girr95 | Girr 95 replicate 3 | Non-toxic | + |

*Key: NG (no growth) - (no pink color) +(faint pink color) ++(obvious pink color) +++ (dark pink color)

#### Gene *91*; Score *0*

Images taken after 5 days at 37 ºC

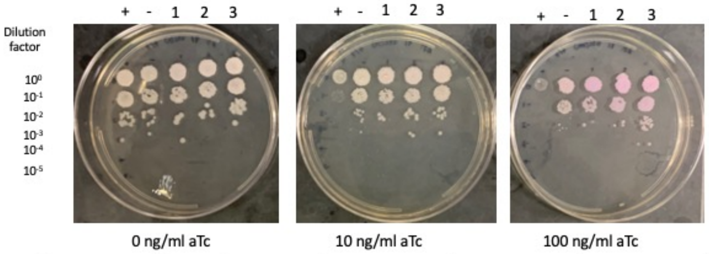


| Lane | Gene ID | Plasmid name | Gene name | Toxic/Non-toxic | Colony color on 100 ng/ml aTc plate* |
|---|---|---|---|---|---|
| + Toxic control | -- | pExTra02 | Fruitloop *52* | Toxic | - |
| - Non-toxic control | -- | pExTra03 | Fruitloop *52 mutant* | Non-toxic | + |
| 1 | -- | pExTra-Girr 91 | Girr 91 replicate 1 | Non-toxic | ++ |
| 2 | -- | pExTra-Girr 91 | Girr 91 replicate 2 | Non-toxic | ++ |
| 3 | -- | pExTra-Girr 91 | Girr 91 replicate 3 | Non-toxic | ++ |

*Key: NG (no growth) - (no pink color) +(faint pink color) ++(obvious pink color) +++ (dark pink color)

#### Gene *96*; Score *0*

Images taken after 5 days at 37 ºC and 3 days at 4°C.

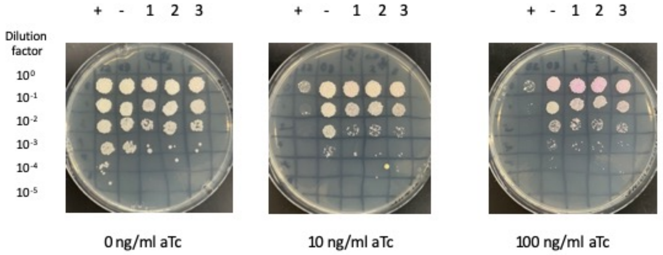


| Lane | Gene ID | Plasmid name | Gene name | Toxic/Non-toxic | Colony color on 100 ng/ml aTc plate* |
|---|---|---|---|---|---|
| + Toxic control | -- | pExTra02 | Fruitloop *52* | Toxic | - |
| - Non-toxic control | -- | pExTra03 | Fruitloop *52* mutant | Non-toxic | + |
| 1 | -- | pExTra-Girr96 | Girr *96* replicate 1 | Non-toxic | ++ |
| 2 | -- | pExTra-Girr96 | Girr *96* replicate 2 | Non-toxic | ++ |
| 3 | -- | pExTra-Girr96 | Girr *96* replicate 3 | Non-toxic | + |

*Key: NG (no growth) - (no pink color) +(faint pink color) ++(obvious pink color) +++ (dark pink color)

#### Gene *92*; Score *0*

Images taken after 4 days at 37 ºC REPLCATE EXPERIMENT 2023

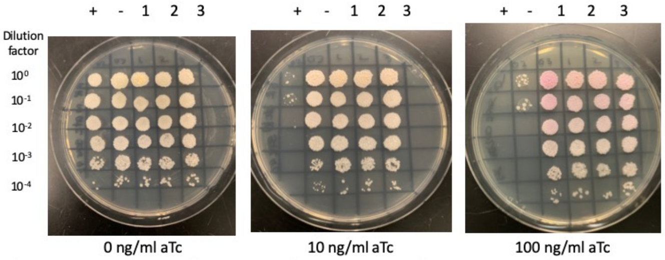


| Lane | Gene ID | Plasmid name | Gene name | Toxic/Non-toxic | Colony color on 100 ng/ml aTc plate* |
|---|---|---|---|---|---|
| + Toxic control | -- | pExTra02 | Fruitloop *52* | Toxic | - |
| - Non-toxic control | -- | pExTra03 | Fruitloop *52* mutant | Non-toxic | +++ |
| 1 | -- | pExTra-Girr92 | Girr *92* replicate 1 | Non-toxic | ++ |
| 2 | -- | pExTra-Girr92 | Girr *92* replicate 2 | Non-toxic | ++ |
| 3 | -- | pExTra-Girr92 | Girr *92* replicate 3 | Non-toxic | ++ |

*Key: NG (no growth) - (no pink color) +(faint pink color) ++(obvious pink color) +++ (dark pink color)

#### Gene *97*; Score *0*

Images taken after 5 days at 37 ºC REPLCATE EXPERIMENT

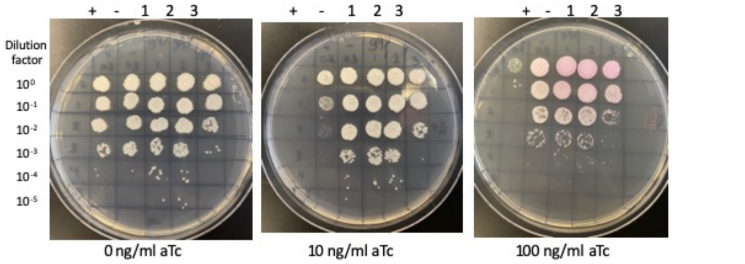


| Lane | Gene ID | Plasmid name | Gene name | Toxic/Non-toxic | Colony color on 100 ng/ml aTc plate* |
|---|---|---|---|---|---|
| + Toxic control | -- | pExTra02 | Fruitloop *52* | Toxic | - |
| - Non-toxic control | -- | pExTra03 | Fruitloop *52* mutant | Non-toxic | + |
| 1 | -- | pExTra-Girr97 | Girr *97* replicate 1 | Non-toxic | ++ |
| 2 | -- | pExTra-Girr97 | Girr *97* replicate 2 | Non-toxic | +++ |
| 3 | -- | pExTra-Girr97 | Girr *97* replicate 3 | Non-toxic | +++ |

*Key: NG (no growth) - (no pink color) +(faint pink color) ++(obvious pink color) +++ (dark pink color)

#### Gene *98*; Score *0*

Images taken after 4 days at 37 ºC

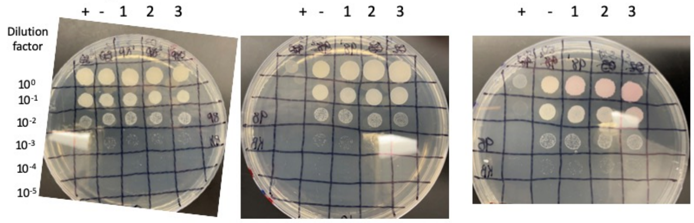


0 ng/ml aTc 10 ng/ml aTc 100 ng/ml aTc

| Lane | Gene ID | Plasmid name | Gene name | Toxic/Non-toxic | Colony color on 100 ng/ml aTc plate* |
|---|---|---|---|---|---|
| + Toxic control | -- | pExTra02 | Fruitloop *52* | Toxic | - |
| - Non-toxic control | -- | pExTra03 | Fruitloop *52* mutant | Non-toxic | + |
| 1 | -- | pExTra-Girr98 | Girr *98* replicate 1 | Non-Toxic | ++ |
| 2 | -- | pExTra-Girr98 | Girr *98* replicate 2 | Non-Toxic | ++ |
| 3 | -- | pExTra-Girr98 | Girr *98* replicate 3 | Non-Toxic | ++ |

*Key: NG (no growth) - (no pink color) +(faint pink color) ++(obvious pink color) +++ (dark pink color)

#### Gene *102*; Score *1*

Images taken after 4 days at 37 ºC SECOND EXPERIMENT

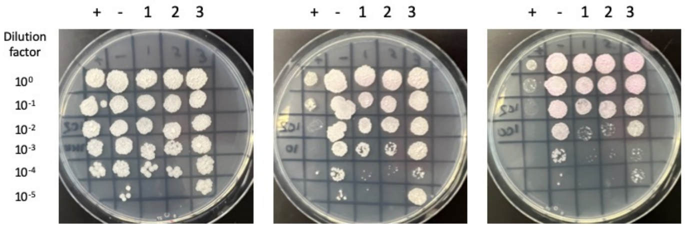


0 ng/ml aTc 10 ng/ml aTc 100 ng/ml aTc

| Lane | Gene ID | Plasmid name | Gene name | Toxic/Non-toxic | Colony color on 100 ng/ml aTc plate* |
|---|---|---|---|---|---|
| + Toxic control | -- | pExTra02 | Fruitloop *52* | Toxic | - |
| - Non-toxic control | -- | pExTra03 | Fruitloop *52* mutant | Non-toxic | + |
| 1 | -- | pExTra-Girr102 | Girr *102* replicate 1 | Toxic | + |
| 2 | -- | pExTra-Girr102 | Girr *102* replicate 2 | Toxic | + |
| 3 | -- | pExTra-Girr102 | Girr *102* replicate 3 | Toxic | + |

*Key: NG (no growth) - (no pink color) +(faint pink color) ++(obvious pink color) +++ (dark pink color)

#### Gene *99*; Score *0*

Images taken after 5 days at 37 ºC

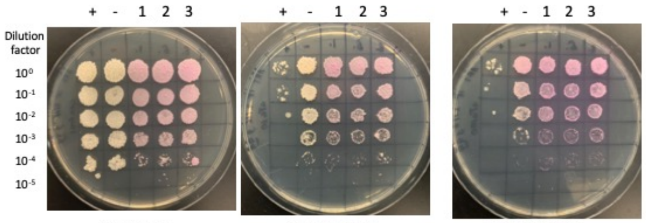


0 ng/ml aTc 10 ng/ml aTc 100 ng/ml aTc

| Lane | Gene ID | Plasmid name | Gene name | Toxic/Non-toxic | Colony color on 100 ng/ml aTc plate* |
|---|---|---|---|---|---|
| + Toxic control | ----- | pExTra02 | Fruitloop *52* | Toxic | - |
| - Non-toxic control | ------ | pExTra03 | Fruitloop *52* mutant | Non-toxic | ++ |
| 1 | ------- | pExTra-Girr99 | Girr *99* replicate 1 | Non-toxic | +++ |
| 2 | ------- | pExTra-Girr99 | Girr *99* replicate 2 | Non-toxic | +++ |
| 3 | --------- | pExTra-Girr99 | Girr *99* replicate 3 | Non-toxic | +++ |

*Key: NG (no growth) - (no pink color) +(faint pink color) ++(obvious pink color) +++ (dark pink color)

#### Gene *103*; Score 0

Images taken after 5 days at 37 ºC

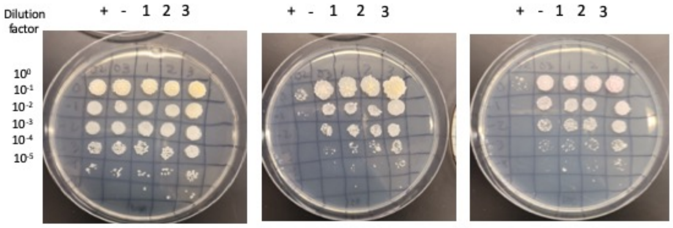


0 ng/ml aTc 10 ng/ml aTc 100 ng/ml aTc

| Lane | Gene ID | Plasmid name | Gene name | Toxic/Non-toxic | Colony color on 100 ng/ml aTc plate* |
|---|---|---|---|---|---|
| + Toxic control | -- | pExTra02 | Fruitloop *52* | Toxic | - |
| - Non-toxic control | -- | pExTra03 | Fruitloop *52* mutant | Non-toxic | + |
| 1 | -- | pExTra-Girr103 | Girr *103* replicate 1 | Non-toxic | + |
| 2 | -- | pExTra-Girr103 | Girr *103* replicate 2 | Non-toxic | + |
| 3 | -- | pExTra-Girr103 | Girr *103* replicate 3 | Non-toxic | ++ |

*Key: NG (no growth) - (no pink color) +(faint pink color) ++(obvious pink color) +++ (dark pink color)

#### Gene *100*; Score *2*

Images taken after 4 days at 37 ºC REPLICATE

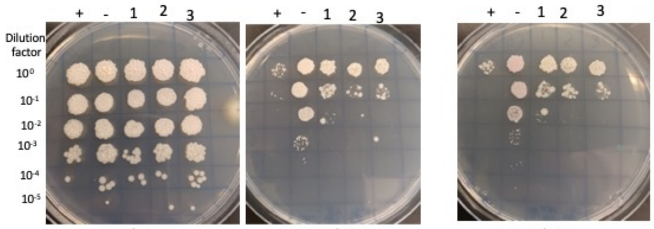


0 ng/ml aTc 10 ng/ml aTc 100 ng/ml aTc

| Lane | Gene ID | Plasmid name | Gene name | Toxic/Non-toxic | Colony color on 100 ng/ml aTc plate* |
|---|---|---|---|---|---|
| + Toxic control | -- | pExTra02 | Fruitloop *52* | Toxic | - |
| - Non-toxic control | -- | pExTra03 | Fruitloop *52* mutant | Non-toxic | + |
| 1 | -- | pExTra-Girr100 | Girr *100* replicate 1 | Toxic | - |
| 2 | -- | pExTra-Girr100 | Girr *100* replicate 2 | Toxic | - |
| 3 | -- | pExTra-Girr100 | Girr *100* replicate 3 | Toxic | - |

*Key: NG (no growth) - (no pink color) +(faint pink color) ++(obvious pink color) +++ (dark pink color)

#### Gene *101*; Score *0*

Images taken after 4 days at 37 ºC and 3 days at RT FIRST EXPERIMENT

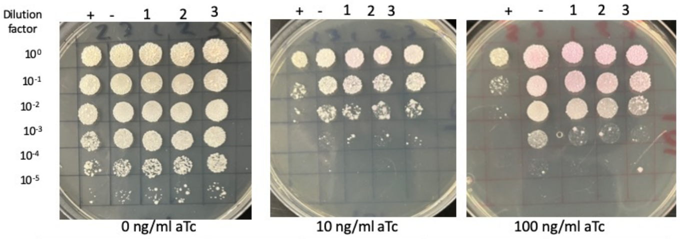


0 ng/ml aTc 10 ng/ml aTc 100 ng/ml aTc

| Lane | Gene ID | Plasmid name | Gene name | Toxic/Non-toxic | Colony color on 100 ng/ml aTc plate* |
|---|---|---|---|---|---|
| + Toxic control | -- | pExTra02 | Fruitloop *52* | Toxic | - |
| - Non-toxic control | -- | pExTra03 | Fruitloop *52* mutant | Non-toxic | ++ |
| 1 | -- | pExTra-Girr101 | Girr 101 replicate 1 | Non-toxic | ++ |
| 2 | -- | pExTra-Girr101 | Girr 101 replicate 2 | Non-toxic | ++ |
| 3 | -- | pExTra-Girr101 | Girr 101 replicate 3 | Non-toxic | ++ |

*Key: NG (no growth) - (no pink color) +(faint pink color) ++(obvious pink color) +++ (dark pink color)

**Supplemental Table 1: DNA oligos used in this study**

| **Oligo Name** | Oligo Sequence (5' → 3') |
|---|---|
| **oGirr1_R** | TGC AGG ATC CGA CTC GAG TGT CGA CTC AGG TCA CAA GCT TCA AAC |
| **oGirr1_F** | ATG CGG AGG AAT CAC TTC CAT ATG CCA CCT GTA CCT AAA G |
| **oGirr2_R** | TGC AGG ATC CGA CTC GAG TGT CGA CTC AGT AGA TCC GTC TAG GC |
| **oGirr2_F** | ATG CGG AGG AAT CAC TTC CAT ATG GCT GTT TTG CAG GTC |
| **oGirr3_R** | TGC AGG ATC CGA CTC GAG TGT CGA CTC AGC GTG ATC CAT CTT CC |
| **oGirr3_F** | ATG CGG AGG AAT CAC TTC CAT ATG ACT GCT TCA ACG CC |
| **oGirr4_R** | TGC AGG ATC CGA CTC GAG TGT CGA CTC AGC GCT GGA TGC TTC TG |
| **oGirr4_F** | ATG CGG AGG AAT CAC TTC CAT ATG GAT CAC GCT GAG TAT GC |
| **oGirr5_R** | TGC AGG ATC CGA CTC GAG TGT CGA CTC AGT GGA GTT CTC CAC G |
| **oGirr5_F** | ATG CGG AGG AAT CAC TTC CAT ATG TCT GAT GAT GTG ACA GC |
| **oGirr6_R** | TGC AGG ATC CGA CTC GAG TGT CGA CTC AGC TGC CCG TCT TAT TG |
| **oGirr6_F** | ATG CGG AGG AAT CAC TTC CAT ATG GCT TTC AAC AAC TTC ATT CC |
| **oGirr7_R** | TGC AGG ATC CGA CTC GAG TGT CGA CTC ATA GCC TGT GAA TCG TGA TCG |
| **oGirr7_F** | ATG CGG AGG AAT CAC TTC CAT ATG CTT GCT ACC GCC G |
| **oGirr8_R** | TGC AGG ATC CGA CTC GAG TGT CGA CTC ACA CCT TCC GCA GC |
| **oGirr8_F** | ATG CGG AGG AAT CAC TTC CAT ATG ACG TTC CCT ACT CCG |
| **oGirr9_R** | TGC AGG ATC CGA CTC GAG TGT CGA CTC AGT CGC CAT ACG CG |
| **oGirr9_F** | ATG CGG AGG AAT CAC TTC CAT ATG GCG AAC GGT CCA AC |
| **oGirr10_R** | TGC AGG ATC CGA CTC GAG TGT CGA CTC AGA TGT ACT GAA CAC CGA TC |
| **oGirr10_F** | ATG CGG AGG AAT CAC TTC CAT ATG GCG ACT GAT TCA GCG |
| **oGirr11_R** | TGC AGG ATC CGA CTC GAG TGT CGA CTC AGC TGC CGT CCG AGT AC |
| **oGirr11_F** | ATG CGG AGG AAT CAC TTC CAT ATG ACG CAG CCA TTG ACC |
| **oGirr13_R** | TGC AGG ATC CGA CTC GAG TGT CGA CTC ACC AGC CGA ACA GAT C |
| **oGirr13_F** | ATG CGG AGG AAT CAC TTC CAT ATG TCT GTG AAG AAA CCC GAG |
| **oGirr12_R** | TGC AGG ATC CGA CTC GAG TGT CGA CTC ACT CGC TAT CTG CCT C |
| **oGirr12_F** | ATG CGG AGG AAT CAC TTC CAT ATG TCT GTG AAG AAA CCC GAG |
| **oGirr14_R** | TGC AGG ATC CGA CTC GAG TGT CGA CTC ACC CTC CCG GCA TGA C |
| **oGirr14_F** | ATG CGG AGG AAT CAC TTC CAT ATG CCG ATC TAC GTG GAC |
| **oGirr15_R** | TGC AGG ATC CGA CTC GAG TGT CGA CTC ACA TTG GGT AGC GGC |
| **oGirr15_F** | ATG CGG AGG AAT CAC TTC CAT ATG GCT AAG AAG CAT TAC CCC |
| **oGirr16_R** | TGC AGG ATC CGA CTC GAG TGT CGA CTC ATC CCT GCG GTG ACA G |
| **oGirr16_F** | ATG CGG AGG AAT CAC TTC CAT ATG TCG AAG TTT GAA CGC GAA TC |
| **oGirr17_R** | TGC AGG ATC CGA CTC GAG TGT CGA CTC ATG ACA GCG GAA GAA C |
| **oGirr17_F** | ATG CGG AGG AAT CAC TTC CAT ATG TCG TGG CCT TTG AAC |
| **oGirr18_R** | TGC AGG ATC CGA CTC GAG TGT CGA CTC ATA CGT CAG TAG CTC CAG AC |
| **oGirr18_F** | ATG CGG AGG AAT CAC TTC CAT ATG ACG TCT TCG TTT GAT CCG |
| **oGirr19_R** | TGC AGG ATC CGA CTC GAG TGT CGA CTC ATC CCC CAA TCT GC |

| | |
|---|---|
| **oGirr19_F** | ATG CGG AGG AAT CAC TTC CAT ATG ACA ATC AAA GGG TAT TTC G |
| **oGirr20_R** | TGC AGG ATC CGA CTC GAG TGT CGA CTC ATA CGG GAT TGA GCG GG |
| **oGirr20_F** | ATG CGG AGG AAT CAC TTC CAT ATG GGC TGG CAT GGC G |
| **oGirr21_R** | TGC AGG ATC CGA CTC GAG TGT CGA CTC AGG CGA CTA TGC GG |
| **oGirr21_F** | ATG CGG AGG AAT CAC TTC CAT ATG GGC ATT CCC AAC GC |
| **oGirr22_R** | TGC AGG ATC CGA CTC GAG TGT CGA CTC AGG GTT GCG CCC GCA AAC |
| **oGirr22_F** | ATG CGG AGG AAT CAC TTC CAT ATG AAC ATC AAA ACT GAT CAT CAG |
| **oGirr23_R** | TGC AGG ATC CGA CTC GAG TGT CGA CTC ACG GCA CCA CCA C |
| **oGirr23_F** | ATG CGG AGG AAT CAC TTC CAT ATG GCT TAT TCG AAG CAG TCG |
| **oGirr24_R** | TGC AGG ATC CGA CTC GAG TGT CGA CTC ATT CCC ACT CGA TCA G |
| **oGirr24_F** | ATG CGG AGG AAT CAC TTC CAT ATG AAA GTT TGG AAC GGC |
| **oGirr25_R** | TGC AGG ATC CGA CTC GAG TGT CGA CTC ACT CGC CGC GAA TGA TTT G |
| **oGirr25_F** | ATG CGG AGG AAT CAC TTC CAT ATG ACC CTC GCT GAT CGA C |
| **oGirr26_R** | TGC AGG ATC CGA CTC GAG TGT CGA CTC ACC AGG GAT CCG TG |
| **oGirr26_F** | ATG CGG AGG AAT CAC TTC CAT ATG CTA CGC AAC ACC ATC |
| **oGirr27_R** | TGC AGG ATC CGA CTC GAG TGT CGA CTC ATG ACA GTC TCC TGA CC |
| **oGirr27_F** | ATG CGG AGG AAT CAC TTC CAT ATG AAC AAG ATC CAC ATC GCC |
| **oGirr28_R** | TGC AGG ATC CGA CTC GAG TGT CGA CTC AGA TGT AGA AGA GGG TGT CG |
| **oGirr28_F** | ATG CGG AGG AAT CAC TTC CAT ATG ACC AAA CGA GGG GC |
| **oGirr29_R** | TGC AGG ATC CGA CTC GAG TGT CGA CTC ATG GCT TGC CGC C |
| **oGirr29_F** | ATG CGG AGG AAT CAC TTC CAT ATG GAC CGT CTC GGA ATC ATC |
| **oGirr30_R** | TGC AGG ATC CGA CTC GAG TGT CGA CTC ATG CCG CCC TCT CG |
| **oGirr30_F** | ATG CGG AGG AAT CAC TTC CAT ATG AAG GTC ACC TAC CGC G |
| **oGirr31_R** | TGC AGG ATC CGA CTC GAG TGT CGA CTC ATT TGT CGC AGG CGA C |
| **oGirr31_F** | ATG CGG AGG AAT CAC TTC CAT ATG AGC TTC ACC TGG TTC C |
| **oGirr32_R** | TGC AGG ATC CGA CTC GAG TGT CGA CTC ATA CGC ACG ATG CGC C |
| **oGirr32_F** | ATG CGG AGG AAT CAC TTC CAT ATG GCG CGC CAA CGC ATC |
| **oGirr33_R** | TGC AGG ATC CGA CTC GAG TGT CGA CTC AGG TCG CGA CGC C |
| **oGirr33_F** | ATG CGG AGG AAT CAC TTC CAT ATG CGT ATA GCG AAT GCA TAT GTG G |
| **oGirr34_R** | TGC AGG ATC CGA CTC GAG TGT CGA CTC ATG CGG CAG TGA CC |
| **oGirr34_F** | ATG CGG AGG AAT CAC TTC CAT ATG CAA GCC ATG TTG ACA CG |
| **oGirr35_R** | TGC AGG ATC CGA CTC GAG TGT CGA CTC ACC GGT CGC GTC G |
| **oGirr35_F** | ATG CGG AGG AAT CAC TTC CAT ATG ATC TGG GAA TCG GTG CG |
| **oGirr36_R** | TGC AGG ATC CGA CTC GAG TGT CGA CTC AGT CAG CGA CAT GAT GG |
| **oGirr36_F** | ATG CGG AGG AAT CAC TTC CAT ATG AAA CAC CAG GAA AGG G |
| **oGirr37_R** | TGC AGG ATC CGA CTC GAG TGT CGA CTC AAA CCC CTC TCA CAG C |
| **oGirr37_F** | ATG CGG AGG AAT CAC TTC CAT ATG TCG CTG ACT AGC GAC C |
| **oGirr38_R** | TGC AGG ATC CGA CTC GAG TGT CGA CTC ACT CAT CAT CCG ATT CG |
| **oGirr38_F** | ATG CGG AGG AAT CAC TTC CAT ATG CAC GTT TCT GGA CC |
| **oGirr39_R** | TGC AGG ATC CGA CTC GAG TGT CGA CTC ACG AGG CGT TGA TTA AGC |
| **oGirr39_F** | ATG CGG AGG AAT CAC TTC CAT ATG AGT GAT CCG CAG TTG G |

| | |
|---|---|
| **oGirr40_R** | TGC AGG ATC CGA CTC GAG TGT CGA CTC ACT CAG CAT GGC TAT ATC G |
| **oGirr40_F** | ATG CGG AGG AAT CAC TTC CAT ATG CGA GAT GTG CAA CTG |
| **oGirr41_R** | TGC AGG ATC CGA CTC GAG TGT CGA CTC AGT TGC ACA TCT CGC ATC |
| **oGirr41_F** | ATG CGG AGG AAT CAC TTC CAT ATG TGT GGT GGT TGT GAG G |
| **oGirr42_R** | TGC AGG ATC CGA CTC GAG TGT CGA CTC AGG ACA GCT CTC CC |
| **oGirr42_F** | ATG CGG AGG AAT CAC TTC CAT ATG ACT GCA GCT ACT GAC C |
| **oGirr43_R** | TGC AGG ATC CGA CTC GAG TGT CGA CTC ATA GGG GAG CAT GAG G |
| **oGirr43_F** | ATG CGG AGG AAT CAC TTC CAT ATG TCA GCC TTC TTT CAG C |
| **oGirr44_R** | TGC AGG ATC CGA CTC GAG TGT CGA CTC ATT TTG TTT CTA ACG CAA ACG |
| **oGirr44_F** | ATG CGG AGG AAT CAC TTC CAT ATG GCA ACT AAG AAA CGC AG |
| **oGirr45_R** | TGC AGG ATC CGA CTC GAG TGT CGA CTC AAT CCG TCA TGG TCC AAG |
| **oGirr45_F** | ATG CGG AGG AAT CAC TTC CAT ATG ACC ACC AAT GAT CGC G |
| **oGirr46_R** | TGC AGG ATC CGA CTC GAG TGT CGA CTC AGT CCT TGC GGC GTT G |
| **oGirr46_F** | ATG CGG AGG AAT CAC TTC CAT ATG AAC GAG AAC AAG GAA CAC C |
| **oGirr47_R** | TGC AGG ATC CGA CTC GAG TGT CGA CTC ATG CGG CGG GGT TTT TC |
| **oGirr47_F** | ATG CGG AGG AAT CAC TTC CAT ATG GCG CGA ACT GCG C |
| **oGirr48_R** | TGC AGG ATC CGA CTC GAG TGT CGA CTC ATG ACA CGG CCG C |
| **oGirr48_F** | ATG CGG AGG AAT CAC TTC CAT ATG TCT GAA CTA CAG CGT ATC AAC C |
| **oGirr49_R** | TGC AGG ATC CGA CTC GAG TGT CGA CTC ATC GCG TCT CCC TC |
| **oGirr49_F** | ATG CGG AGG AAT CAC TTC CAT ATG AGC TTC TCT TTC TAT GCA G |
| **oGirr50_R** | TGC AGG ATC CGA CTC GAG TGT CGA CTC ACG CGG ACC TCG |
| **oGirr50_F** | ATG CGG AGG AAT CAC TTC CAT ATG AGC ACT CCC AGA TGG |
| **oGirr51_R** | TGC AGG ATC CGA CTC GAG TGT CGA CTC ACG CTG TCT CCC C |
| **oGirr51_F** | ATG CGG AGG AAT CAC TTC CAT ATG AGT ACG TCT GCT CCT AAG |
| **oGirr52_R** | TGC AGG ATC CGA CTC GAG TGT CGA CTC ATC TCG GGG TGA TGC |
| **oGirr52_F** | ATG CGG AGG AAT CAC TTC CAT ATG AAT CTT GTT GAG CGT TTG AAC |
| **oGirr53_R** | TGC AGG ATC CGA CTC GAG TGT CGA CTC ATG CTG CTT CTC CC |
| **oGirr53_F** | ATG CGG AGG AAT CAC TTC CAT ATG CTA GAT CGA GAT CCT AAA CC |
| **oGirr54_R** | TGC AGG ATC CGA CTC GAG TGT CGA CTC ACA CCC ACC CCG TG |
| **oGirr54_F** | ATG CGG AGG AAT CAC TTC CAT ATG AGG CGC AAC GAG AAG TC |
| **oGirr55_R** | TGC AGG ATC CGA CTC GAG TGT CGA CTC ACG CGC CCC ACC TC |
| **oGirr55_F** | ATG CGG AGG AAT CAC TTC CAT ATG CCG AAT TCC CCG TTC ATC |
| **oGirr56_R** | TGC AGG ATC CGA CTC GAG TGT CGA CTC ATG CAA CGG ACT CC |
| **oGirr56_F** | ATG CGG AGG AAT CAC TTC CAT ATG AGC ATC GAT TGG TTC G |
| **oGirr57_R** | TGC AGG ATC CGA CTC GAG TGT CGA CTC ATG CAA CGG ACT CC |
| **oGirr57_F** | ATG CGG AGG AAT CAC TTC CAT ATG ACC GAC CTG TCT C |
| **oGirr58_R** | TGC AGG ATC CGA CTC GAG TGT CGA CTC ACG AAG CCT CCA ACC |
| **oGirr58_F** | ATG CGG AGG AAT CAC TTC CAT ATG AGC AAC GGG AAC AGG |
| **oGirr59_R** | TGC AGG ATC CGA CTC GAG TGT CGA CTC AGT TTT CGT CCT TAT CTC G |
| **oGirr59_F** | ATG CGG AGG AAT CAC TTC CAT ATG TGC GTG TGC GGC |
| **oGirr60_R** | TGC AGG ATC CGA CTC GAG TGT CGA CTC ATG CTG TCC ACC TG |

| | |
|---|---|
| **oGirr60_F** | ATG CGG AGG AAT CAC TTC CAT ATG GTT GTT GAT ACA CGG G |
| **oGirr61_R** | TGC AGG ATC CGA CTC GAG TGT CGA CTC ATT CGG TCA CCT CCG |
| **oGirr61_F** | ATG CGG AGG AAT CAC TTC CAT ATG GAC AGC ATG AGC AAC |
| **oGirr62_R** | TGC AGG ATC CGA CTC GAG TGT CGA CTC ATG CGG GGG CGC C |
| **oGirr62_F** | ATG CGG AGG AAT CAC TTC CAT ATG AGC GAC GTG GAC GTT G |
| **oGirr63_R** | TGC AGG ATC CGA CTC GAG TGT CGA CTC ACT TCG CAG CCT C |
| **oGirr63_F** | ATG CGG AGG AAT CAC TTC CAT ATG AGT CGC CGG TTT AC |
| **oGirr64_R** | TGC AGG ATC CGA CTC GAG TGT CGA CTC ATG ACG CGT CCT CTA G |
| **oGirr64_F** | ATG CGG AGG AAT CAC TTC CAT ATG ACC GGC CAC GTG TC |
| **oGirr65_R** | TGC AGG ATC CGA CTC GAG TGT CGA CTC ACC TGA ACA GCC CCT TC |
| **oGirr65_F** | ATG CGG AGG AAT CAC TTC CAT ATG GTT CCG TGC CCG C |
| **oGirr66_R** | TGC AGG ATC CGA CTC GAG TGT CGA CTC ATG GGA CCT GCC AG |
| **oGirr66_F** | ATG CGG AGG AAT CAC TTC CAT ATG TAC ACG GTT TCT GGG |
| **oGirr67_R** | TGC AGG ATC CGA CTC GAG TGT CGA CTC ATG CGG TGA CTT CTT C |
| **oGirr67_F** | ATG CGG AGG AAT CAC TTC CAT ATG ATC ACC GTT GCT TGC |
| **oGirr68_R** | TGC AGG ATC CGA CTC GAG TGT CGA CTC ACG CGG CCC CCC C |
| **oGirr68_F** | ATG CGG AGG AAT CAC TTC CAT ATG AAA CAC ATC GTG ATG TTC TCC G |
| **oGirr69_R** | TGC AGG ATC CGA CTC GAG TGT CGA CTC ACG TGT CCT CCT CTT C |
| **oGirr69_F** | ATG CGG AGG AAT CAC TTC CAT ATG ACG TGC CTG TTG TG |
| **oGirr70_R** | TGC AGG ATC CGA CTC GAG TGT CGA CTC ACT GTC CAA TGG CCT TTC |
| **oGirr70_F** | ATG CGG AGG AAT CAC TTC CAT ATG AGG ATC AGG TCA ACG AAA C |
| **oGirr71_R** | TGC AGG ATC CGA CTC GAG TGT CGA CTC AGG CAC GAT CAA TCG |
| **oGirr71_F** | ATG CGG AGG AAT CAC TTC CAT ATG AAT TAC CGG CAG ATC G |
| **oGirr72_R** | TGC AGG ATC CGA CTC GAG TGT CGA CTC ACG ACA TCC GCC C |
| **oGirr72_F** | ATG CGG AGG AAT CAC TTC CAT ATG CCT GAC CGG TAC G |
| **oGirr73_R** | TGC AGG ATC CGA CTC GAG TGT CGA CTC ATC GAC GGG CCA C |
| **oGirr73_F** | ATG CGG AGG AAT CAC TTC CAT ATG AGC CAC ACG CTG AC |
| **oGirr74_R** | TGC AGG ATC CGA CTC GAG TGT CGA CTC ATT CGT CGC CTT TCG |
| **oGirr74_F** | ATG CGG AGG AAT CAC TTC CAT ATG ACG ATG TTT GTG TCG TC |
| **oGirr75_R** | TGC AGG ATC CGA CTC GAG TGT CGA CTC AGC CAC GGT CTT C |
| **oGirr75_F** | ATG CGG AGG AAT CAC TTC CAT ATG AAC AAC CCC GAG TTG |
| **oGirr76_R** | TGC AGG ATC CGA CTC GAG TGT CGA CTC ACT TGT CTT CCC CCT C |
| **oGirr76_F** | ATG CGG AGG AAT CAC TTC CAT ATG GCT GAT CTC GGA GTG |
| **oGirr77_R** | TGC AGG ATC CGA CTC GAG TGT CGA CTC ATG CTT CCT CCC CTG TAG |
| **oGirr77_F** | ATG CGG AGG AAT CAC TTC CAT ATG AGC GGC GAC ATC AAC |
| **oGirr78_R** | TGC AGG ATC CGA CTC GAG TGT CGA CTC ACG CTT CCT CCC C |
| **oGirr78_F** | ATG CGG AGG AAT CAC TTC CAT ATG AGC GAC CCG GTA AC |
| **oGirr79_R** | TGC AGG ATC CGA CTC GAG TGT CGA CTC ATC GGC CAC ATG C |
| **oGirr79_F** | ATG CGG AGG AAT CAC TTC CAT ATG AGT AGC GAA GCC CAG |
| **oGirr80_R** | TGC AGG ATC CGA CTC GAG TGT CGA CTC ACA GTT CCT GCT CC |
| **oGirr80_F** | ATG CGG AGG AAT CAC TTC CAT ATG TTT CCG ATT ACC GAC AC |

| | |
|---|---|
| **oGirr81_R** | TGC AGG ATC CGA CTC GAG TGT CGA CTC ATC GGG TCT CTC CTC G |
| **oGirr81_F** | ATG CGG AGG AAT CAC TTC CAT ATG AGC AGC GAA GCC CAA AAC |
| **oGirr82_R** | TGC AGG ATC CGA CTC GAG TGT CGA CTC AGG AAC CCA CTT TCG C |
| **oGirr82_F** | ATG CGG AGG AAT CAC TTC CAT ATG ACC TTG AGC GAT GCA ATA G |
| **oGirr83_R** | TGC AGG ATC CGA CTC GAG TGT CGA CTC ACC GCG GAA CCT CC |
| **oGirr83_F** | ATG CGG AGG AAT CAC TTC CAT ATG CGC CGA TTG CGC TG |
| **oGirr84_R** | TGC AGG ATC CGA CTC GAG TGT CGA CTC AGC AGG ACG ATC CTT TC |
| **oGirr84_F** | ATG CGG AGG AAT CAC TTC CAT ATG ATT CAG GTT CAT TGC AGG |
| **oGirr85_R** | TGC AGG ATC CGA CTC GAG TGT CGA CTC ATG GTT GGG CCT C |
| **oGirr85_F** | ATG CGG AGG AAT CAC TTC CAT ATG GCT CAC GAA AGG ATC |
| **oGirr86_R** | TGC AGG ATC CGA CTC GAG TGT CGA CTC AGA ACG GCG GAG CCC |
| **oGirr86_F** | ATG CGG AGG AAT CAC TTC CAT ATG AGT ACC CCT GAG CGT G |
| **oGirr87_R** | TGC AGG ATC CGA CTC GAG TGT CGA CTC ACG GTT GGT CCT TTT C |
| **oGirr87_F** | ATG CGG AGG AAT CAC TTC CAT ATG CCG AAA CCA CCT G |
| **oGirr88_R** | TGC AGG ATC CGA CTC GAG TGT CGA CTC ATT TGT GGT GTC CTT TGC |
| **oGirr88_F** | ATG CGG AGG AAT CAC TTC CAT ATG ACC TTG TCC GTG ATT C |
| **oGirr89_R** | TGC AGG ATC CGA CTC GAG TGT CGA CTC AGA CAT CAG TGA TCC C |
| **oGirr89_F** | ATG CGG AGG AAT CAC TTC CAT ATG AGC GTC TAC GCA C |
| **oGirr90_R** | TGC AGG ATC CGA CTC GAG TGT CGA CTC ACT TGT CGA ACG CC |
| **oGirr90_F** | ATG CGG AGG AAT CAC TTC CAT ATG TCT GAT GCT CGT GTG |
| **oGirr91_F** | ATG CGG AGG AAT CAC TTC CAT ATG AGC ATG GAC TTC CAC |
| **oGirr91_R** | TGC AGG ATC CGA CTC GAG TGT CGA CTC ATT CTG CTA ATT TTA CCT GTT C |
| **oGirr92_R** | TGC AGG ATC CGA CTC GAG TGT CGA CTC ATG CAA TCC GCA GC |
| **oGirr92_F** | ATG CGG AGG AAT CAC TTC CAT ATG AGC CGG GTC TTT C |
| **oGirr93_R** | TGC AGG ATC CGA CTC GAG TGT CGA CTC AGA TTC GGC GGG G |
| **oGirr93_F** | ATG CGG AGG AAT CAC TTC CAT ATG CCC ACC TTC GCC |
| **oGirr95_R** | TGC AGG ATC CGA CTC GAG TGT CGA CTC ATT CCT GCC TGC C |
| **oGirr95_F** | ATG CGG AGG AAT CAC TTC CAT ATG CCG TTG CTC CAC |
| **oGirr96_R** | TGC AGG ATC CGA CTC GAG TGT CGA CTC ATC CTC GTA TGC ACC ACC |
| **oGirr96_F** | ATG CGG AGG AAT CAC TTC CAT ATG AAA CGC CGC GCG |
| **oGirr97_F** | ATG CGG AGG AAT CAC TTC CAT ATG AAC CTC ACA GAA TTT CTC AC |
| **oGirr97_R** | TGC AGG ATC CGA CTC GAG TGT CGA CTC ACT GTT CAG CTG CTT TC |
| **oGIRR_gp46_ATG34_For** | ATG CGG AGG AAT CAC TTC CAT ATG ACA ACG CCA CCA GGC AG |
| **oGIRR_gp46_TGA74_Rev** | TGC AGG ATC CGA CTC GAG TGT CGA CTC ACA CGG TGG ATG CGG T |
| **oGirr_2i_SEQ** | GCAGAACATGCCGAAGCGG |
| **oGirr_14ia_SEQ** | GCGAAGGCTGCTACTCAG |
| **oGirr_14ib_SEQ** | TCGCAGCGGCGTTCATTGC |
| **oGirr_14ic_SEQ** | CTGGGTCGGGTTATAGG |
| **oGirr_14id_SEQ** | GCAGCGGGGCAGGATTCG |
| **oGirr_15i_SEQ** | CCGCTGTACTACGAAGG |
| **oGirr_16i_SEQ** | TTGTCGGTGCAGGCTTTACC |

| | |
|---|---|
| **oGirr_18ia_SEQ** | CACGTTGAAGGCACTTCGG |
| **oGirr_18ib_SEQ** | GGCCTTCCGGGTATGG |
| **oGirr_18ic_SEQ** | ATCGATGTAGTTCACCCAG |
| **oGirr_19ia_SEQ** | CCGACGTGGCCGTGGACGG |
| **oGirr_19ib_SEQ** | CGGCACCGGCCAGTTTTGG |
| **oGirr_64ia_SEQ** | GAATAACCTAACCCTCTACC |
| **oGirr_64ib_SEQ** | GAGACCGAGCGGGAAGTGG |
| **oGirr_64ic_SEQ** | GTGCGCGTCCACCTGCC |
| **pExTra_seqF_SEQ** | GTACCCGTGTGTACGACCAGC |
| **pExTra_universalR** | CCCTTCGAGACCATAGATCTGTTCC |
| **pExTra_F** | GTCGACACTCGAGTCGGATCCTG |
| **pExTra_R** | ATGGAAGTGATTCCTCCGCATGC |
